# Pedigree-based estimation of human mobile element retrotransposition rates

**DOI:** 10.1101/506691

**Authors:** Julie Feusier, W. Scott Watkins, Jainy Thomas, Andrew Farrell, David J. Witherspoon, Lisa Baird, Hongseok Ha, Jinchuan Xing, Lynn B. Jorde

## Abstract

Germline mutation rates in humans have been estimated for a variety of mutation types, including single nucleotide and large structural variants. Here we directly measure the germline retrotransposition rate for the three active retrotransposon elements: L1, Alu, and SVA. We utilized three tools for calling Mobile Element Insertions (MEIs) (MELT, RUFUS, and TranSurVeyor) on blood-derived whole genome sequence (WGS) data from 603 CEPH individuals, comprising 33 three-generation pedigrees. We identified 27 *de novo* MEIs in 440 births. The retrotransposition rate estimates for Alu elements, one in 40, is roughly half the rate estimated using phylogenetic analyses, a difference in magnitude similar to that observed for single nucleotide variants. The L1 retrotransposition rate is one in 62 births and is within range of previous estimates (1:20-1:200 births). The SVA retrotransposition rate, one in 55 births, is much higher than the previous estimate of one in 900 births. Our large, three-generation pedigrees allowed us to assess parent-of-origin effects and the timing of insertion events in either gametogenesis or early embryonic development. We find a statistically significant paternal bias in Alu retrotransposition. Our study represents the first in-depth analysis of the rate and dynamics of human retrotransposition from WGS data in three-generation human pedigrees.

## Introduction

Non-LTR retrotransposons have played a large role in shaping the human genome by creating structural variation and influencing gene expression (Elbarbary et al. 2016; Bourque et al. 2018). In addition, there are at least 130 documented instances of retrotransposition events associated with human disease (Hancks and Kazazian 2016; Kazazian and Moran 2017). These retrotransposons mobilize via a “copy and paste” mechanism using an mRNA intermediate that is reverse-transcribed into the genome. There are three currently active non-LTR retrotransposons in humans: the autonomous long interspersed element 1 (L1), and two non-autonomous elements, the Alu short interspersed elements (SINE), and the composite element SINE-R-VNTR-Alu (SVA). These three retrotransposon families alone account for >25% of the human genome, and younger copies are polymorphic for their presence or absence in humans (Cordaux and Batzer 2009). There are more than 1.5 million non-LTR retrotransposons in the human genome (Cordaux and Batzer 2009), and a small fraction of them are active and still capable of creating new mobile element insertions (MEIs) in somatic and germline tissue.

Alu, L1, and SVA germline retrotransposition rates have been estimated through phylogenetic and disease-based studies. It is estimated that one *de novo* Alu insertion occurs in every ~20 births and a *de novo* L1 insertion event occurs once in everŷ150 live human births (Ewing and Kazazian 2010; Huang et al. 2010; Xing et al. 2009; Kazazian 1999; Li et al. 2001; Hormozdiari et al. 2011; Cordaux et al. 2006; Deininger and Batzer 1999; Hancks and Kazazian 2012). There are only a few thousand SVA elements in the human genome, and the current estimate for the rate of new SVA insertion events is one in every ~900 live human births (Xing et al. 2009). Although previous studies have identified *de novo* Alu, L1, and SVA insertions in large cohorts (Werling et al. 2018; Gardner et al. 2018), there has not yet been a rigorous empirical study of heritable germline retrotransposition and retrotranspositional timing in multi-generation pedigrees. Moreover, it is unknown whether human germline retrotransposition is affected by the parent’s age or sex, or whether retrotransposition rates differ among pedigrees.

To characterize *de novo* MEIs in a non-disease cohort, we analyzed WGS data from blood-derived DNA obtained in the 1980s and 1990s from 603 Utah Centre d’Etude du Polymorphisme Humain (CEPH) pedigree members. These individuals belong to 33 distinct three-generation families, with 4-15 offspring in the F2 generation. We utilize three MEI-calling tools (MELT, RUFUS, and TranSurVeyor) to identify *de novo* L1, SVA, and Alu retrotransposition events (Gardner et al. 2017; Rajaby and Sung 2018). With the three-generation pedigrees we are able to assess the parental origin and, for the first time, developmental timing of retrotransposition events. We now provide the first direct estimates of *de novo* Alu, L1, and SVA retrotransposition rates based on WGS data from three-generation families. We find the Alu *de novo* insertion rate to be lower than phylogenetic-based estimates and the L1 and SVA insertion rates to be substantially higher than previous estimates. In addition, our three-generation pedigree data allow us to assess and verify Mendelian transmission patterns of *de novo* MEIs, permitting an evaluation of the accuracy of several popular MEI-calling methods.

## Results

### Identification of a *de novo* AluYb8 element using ME-Scan targeted sequencing protocol

We performed ME-Scan on blood-derived DNA from 464 of the 603 CEPH individuals as a pilot study to detect *de novo* AluYb8/9 elements (Supplemental Table S1). ME-Scan is a targeted sequencing protocol that amplifies specific regions of the genome prior to Illumina sequencing (Witherspoon et al. 2010; Ha et al. 2016; Witherspoon et al. 2013; Feusier et al. 2017; Ha et al. 2017). This protocol targets the 7bp insertion that is diagnostic of Yb8/9 elements (Witherspoon et al. 2013, 2010; Feusier et al. 2017). We identified and validated one *de novo* Alu Yb8 insertion, Alu #1 (Table 1 and Supplemental Table S2).

**Table 1:**
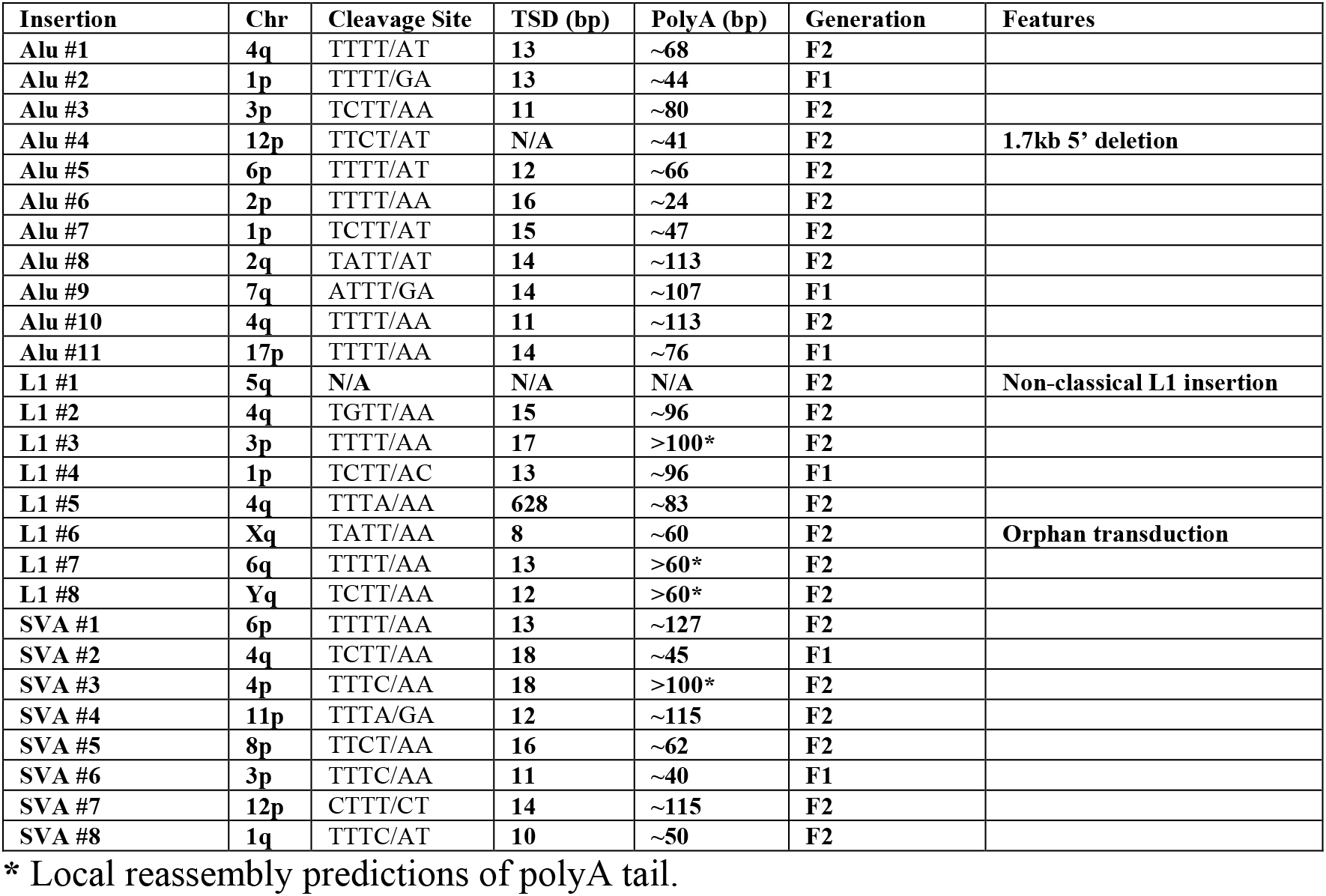
Characteristics of 27 *de novo* MEIs identified in 440 births

### Analysis of *de novo* MEIs in Whole-Genome Sequences of 33 three-generation pedigrees

Blood-derived DNA samples from 603 individuals in 33 three-generation pedigrees were whole-genome sequenced at an average depth of ~30X using Illumina paired-end technology (Supplemental Table S3). MELT identified 911 candidate *de novo* loci from 12,598 called Alu, SVA, and L1 loci. These candidates were evaluated in IGV for characteristic signatures of MEIs, including a target site duplication (TSD), a polyA tail, and split/discordant reads with pairs that mapped to a retrotransposon family (Methods, Supplemental Data S1). Twenty loci met these criteria, and all 20 loci were validated via PCR and Sanger sequencing (Supplemental Figures S1 and S2). TranSurVeyor identified 86,745 breakpoints, including 15 of the 20 identified by MELT and an additional six loci not found by MELT. (Supplemental Data S2). Of these 26 *de novo* MEIs, the RUFUS algorithm called 23 from 56,899 breakpoints (Supplemental Data S3) and found another *de novo* locus (https://github.com/jandrewrfarrell/RUFUS). In total, we identified and PCR-validated eight L1, eight SVA, and 11 Alu mobile element insertions in 17 of 33 CEPH pedigrees (Table 1, Supplemental Figures S1 and S2, and Supplemental Tables S4-S7). PCR validiation showed that every locus with preliminary evidence of a MEI event was a true-positive *de novo* insertion.

MELT uses a transposon reference file to identify and characterize non-reference MEI for each transposon family (Gardner et al. 2017). In contrast, RUFUS and TranSurVeyor identify breakpoints regardless of the transposon family, each producing tens of thousands of false-positive breakpoints (Rajaby and Sung 2018). Our results demonstrate the importance of utilizing different tools for MEI detection (Rishishwar et al. 2016; Ewing 2015; Goerner-Potvin and Bourque 2018), because only 52% of validated *de novo* MEIs were detected by all three tools, and 11% of *de novo* MEIs were detected by a single tool (Supplemental Table S4).

With our three-generation pedigrees, we were able to identify obligate carriers of a MEI in the F1 generation as individuals whose parent (P0) carried the MEI and whose offspring (F2) inherited the MEI (Supplemental Figure S3). This allowed us to estimate MELT’s sensitivity to detect all MEIs as 94% when using its standard filters (which exclude many incorrect calls) and 68% without filters. For *de novo* MEIs only, we estimate sensitivity values for MELT, RUFUS, and TranSurVeyor calls as ~74%, 88%, and 77%, respectively. Using only loci that passed MELT’s filters (i.e. “PASS”) would have reduced the *de novo* candidate list from 911 to 218 loci, but 40% (8 of 20) of the *de novo* loci would have gone undetected.

Twenty-five *de novo* MEIs contain all of the hallmarks of L1-mediated retrotransposition: a polyA tail, a TSD, and the endonuclease cleavage site motif (5’-TTTT/AA-3’) (reviewed in (Cordaux and Batzer 2009; Hancks and Kazazian 2016)). Alu #4 is full-length but has a 1.7kb deletion at its 5’ end, which may have occurred during insertion, and thus does not have a TSD. L1 #1 is 5’ and 3’ truncated, does not have hallmarks of retrotransposition, and contains a deletion of an “A” at the insertion site. This indicates a non-classical L1 insertion (NCL1) event, which is hypothesized to play a role in double-stranded break repair (Sen et al. 2007).

The genomic context of the 27 *de novo* insertions events is shown in Figure 1. Detailed information on each breakpoint is provided in Supplemental Figure 2. The MEIs are randomly distributed across the genome (Figure 1A). Forty-one percent of the loci inserted outside of repetitive DNA regions (Figure 1B). L1 #7 inserted 25bp away from exon 4 in *PM20D2* (Supplemental Figure S2). L1 # 5 inserted within the 3’ UTR of *PGRMC2* and created a 628bp TSD (Supplemental Figure S2). As expected for a non-disease cohort and this number of MEIs, we did not find any *de novo* MEIs in exons.

**Figure 1:**
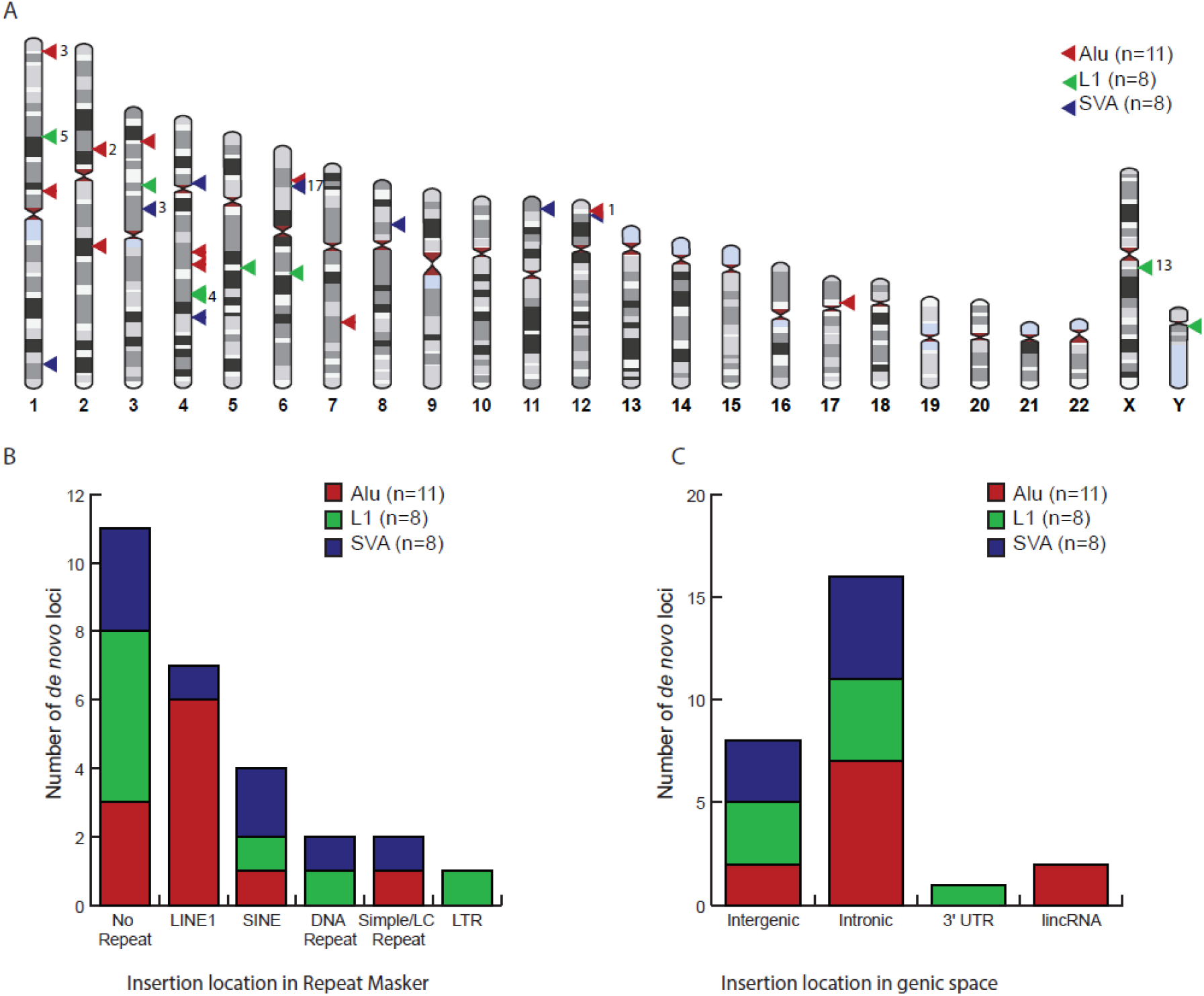
Distribution of *de novo* MEIs throughout the genome. A: Genomic map of *de novo* MEIs using <u>HumanIdiogramLibrary</u> (Collins). The numbers to the right of the triangles indicate the chromosome of the source element. B: Repeat Masker (UCSC genome browser) context of *de novo* MEIs (Kent et al. 2002). C: Genic context of *de novo* MEIs (UCSC genome browser) (Kent et al. 2002). The genomic context in B and C was determined using the TSD region of each locus.

**Figure 2:**
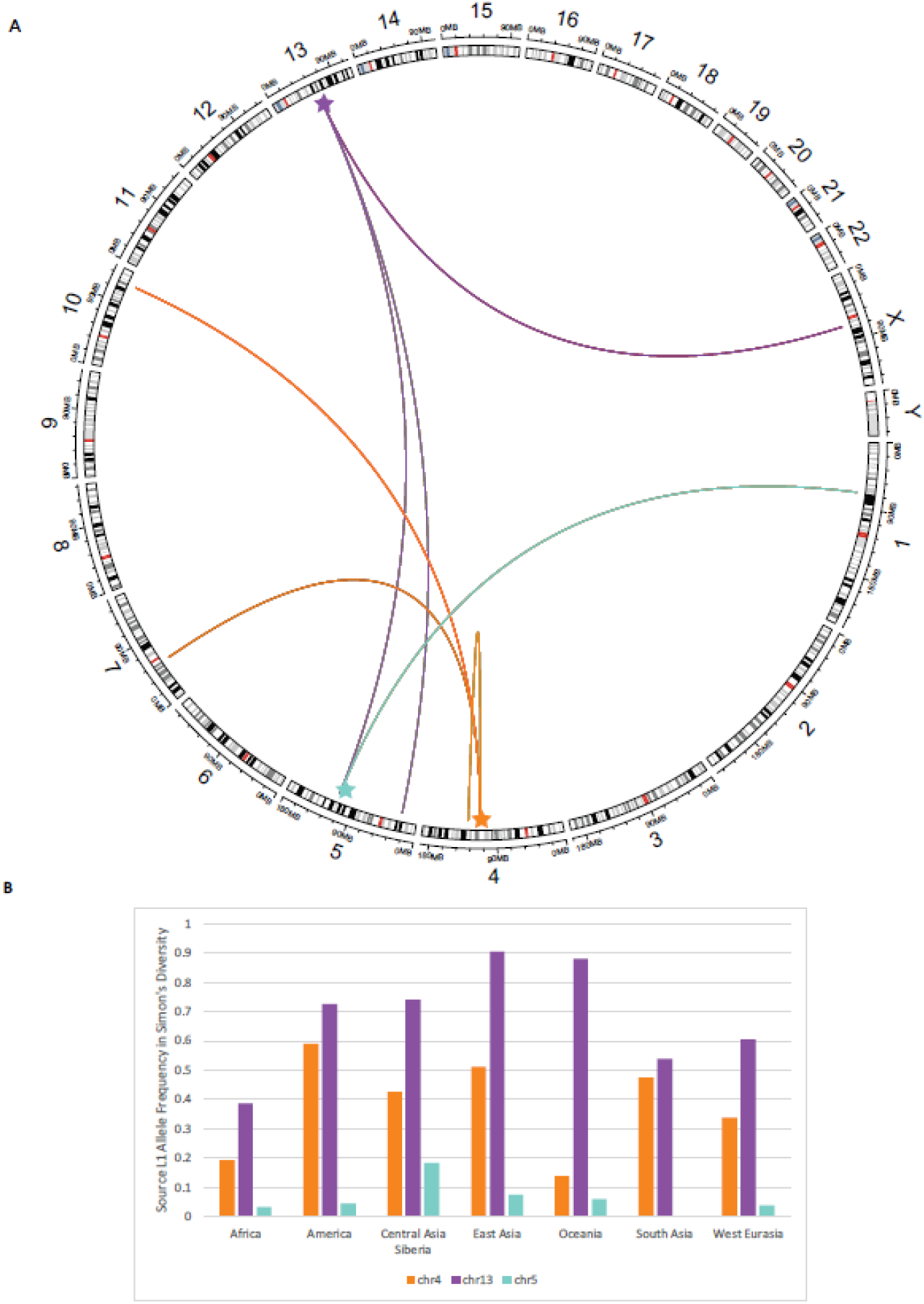
Three source L1 elements identified by 3’ transductions. A: Circlize plot of L1 elements to identified offspring elements in the CEPH dataset (Gu et al. 2014). Source elements are highlighted with a star. B: Allele frequency of the three source elements in the Simons Genome Diversity Project (Mallick et al. 2016). Genotypes were manually typed from IGV screenshots (Supplemental Table S9).

### Subfamily analysis of the *de novo* MEIs

The numbers of *de novo* MEIs in different subfamilies are shown in Table 2. Eleven Alu elements belonged to seven subfamilies. Alu elements #1 and #5 are exact matches to the Yb8 subfamily, while Alu #8 and #9 belong to the Ya5 subfamily. Alu #10 is truncated by >250bp and can theoretically belong to many Y (or the older S) subfamilies (Kryatova et al. 2017). (Sequence alignment and fasta files for the 11 Alu elements are presented in Supplemental Figure S4 and Supplemental Data S4.) We matched the full length L1 #2 to the young L1Ta1d subfamily. We did not get sequence information for the other full-length L1 (L1 #7), and the other six elements are too truncated for classification. SVA #4-6 and #8 contain part of the 5’ transduction of *MAST2* exon 1 and therefore belong to the SVA_F1 subfamily (Damert et al. 2009; Hancks et al. 2009; Bantysh and Buzdin 2009). SVA #5 additionally contains the 3’ transduction of an AluSp, which is present in the SVA_F1 master element H10_1 (Damert et al. 2009; Hancks et al. 2009). The sequences of the SINE-R regions for SVA #1-3 and #7 align to the other known active subfamilies, D-F. The subfamily assignment for each element is in Supplemental Table S4.

**Table 2:**
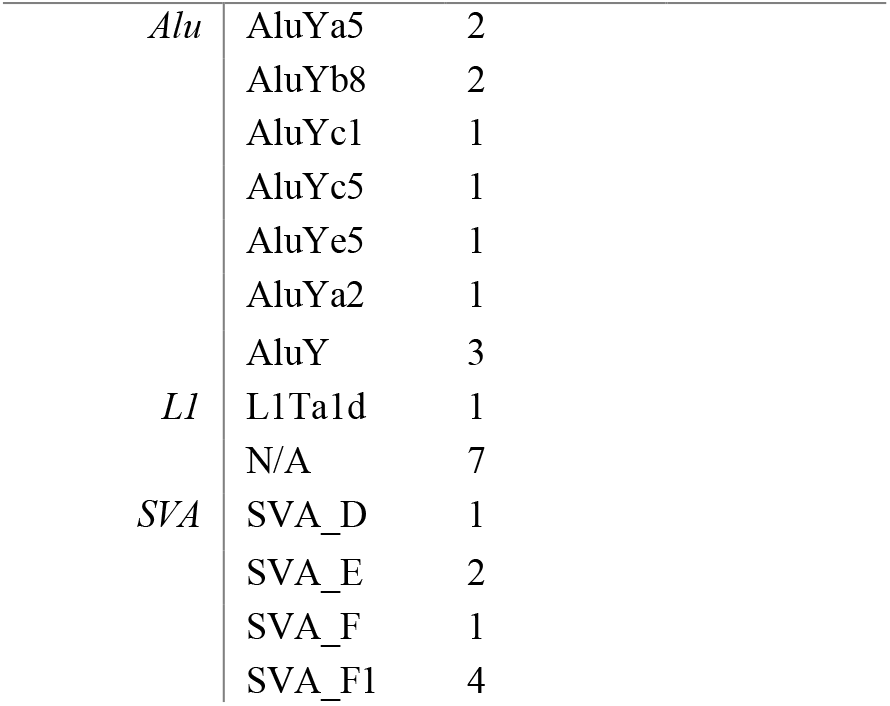
Subfamily composition of de novo MEIs

|  |  |  |
| --- | --- | --- |
| Alu | AluYa5 | 2 |
|  | AluYb8 | 2 |
|  | AluYc1 | 1 |
|  | AluYc5 | 1 |
|  | AluYe5 | 1 |
|  | AluYa2 | 1 |
|  | AluY | 3 |
| L1 | L1Ta1d | 1 |
|  | N/A | 7 |
| SVA | SVA_D | 1 |
|  | SVA_E | 2 |
|  | SVA_F | 1 |
|  | SVA_F1 | 4 |

### Several *de novo* MEIs have hallmarks of retrotransposition activity

To determine whether any of the *de novo* Alu elements are capable of further retrotransposition, we examined each element for its potential capacity for retrotransposition activity. Hallmarks of active *Alu* elements include intact box A and B internal RNA polymerase III (pol III) promoters (Mills et al. 2007; Bennett et al. 2008; Comeaux et al. 2009), intact SRP9/14 sites, an uninterrupted poly(A) tail at least 20 bases long (Dewannieux and Heidmann 2005), and a pol III termination sequence, TTTT, preferably within 15bp of the TSD downstream of the poly(A) tail (Comeaux et al. 2009). In addition, there are 124 conserved nucleotides in active Alu elements, and multiple mutations in these nucleotides may affect retrotransposition efficiency (Bennett et al. 2008). Alu elements #1 and #8 contain all of these hallmarks and therefore may be active (Supplementary Data S4).

To identify potentially active L1/SVA elements, we focused on the full-length *de novo* elements in our dataset. L1 #2 is potentially active because it is not truncated relative to its source element and has two intact open reading frames (ORFs 1 and 2) as determined by L1Base2 (Penzkofer et al. 2017). L1 #7 is full-length, but we were unable to sequence the ORFs to determine activity potential. The other six L1 elements are 5’ truncated and therefore not active. SVA #2 is the only element with the CCCTCT hexamer promoter and may be active, although we were unable to sequence through the VNTR region. SVA #6 and #8 are *de novo* SVA_F1 elements with the full *MAST2* promoter and therefore could be active. The other SVA elements do not contain the CCCTCT hexamer but may be transcribed if they inserted downstream of a promoter.

### Identification of source elements

We used the human reference genome (hg19) and reconstructed fasta files from the MELT output to identify potential source elements of the *de novo* MEIs (Figure 1, Methods). Alu #2 and #4 each had a unique match to a reference Alu element (hg19 chr3:190,156,698-190,156,966 and chr1:246470713-246471020, respectively). Alu #3 is 40bp truncated but uniquely matches a full-length polymorphic Alu element identified by MELT (hg19 chr2:185125618). SVA #3 is identical to a reference SVA_D element (hg19 chr17:42314401-42316970) except for a 725bp deletion region as a result of splicing (Supplemental Figure S2). SVA #6 contains a 22bp deletion within the *MAST2* promoter, which is unique to an SVA_F1 element (hg19 chr3:48251893-48254907) (Damert et al. 2009). There were too many potential source elements to pinpoint the candidate source element for the remaining eight Alu and six SVA elements.

We identified the unique source element for the three L1 elements with 3’ transductions (Figures 1 and 2). L1 #2 contains an 82bp 3’ transduction that maps to an active L1 on 4q25. L1 #4 contains an 846bp 3’ transduction from a L1 on chr5q22. L1 #6 is a 497bp orphan 3’ transduction (i.e., the entire L1 was 5’ truncated) that maps to the 3’ end of a ~2kb 3’ transduction from chr13q21.2. We identified four additional 3’ transduction events from these source elements in our dataset by examining the source loci in IGV (Figure 2A, Supplemental Table S8). All three source elements are non-reference insertions, polymorphic across nearly all of the major population groups in the Simons Genome Diversity Project, and are active in cancer genomes (Figure 2B and Supplement Figure S5) (Mallick et al. 2016; Tubio et al. 2014).

### Transmission of *de novo* F1 MEIs

We analyzed haplotypes of six F1 MEIs (three Alu, one L1, and two SVA) to infer the stage at which the retrotransposition event occurred during development. Using the haplotype of the F2 individual(s) that inherited the *de novo* MEI, all six F1 insertions were phased to the paternal (maternal grandfather) chromosome (Supplemental Tables S10-S15). The three F1 Alu elements were transmitted to the F2 at roughly Mendelian ratios, and the Alu insertions were always co-inherited with the maternal grandfather’s chromosome. However, L1 #4 and SVA #2 and #6 are not transmitted at the expected ratios (χ2 test with 1 degree of freedom, two-tailed p-value < 0.02). These MEIs were only transmitted to one offspring each, and there were multiple offspring in each pedigree that inherited the maternal grandfathers’ haplotypes but not the MEIs. Examples of these transmission events are shown in Figure 3. Both the transmission frequency and haplotype inconsistencies indicate that the L1/SVA insertions are germline/somatic mosaic in the F1 individuals, in contrast to the F1 Alu insertions.

**Figure 3:**
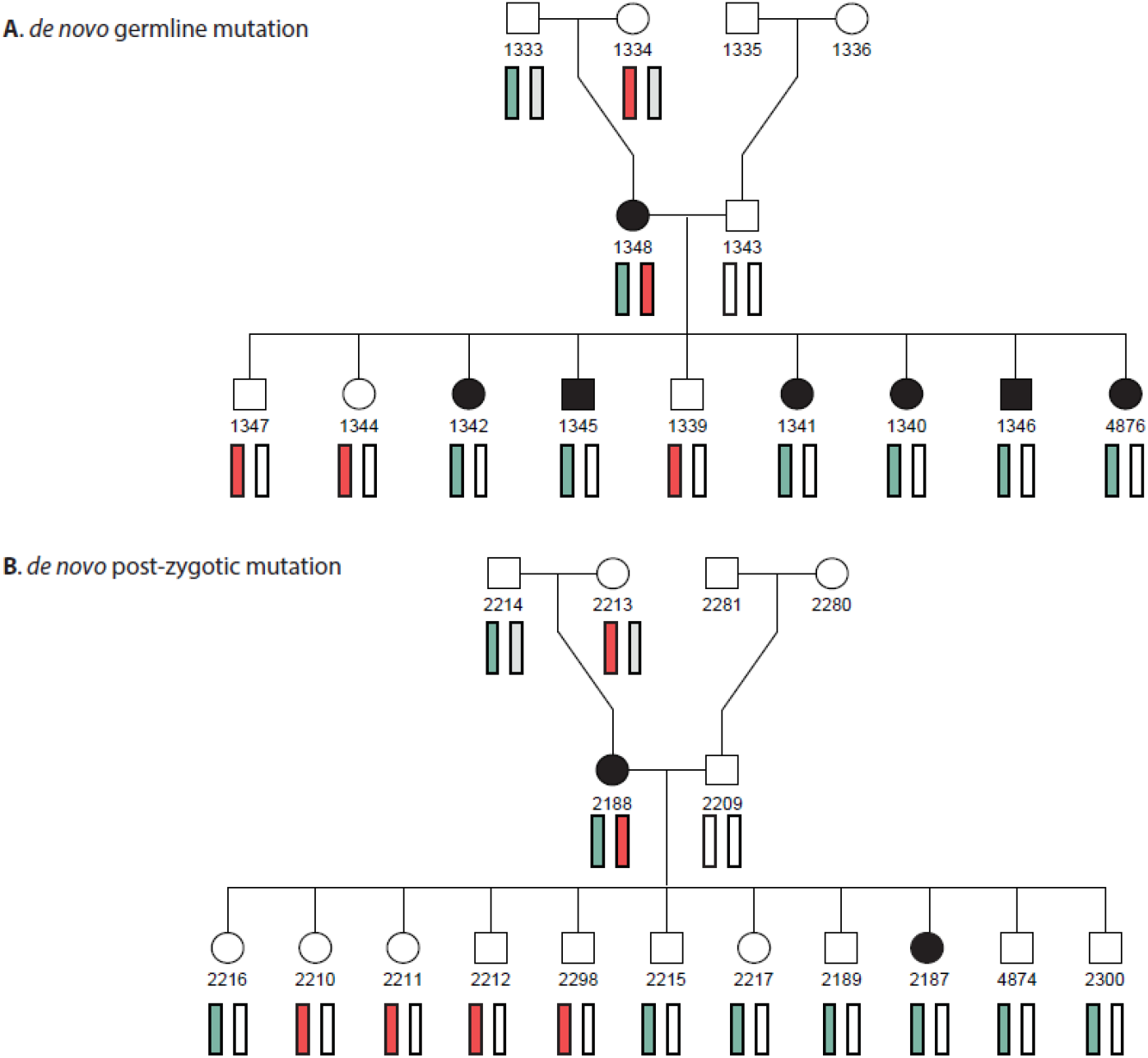
Tracking *de novo* retrotransposition in multigenerational pedigrees. The maternal grandfather’s haplotype is shown in light blue and maternal grandmother’s haplotype is shown in red. A: Evidence of *de novo* retrotransposition in the germline. This panel illustrates co-transmission of Alu #9 (filled individuals) and the maternal grandfather’s haplotype in Pedigree 1423. B: Evidence of *de novo* retrotransposition in the post-zygotic stage. This panel illustrates the germline mosaic transmission of SVA #7 with the maternal grandfather’s haplotype in Pedigree 1463. Six other siblings inherited the grandfather’s haplotype without the SVA.

### Parental Origin of *de novo* MEIs

We identified the parental origin of 17 (six F1 and 11 F2) of the *de novo* MEIs using sex chromosome hemizygosity, haplotype phasing, and SNP-based phasing approaches (Methods). Eight Alu elements were transmitted on the paternal chromosome, and one element was transmitted on the maternal chromosome (exact binomial test, p-value < 0.04). Two and three of the SVA elements were transmitted on the maternal and paternal chromosomes, respectively. In addition to the two hemizygous L1 elements on the X and Y chromosomes, another L1 element was transmitted on a paternal chromosome. The average paternal age at conception for paternally-phased de novo MEI (31.21 years) is the nearly identical to the total average father age (31.41 years) (two tailed p-value > 0.9254). The average mother’s age for children with phased de novo MEI is four years younger than the average age (24.2 vs 28.5 years) but not statistically different (two-tailed p-value < 0.232). We conclude that there is a statistically significant paternal sex bias with respect to Alu retrotransposition, and that, although the data are limited, there were no parental age biases in retrotransposition.

## Discussion

With rapid advances in next-generation sequencing technology, a large number of human pedigrees have been sequenced, and many studies have directly estimated the single nucleotide *de novo* mutation rate (Roach et al. 2010; Jónsson et al. 2017). New technology also affords an opportunity to estimate the rate of *de novo* retrotransposition, which generates genomic variation through an entirely different mutation mechanism. From 440 births, we estimate an Alu retrotransposition rate of ~1:40 births (95% CI 22.6-80.0), an SVA rate of ~1:55 births (95% CI 28.2-126.6), and a L1 rate of ~1:62 births (95% CI 30.8-156.3) (Figure 4).

**Figure 4:**
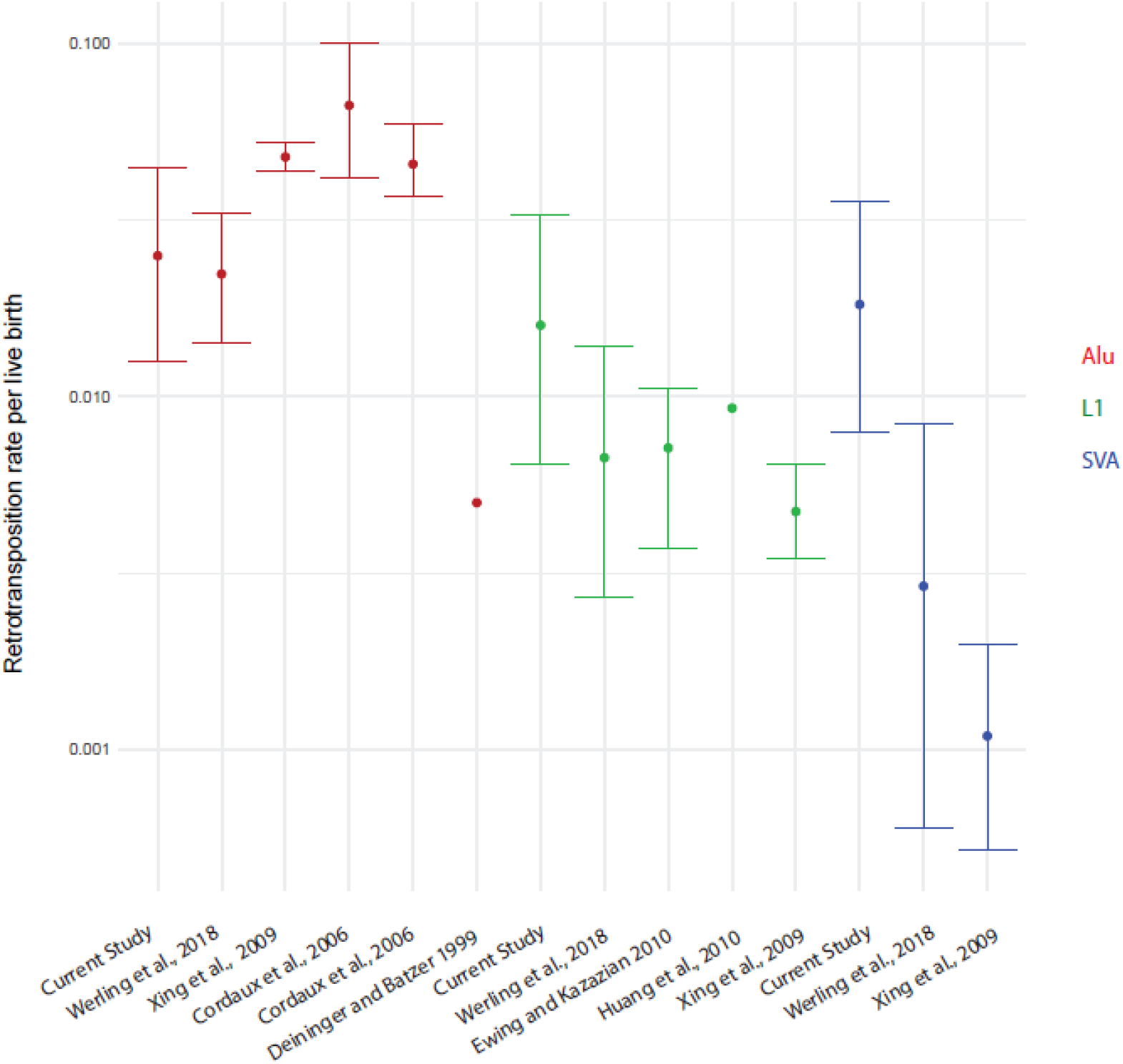
Estimated germline retrotransposition rates. Estimated germline retrotransposition rates for previous studies are listed (Xing et al. 2009; Cordaux et al. 2006; Deininger and Batzer 1999; Ewing and Kazazian 2010). Confidence intervals are shown if available from the study. Rates and binomial 95% C.I. were determined for (Werling et al. 2018) and this study. Alu element rates are shown in red, L1 in green, and SVA in blue.

MELT was used previously to identify *de novo* MEI transmission in 519 quartets in the Simons Simplex Collection (SSC) (Werling et al. 2018). Using these published data, we estimated comparative retrotransposition rates for Alu, L1, and SVA elements (Figure 4). The Alu retrotransposition rate in SSC is nearly identical to the estimate in this study, but our L1 and SVA retrotransposition rates are 2.4x and 6.3x higher but do not differ significantly (see confidence limits in Figure 4) (Werling et al. 2018). The latter differences reflect in part our use of multiple MEI-calling tools, which showed that MELT detects 91% of *de novo* Alu elements but only 75% of L1 and 50% of SVA elements. These two datasets both estimate an Alu retrotransposition rate that is 2-fold lower than previous phylogenetic and disease-based estimates. Given MELT’s high sensitivity for Alu detection (Gardner et al. 2017) as well as the use of multiple MEI-calling tools, it is unlikely that our lower rate is due to false negative calls. Instead, it is likely that the phylogenetically estimated rate is affected by assumptions about the divergence time of humans and chimpanzees, the effective population size of the human-chimpanzee ancestral population, and retrotransposition rate variation over time (Roach et al. 2010; Cordaux and Batzer 2009; Campbell and Eichler 2013; Ségurel et al. 2014).

While preliminary, our results suggest there may be differences in retrotransposition timing among the non-LTR retrotransposon families. All 3 Alu elements in the F1 generation conform to Mendelian expectations and co-segregate with the paternal grandfather’s chromosome, indicating retrotransposition in the germline. Further, there is a paternal sex bias in *de novo* Alu retrotransposition, which is similar to the paternal transmission bias seen in single nucleotide variants and short tandem repeats (Willems et al. 2017; Jónsson et al. 2017). We found evidence of L1 retrotransposition events in both the germline (L1 #6 and #8) and in early embryogenesis (L1 #4) (Figure 3), which corroborates previous findings (Richardson et al. 2017; van den Hurk et al. 2007). The two F1 SVA elements are suspected to be mosaic in the germline and somatic tissue in the mother and likely arose during embryogenesis. Notably, inheritance of germline/somatic mosaic L1 and SVA elements has thus far only been seen in females: three in this study and four in a recent mouse study (Richardson et al. 2017). The observation of likely post-zygotic SVA element insertions suggests that SVA insertions may be underreported in studies of somatic or tumor cells.

Our data allow us to identify the subfamily distribution of active mobile element subfamilies. Yb8 and Ya5 subfamilies accounted for 72% of 322 polymorphic Alu elements in a recent study (Konkel et al. 2015), yet only 36% of *de novo* Alu elements identified here belong to the Ya5 or Yb8 subfamilies (Fisher exact test, p-value <0.02). Indeed, the variety of *de novo* Alu sequences detected here corroborates the “stealth model” hypothesis of Alu amplification, where there are multiple active subfamilies that proliferate, rather than one large, active subfamily/locus (Konkel et al. 2015; Han et al. 2005; Deininger and Batzer 1999; Deininger et al. 1992). We detected insertions of all active SVA subfamilies, with the youngest SVA_F1 subfamily accounting for half of the *de novo* SVA elements in this study (Damert et al. 2009; Hancks et al. 2009; Bantysh and Buzdin 2009). Our data show that there are many active AluY subfamilies, and the youngest SVA subfamily, SVA_F1, may be currently one of the most active SVA subfamilies.

In addition to the three non-LTR retrotransposon families, there are other substrates of retrotransposition in the human genome. Processed pseudogene insertions occur when processed mRNA is inserted into the genome using the L1 machinery (Ewing et al. 2013; Abyzov et al. 2013; Schrider et al. 2013). There are also several polymorphic HERV-K (HML2) elements in humans, including at least one potentially active insertion (Wildschutte et al. 2016). We searched for HERV-K (HML2) elements using MELT and did not identify any candidate *de novo* loci (Supplemental Data S1, Methods). RUFUS and TranSurVeyor did not detect any *de novo* pseudogene or HERV-K (HML2) insertions. Processed pseudogene retrotransposition events are rare (Gardner et al. 2018), and tools specific to identifying these events in WGS would allow for retrotransposition rate estimates in pedigrees.

Retrotranspositional activity may differ across pedigrees and populations, similar to the way in which polymorphic PRDM9 variants affect recombination hotspot activity (Kong et al. 2010; Baudat et al. 2010). It is predicted that every human contains 80-100 active L1 elements (Brouha et al. 2003), and this may influence variation in retrotransposition activity among humans. The three source L1 elements in this study are present in all major regional groups in the Simons Genome Diversity Project dataset (Mallick et al. 2016), which suggests that these elements may also be active in non-European populations (Figure 2 and Supplemental Figure S5). We identified an overabundance (six) of *de novo* MEIs in pedigree 1331; siblings 8549 and 8310 were also the only individuals with more than one *de novo* MEI in the dataset (Alu #1 and SVA #5 in 8549, and L1 #5 and SVA #1 in 8310). Preliminary analysis of the pedigree did not reveal any pathogenic SNPs in a gene list of proteins that restrict retrotransposition activity (Goodier 2016). Future studies of retrotransposition in large pedigree-based cohorts may help to elucidate variants and genetic factors involved in the regulation of L1-mediated retrotransposition activity.

## Methods

### CEPH individuals

Blood-derived DNA samples from 603 individuals, including 458 trios within larger pedigrees, were collected from either the original CEPH cohort (Dausset et al. 1990) or the Utah Genetic Reference Project (Prescott et al. 2008). These samples were whole-genome sequenced at ~30x coverage at Washington University in St. Louis. Evaluation by peddy identified ten individuals with a het_ratio > 0.2 who were also declared duplicates, indicating sample contamination prior to sequencing (Pedersen and Quinlan 2017). All 18 trios with these individuals were removed from the rate estimate post-IGV evaluation. Therefore, 440 births were used in the rate estimates.

### Identification of MEIs in the CEPH dataset

We used three complementary approaches to identify *de novo* MEIs in this dataset. All 603 individuals were joint-called with the Melt-Split protocol in MELT (v2.14) for detection of Alu, L1, SVA, and HERVK (HML2) elements using the consensus transposon files provided by MELT (Gardner et al. 2017). Approximate coverage estimates for the IndivAnalysis step were calculated using the covstats tool (goleft v0.1.17 https://github.com/brentp/goleft). To increase sensitivity, loci were not filtered using the filtering criteria provided by MELT. To identify *de novo* MEIs in F1 and F2 simultaneously, Genotype Query Tools (Layer et al. 2015) was used to identify loci that were restricted to a unique CEPH pedigree and homozygous reference in the grandparents (and parents missing at least one grandparent) in that family. All 458 trios were processed through RUFUS, and all structural variant breakpoints were extracted for detection of L1-associated retrotransposition events (https://github.com/jandrewrfarrell/RUFUS). RUFUS was unable to successfully process trios 1788, 2020, and 4877. Each sample was individually processed through TranSurVeyor, and unfiltered breakpoints with less than 4 discordant reads of support were removed (Rajaby and Sung 2018). Then, we merged overlapping breakpoints in each individual using the bedtools merge command (Quinlan and Hall 2010), and merged samples into three bed files: F1, F2, and P0. We next used bedtools intersect to identify F1 loci absent in the P0, and F2 loci absent in the F1 and P0 bed files. Sensitivity of the tools were measured after all 27 *de novo* MEIs were validated. Each locus was visualized as a trio in IGV to identify candidate *de novo* MEIs (Robinson et al. 2013).

### ME-Scan Protocol

ME-Scan was performed on blood-derived DNA of 464/603 CEPH individuals prior to sequencing on the Illumina 2000 platform. Two individuals (grandparent and offspring) failed sequencing and were dropped from the analysis. Data were mapped to hg19 using bwa align (bwa version 0.7.9a) (Li and Durbin 2009) and uploaded to SQL developer for analysis. Read set processing was the same as described in (Witherspoon et al. 2013). A detailed report of the ME-Scan protocol including primers is reported in (Feusier et al. 2017) and Supplementary Methods. *De novo* elements were detected by finding loci that were present in at least one offspring and absent in the parents/grandparents. Loci with less than eight reads of support were removed, based on (Feusier et al. 2017).

### Analysis of candidate breakpoints for validation

After IGV evaluation, the *de novo* TE insertion breakpoints provided by the three tools were further analyzed by extracting the reads mapped to a 250 bp region flanking the breakpoint in each individual. Discordant reads mapped to that 500 bp window were identified and mates of those discordant reads mapped elsewhere in the bam file were collected (Li et al. 2009 and http://broadinstitute.github.io/picard/). A local *de novo* assembly of all the extracted reads was performed (Huang and Madan 1999) for each breakpoint in each individual. The assembled contigs were further probed for the presence of TEs. A custom perl script was written to perform these steps and is available upon request.

### PCR/Sanger Validation of de novo MEIs

PCR amplifications of ~10-25ng of template DNA (blood-derived or transformed lymphoblast DNA) were performed in 25μl reactions according to the Phusion Hot Start Flex DNA Polymerase protocol (using 5x GC buffer) and Q5 Hot Start DNA Polymerase (using GC Enhancer). The thermocycler conditions were: initial denaturation at 98°C for 30s, 40 cycles of denaturation at 98°C for 10s, optimal annealing temperature (58°C-68°C) for 30s, extension at 72°C for 30s-3 minutes, and a final extension at 72°C for 5 minutes. Every primer set reaction was performed on the pedigree with the candidate *de novo* MEI, a positive control, and H2O. PCR amplicons were run on a 1-2% gel containing 0.12mg/ml ethidium bromide for 75-90 min at 120 V. Gels were imaged using a Fotodyne Analyst Investigator Eclipse machine. Bands were cut out and purified for Sanger sequencing using the Qiagen Qiaquick gel extraction kit.

L1 and SVA elements were amplified using the ThermoFisher Platinum SuperFi DNA polymerase and cloned using ThermoFisher Zero Blunt Topo II/4 kits. We followed the Platinum SuperFi PCR setup for 25ul reactions using 2ul of starting DNA (~5ng/ul). For the PCR procedure, each reaction was denatured at 98°C for 30 seconds, and then amplified for 35 cycles (98°C for 10 seconds, annealing for 10 seconds, an extension of 72°C for 30 seconds or longer based on amplicon size). Annealing temperatures were estimated for each primer pair based on ThermoFisher calculations. A final extension was performed at 72°C for 5 minutes. The Invitrogen PureLink Quick Plasmid MiniPrep Kit was used to extract DNA from the clones. Clones were Sanger sequenced through the whole length of the fragment (Supplemental Table S7). We used several internal primers from previous studies (Scott et al. 2016; Feusier et al. 2017). The 3 F1 L1/SVA elements were analyzed in the F2 offspring due to the availability of DNA.

### Investigation of Source Elements

MELT lists differences from the consensus for each MEI locus in the DIFF section of the INFO column (Gardner et al. 2017). These differences were extracted and converted to fasta format using the MELT consensus transposon fasta file as the reference. A custom python script was used for this step and is available upon request. Each *de novo* MEI sequence was compared to the MELT fasta file using the “grep” command to identify potential source elements. The *de novo* MEIs were also compared to the hg19 reference genome using Blat (Kent 2002; Kent et al. 2002).

### Source elements in Simons Genome Diversity Project

DNA sequences (hg19) for 279 individuals from the Simons Genome Diversity Project were downloaded from the European Nucleotide Archive at the European Bioinformatics Institute (PRJEB9586). DNA samples were remapped to hg38. The locations of the three source elements were converted to hg38 using LiftOver in the UCSC Genome Browser (Kent et al. 2002). Individuals were genotyped from IGV screenshots for each of the three source elements (Supplemental Table S9).

### Analysis of Parental Origin

For haplotype phasing of *de novo* MEIs in the F1 individuals, we extracted SNPs in a 200kb window surrounding the MEI’s position (Supplemental Tables S10-S15). We filtered on SNPs that were heterozygous in the F1 and their parents and absent in the F1’s spouse. The F2 individuals were assigned a P0 haplotype based on the transmission of the SNPs from the P0 individuals. Then, the transmission of the *de novo* MEI was placed on the F2’s P0 haplotype to determine parental origin. There was no evidence of recombination between the *de novo* MEI and the markers on the P0 haplotype.

Parental origin of *de novo* MEIs in F2 individuals was analyzed using sex chromosome hemizygous status and SNP-phasing (Supplemental Tables S4-S5). We considered a hemizygous insertion on a sex chromosome to be a retrotransposition event on the parental chromosome in the germline. For SNP-phasing, we considered informative SNPs to be either heterozygous to one parent and the F2, or heterozygous in the F2 and homozygous ref/alt in the F1. Paired-end reads that connected the MEI and a nearby informative SNP confirmed parental origin. For Alu elements with SNPs < 5kb away, we designed primers that amplified the Alu-SNP region, and confirmed the SNPs of the F2 and their parents via Sanger sequencing (primers listed in Supplemental Table S5).

## Data Access

Data are available from the authors upon request.

## Acknowledgments

Funding for this project was provided by the Utah Genome Project, the George S. and Dolores Doré Eccles Foundation, and NIH grants GM118335 and GM059290 (to L.B. Jorde) and R00HG005846 (to H. Ha and J. Xing). We thank Aaron Quinlan, Brent Pedersen, and Thomas Sasani for helpful discussions. Sanger sequencing was performed at the DNA Sequencing Core Facility, University of Utah. We thank William Richards and Matt Velinder for their help with running and troubleshooting RUFUS. Finally, we wish to thank the investigators who organized the original Utah CEPH collection, in particular Ray White and Mark Leppert, as well as all of the families who generously participated in the project.

## Disclosure Declaration

We have no disclosures.

## Additional Files

Supplemental Figure S1: Pedigree/PCR information for 14 families with *de novo* MEI

Supplemental Figure S2: Detailed description of MEI breakpoint

Supplemental Figure S3: False-negative inheritance rate of MELT in CEPH dataset

Supplemental Figure S4: Alignment of 11 *de novo* Alu elements to AluY consensus

Supplemental Figure S5: Presence of source L1 elements within SGDP populations

Supplemental Methods

Supplemental Data S1: All Alu, L1, SVA, and HERVK loci called by MELT

Supplemental Data S2: Bed file of 86,745 candidate breakpoints called by TranSurVeyor

Supplemental Data S3: Bed file of 56,899 candidate breakpoints called by RUFUS

Supplemental Data S4: Detailed fasta files of 27 *de novo* MEI

Supplemental Tables S1-S15

