## Supplemental Figures 1-5 for "Pedigree-based estimation of human mobile element retrotransposition rates"

**Supplemental Material**

**Supplemental Figure S1.** Pedigree/PCR information for 17 families with *de novo* MEI

**Supplemental Figure S2**. Detailed description of MEI breakpoint

**Supplemental Figure S3.** False-negative inheritance rate of MELT in CEPH dataset

**Supplemental Figure S4.** Alignment of 11 *de novo* Alu elements to AluY consensus

**Supplemental Figure S5.** Presence of source L1 elements within SGDP populations

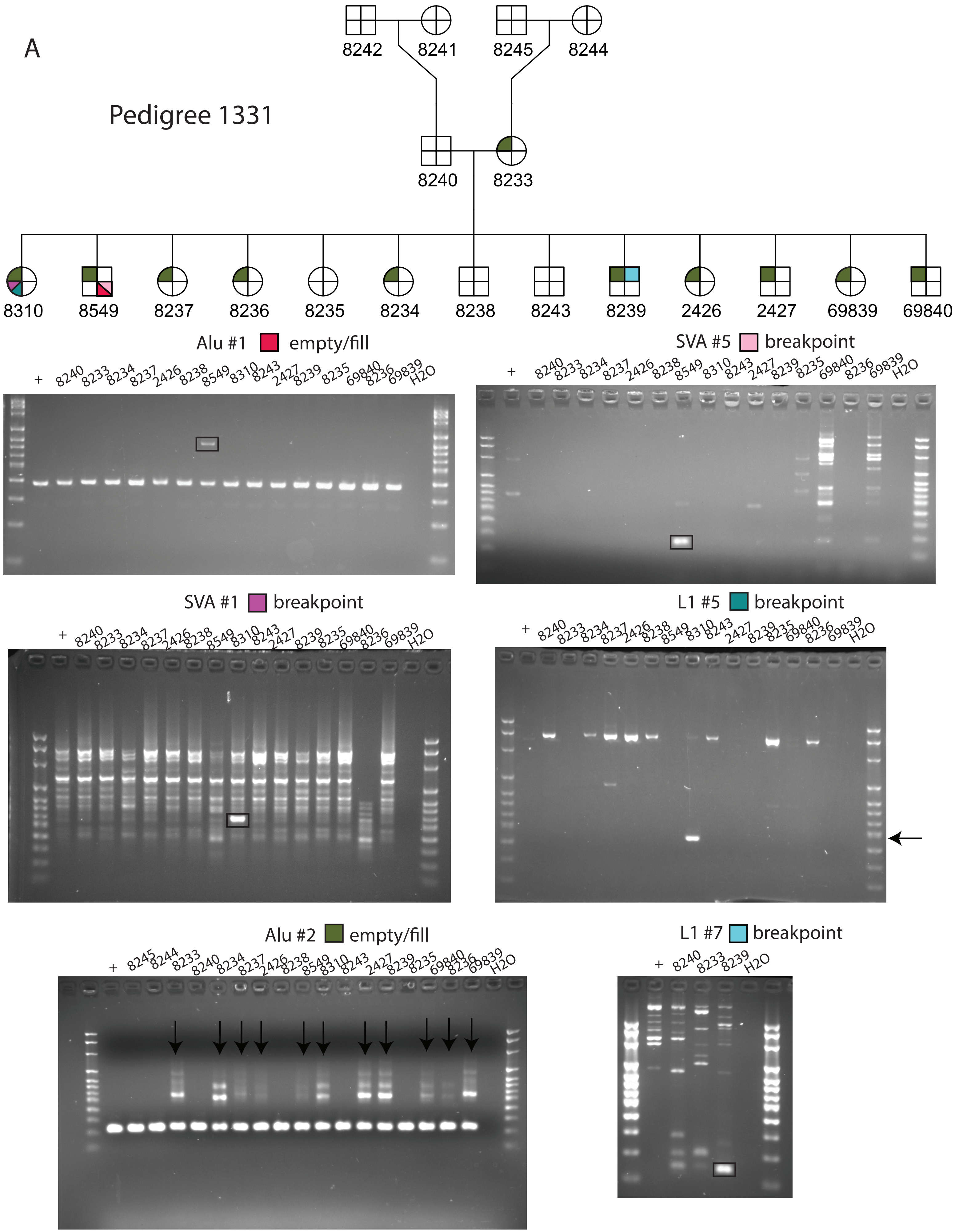
A
Pedigree 1331
8242
8241
8245
8244
8240
8233
8310
8549
8237
8236
8235
8234
8238
8243
8239
2426
2427
69839
69840
Alu #1 empty/fill
+ 8240 8233 8234 8237 2426 8238 8549 8310 8243 2427 8239 8235 69840 8236 69839 H2O
SVA #5 breakpoint
+ 8240 8233 8234 8237 2426 8238 8549 8310 8243 2427 8239 8235 69840 8236 69839 H2O
SVA #1 breakpoint
+ 8240 8233 8234 8237 2426 8238 8549 8310 8243 2427 8239 8235 69840 8236 69839 H2O
L1 #5 breakpoint
+ 8240 8233 8234 8237 2426 8238 8549 8310 8243 2427 8239 8235 69840 8236 69839 H2O
Alu #2 empty/fill
+ 8245 8244 8233 8240 8234 8237 2426 8238 8549 8310 8243 2427 8239 8235 69840 8236 69839 H2O
L1 #7 breakpoint
+ 8240 8233 8239 H2O

B

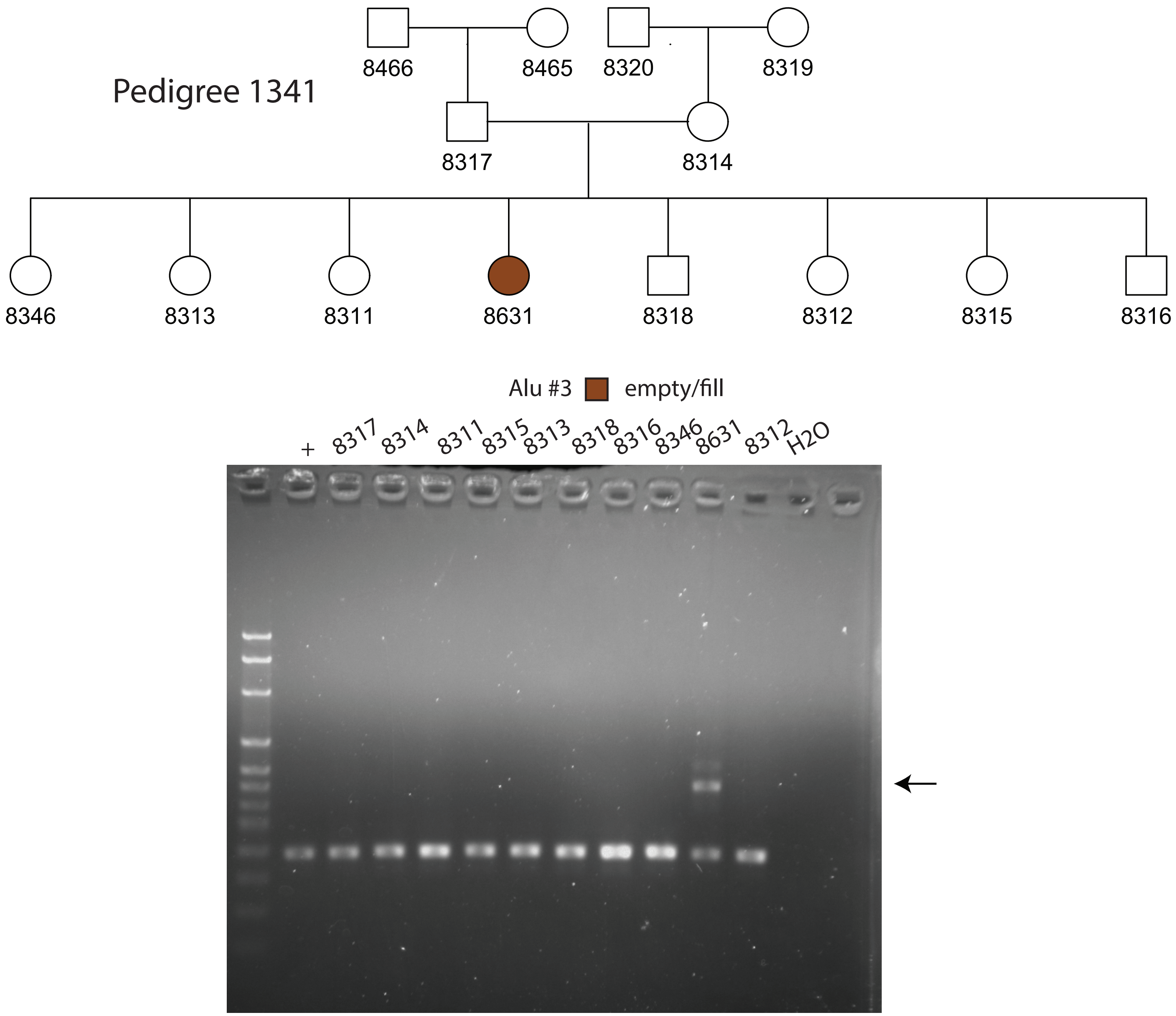

Pedigree 1341
8466
8465
8320
8319
8317
8314
8346
8313
8311
8631
8318
8312
8315
8316
Alu #3
empty/fill
+
8317
8314
8311
8315
8313
8318
8316
8346
8631
8312
H2O

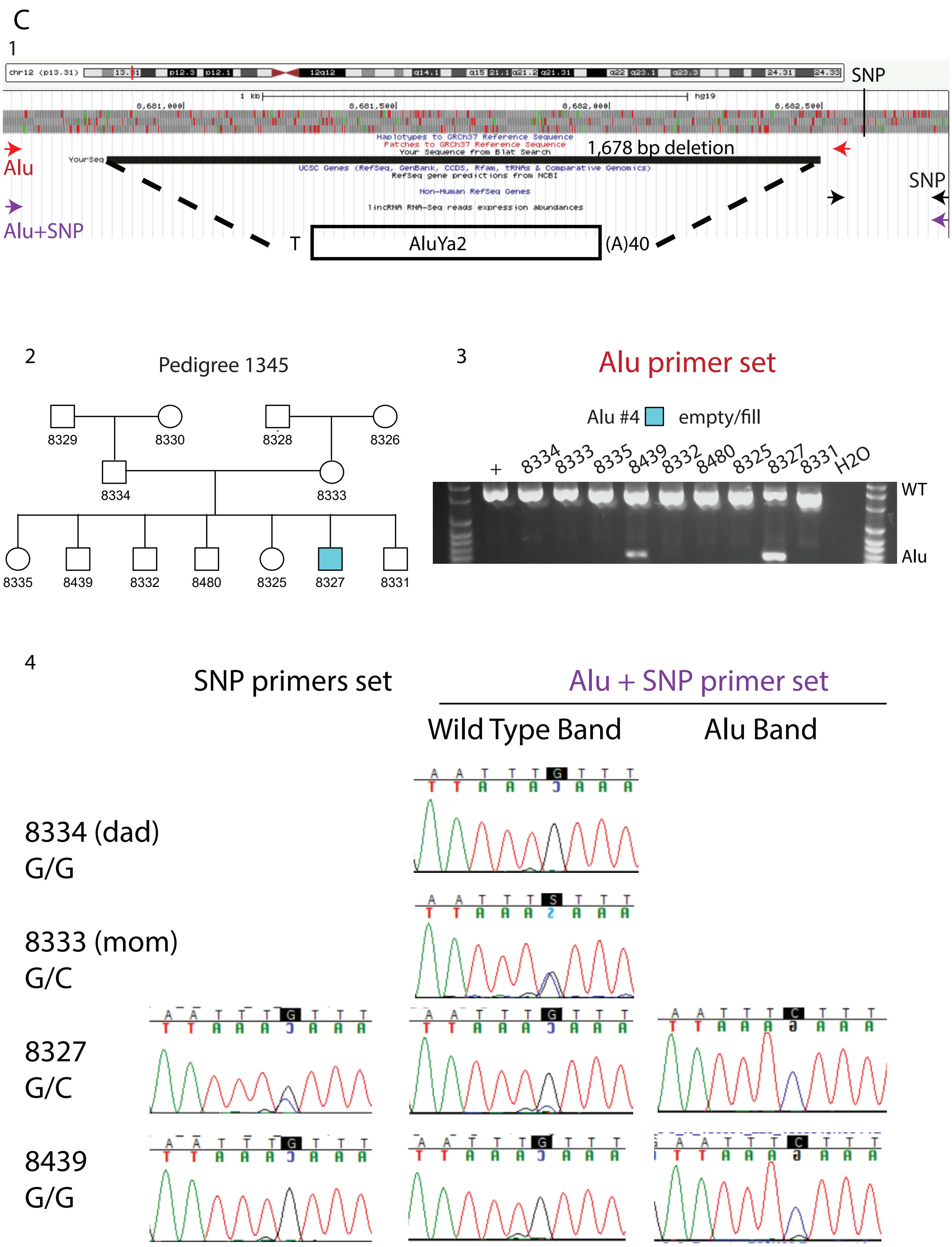
C
1
SNP
1,678 bp deletion
Alu
Alu+SNP
SNP
T
AluYa2
(A)40
2
Pedigree 1345
8329
8330
8328
8326
8334
8333
8335
8439
8332
8480
8325
8327
8331
3
Alu primer set
Alu #4
empty/fill
+
8334 8333 8335 8439 8332 8480 8325 8327 8331 H2O
WT
Alu
4
SNP primers set
Alu + SNP primer set
Wild Type Band
Alu Band
8334 (dad)
G/G
8333 (mom)
G/C
8327
G/C
8439
G/G

D

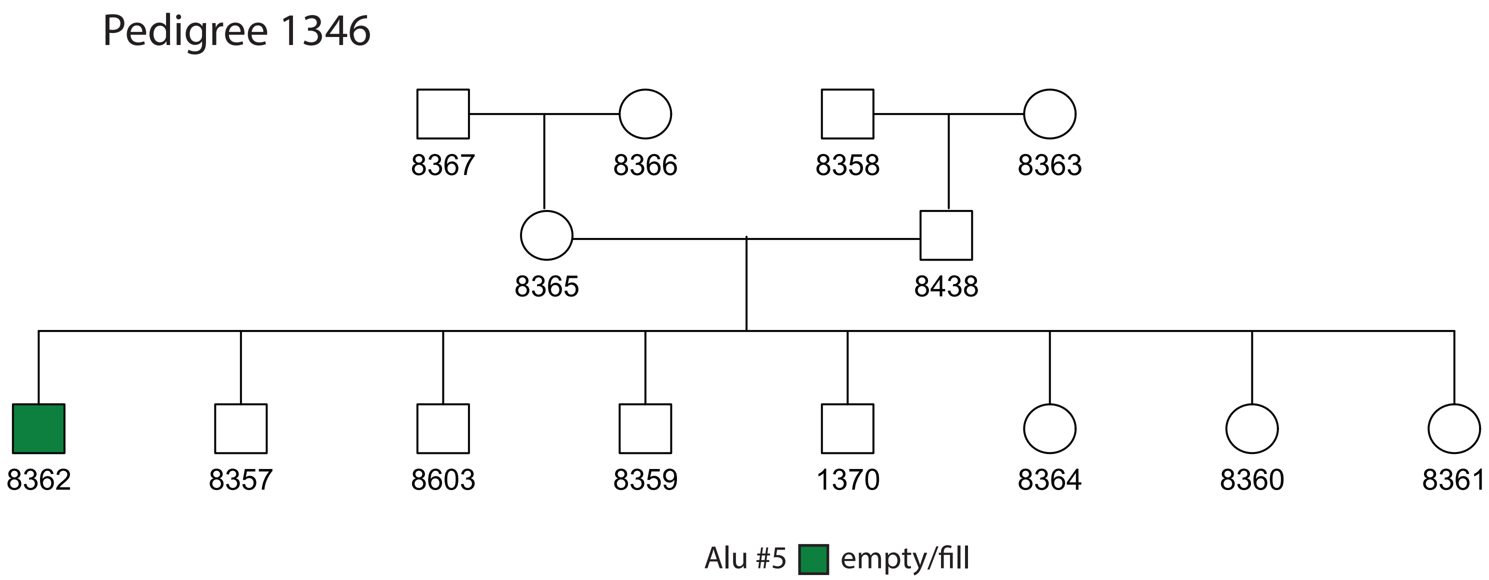

Pedigree 1346
8367
8366
8358
8363
8365
8438
8362
8357
8603
8359
1370
8364
8360
8361
Alu #5
empty/fill


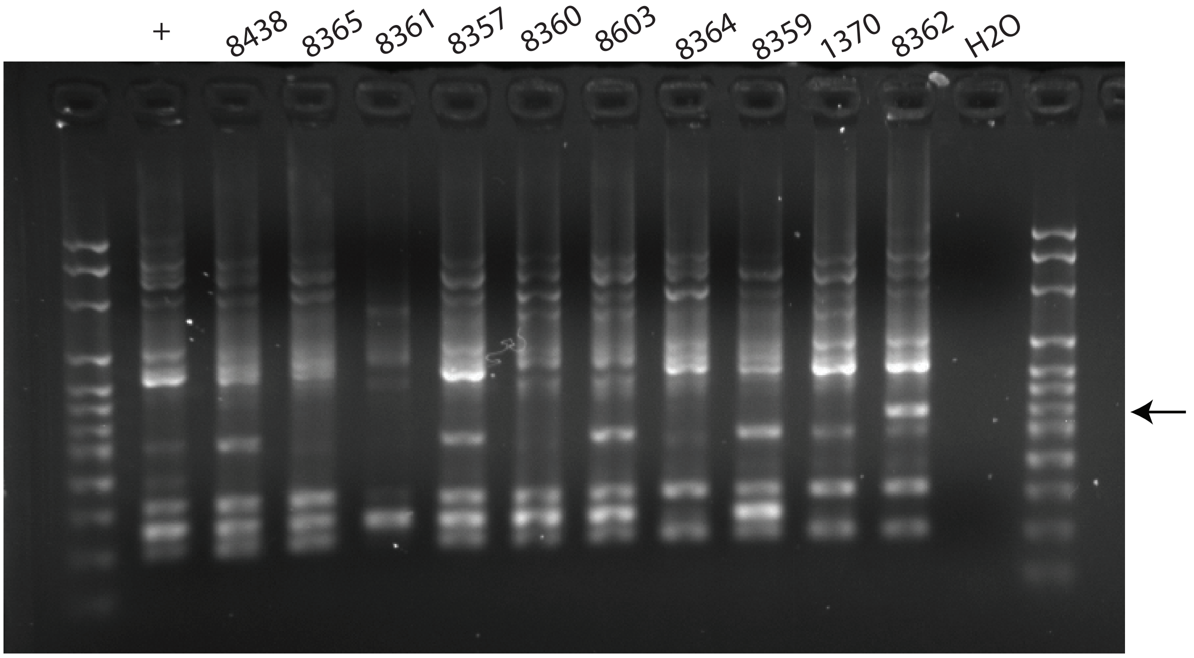

+
8438
8365
8361
8357
8360
8603
8364
8359
1370
8362
H2O

E

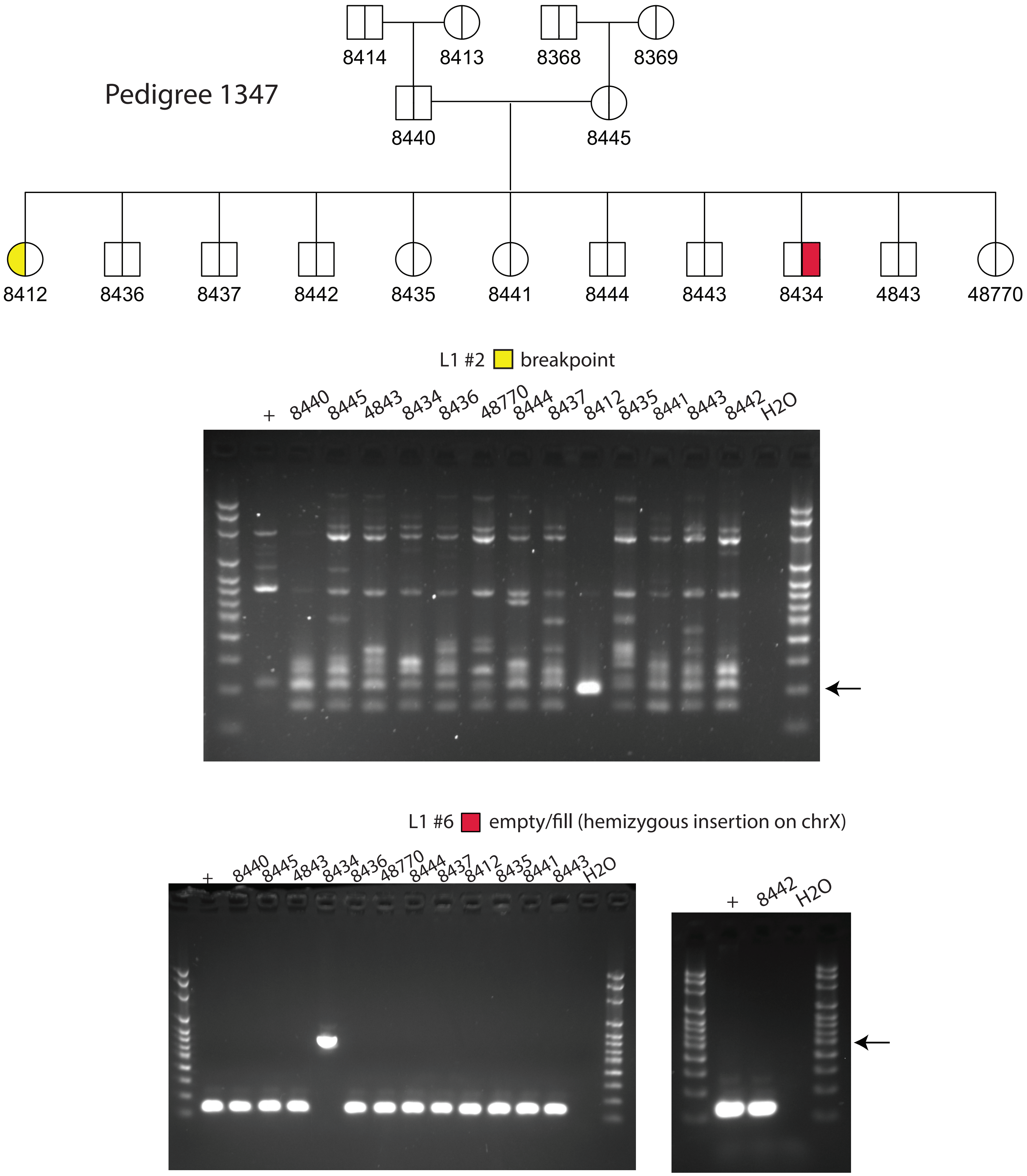

Pedigree 1347
8414
8413
8368
8369
8440
8445
8412
8436
8437
8442
8435
8441
8444
8443
8434
4843
48770
L1 #2 breakpoint
+ 8440 8445 4843 8434 8436 48770 8444 8437 8412 8435 8441 8443 8442 H2O
L1 #6 empty/fill (hemizygous insertion on chrX)
+ 8440 8445 4843 8434 8436 48770 8444 8437 8412 8435 8441 8443 H2O
+ 8442 H2O

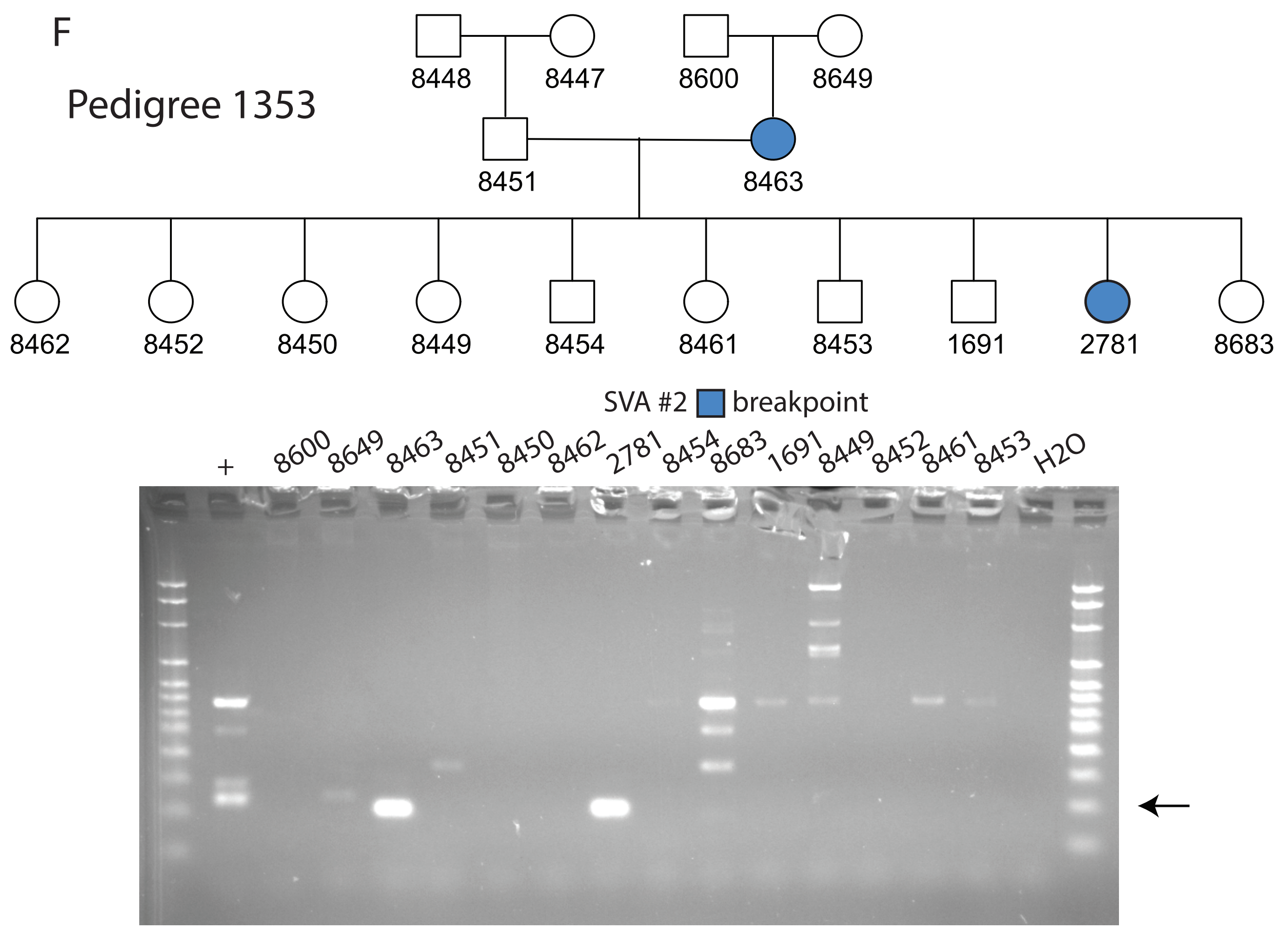
F
Pedigree 1353
8448
8447
8600
8649
8451
8463
8462
8452
8450
8449
8454
8461
8453
1691
2781
8683
SVA #2 breakpoint
+
8600
8649
8463
8451
8450
8462
2781
8454
8683
1691
8449
8452
8461
8453
H2O

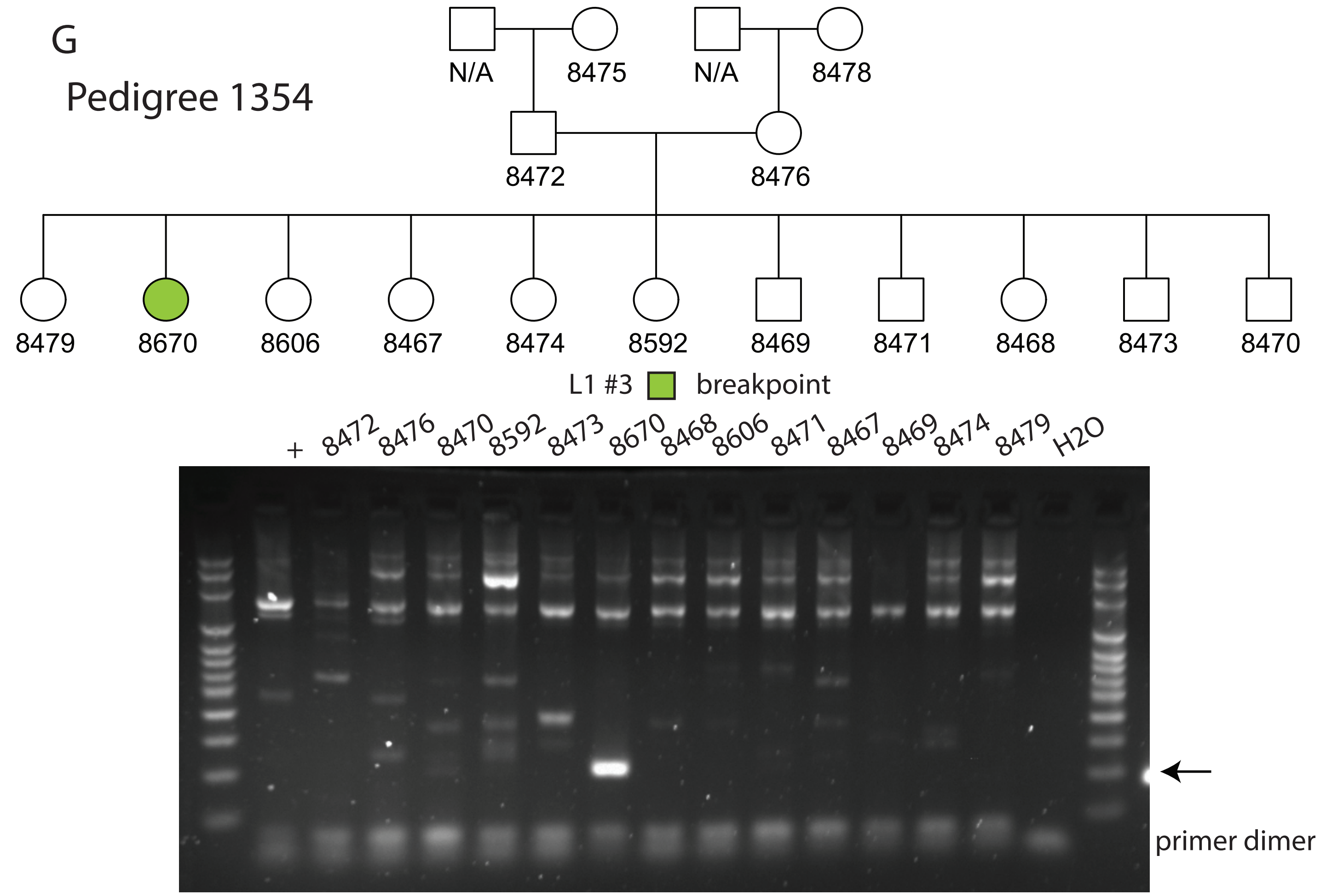
G
Pedigree 1354
N/A
8475
N/A
8478
8472
8476
8479
8670
8606
8467
8474
8592
8469
8471
8468
8473
8470
L1 #3 breakpoint
+
8472
8476
8470
8592
8473
8670
8468
8606
8471
8467
8469
8474
8479
H2O
primer dimer

H

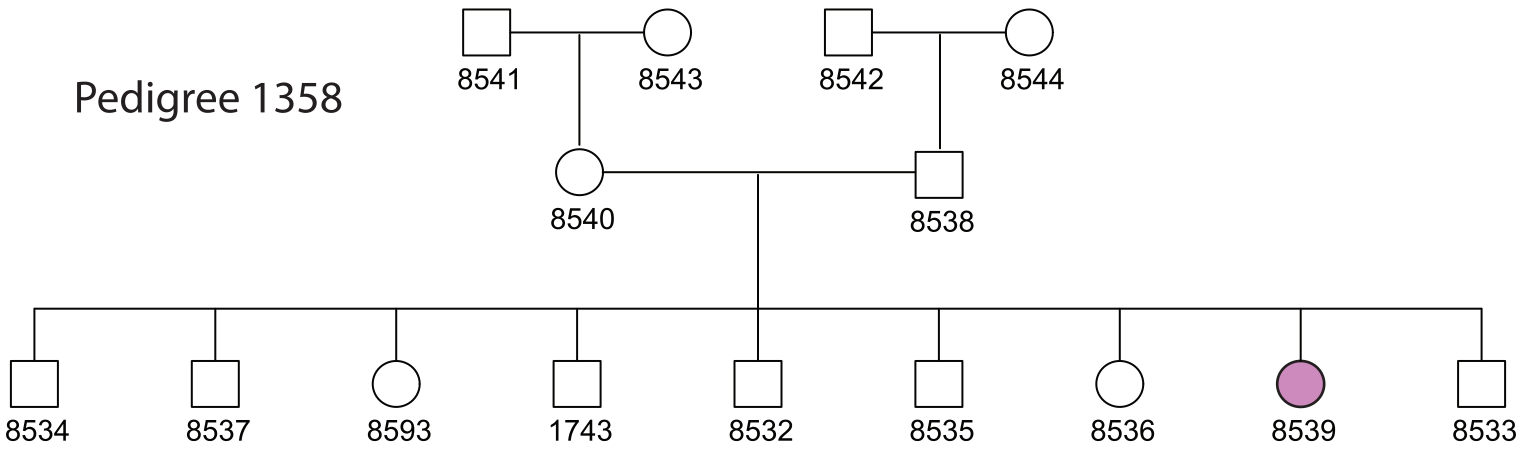
Pedigree 1358
8541
8543
8542
8544
8540
8538
8534
8537
8593
1743
8532
8535
8536
8539
8533

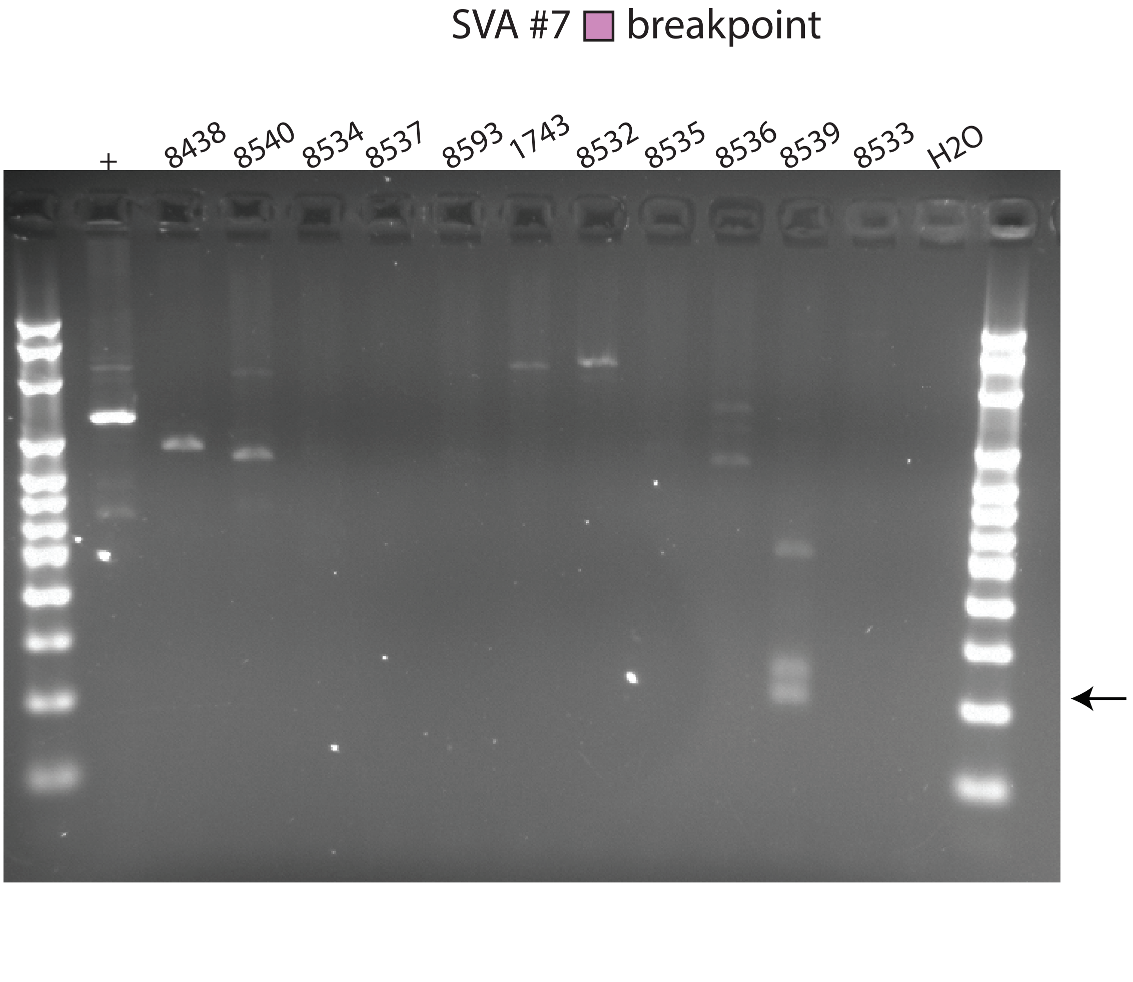
SVA #7 breakpoint
+
8438
8540
8534
8537
8593
1743
8532
8535
8536
8539
8533
H2O

I

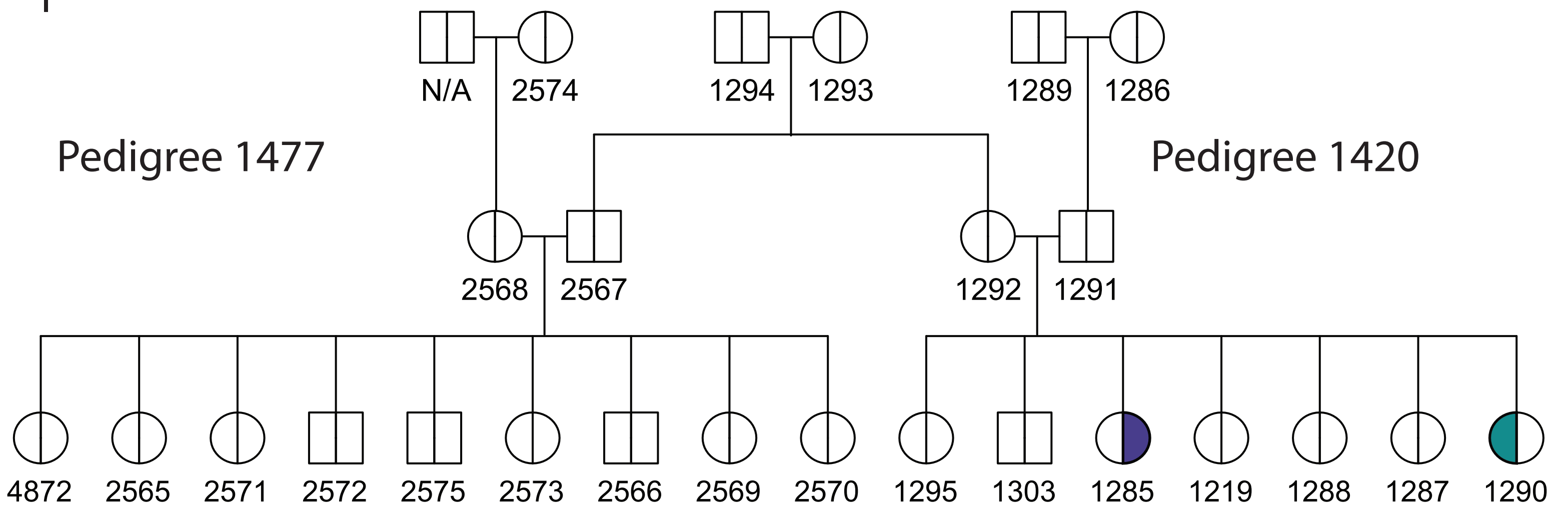
N/A
2574
1294
1293
1289
1286
Pedigree 1477
Pedigree 1420
2568
2567
1292
1291
4872
2565
2571
2572
2575
2573
2566
2569
2570
1295
1303
1285
1219
1288
1287
1290

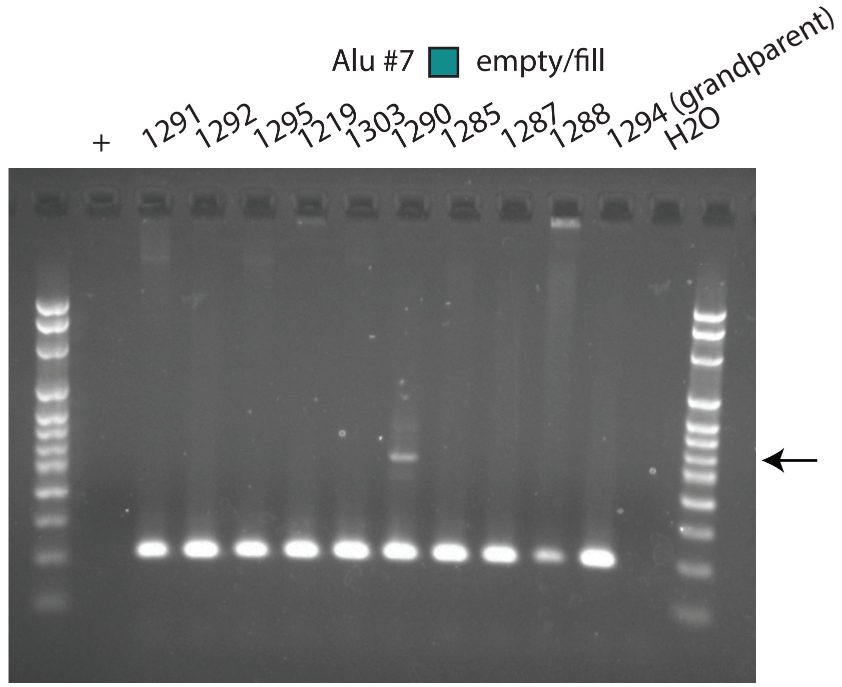
Alu #7 empty/fill
+
1291
1292
1295
1219
1303
1290
1285
1287
1288
1294 (grandparent)
H2O

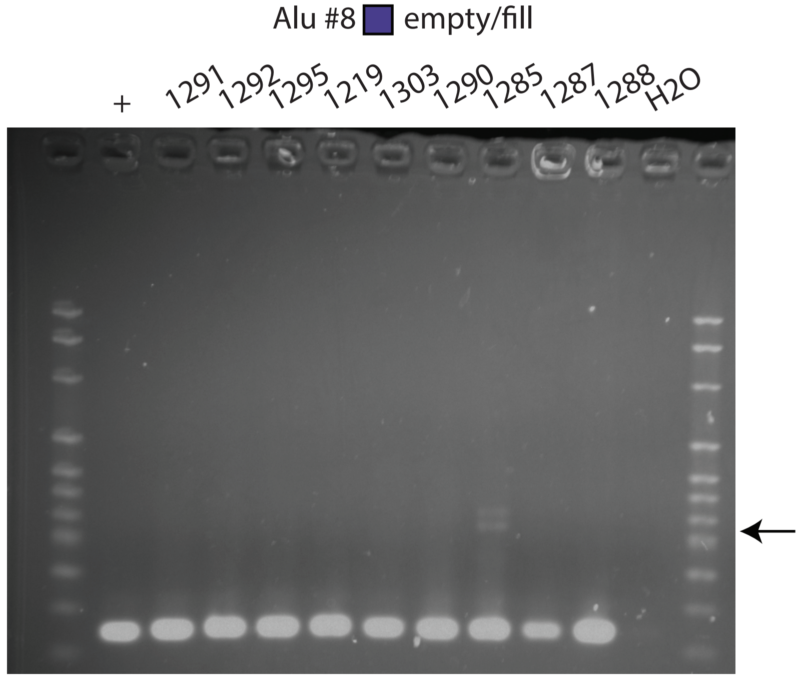
Alu #8 empty/fill
+
1291
1292
1295
1219
1303
1290
1285
1287
1288
H2O

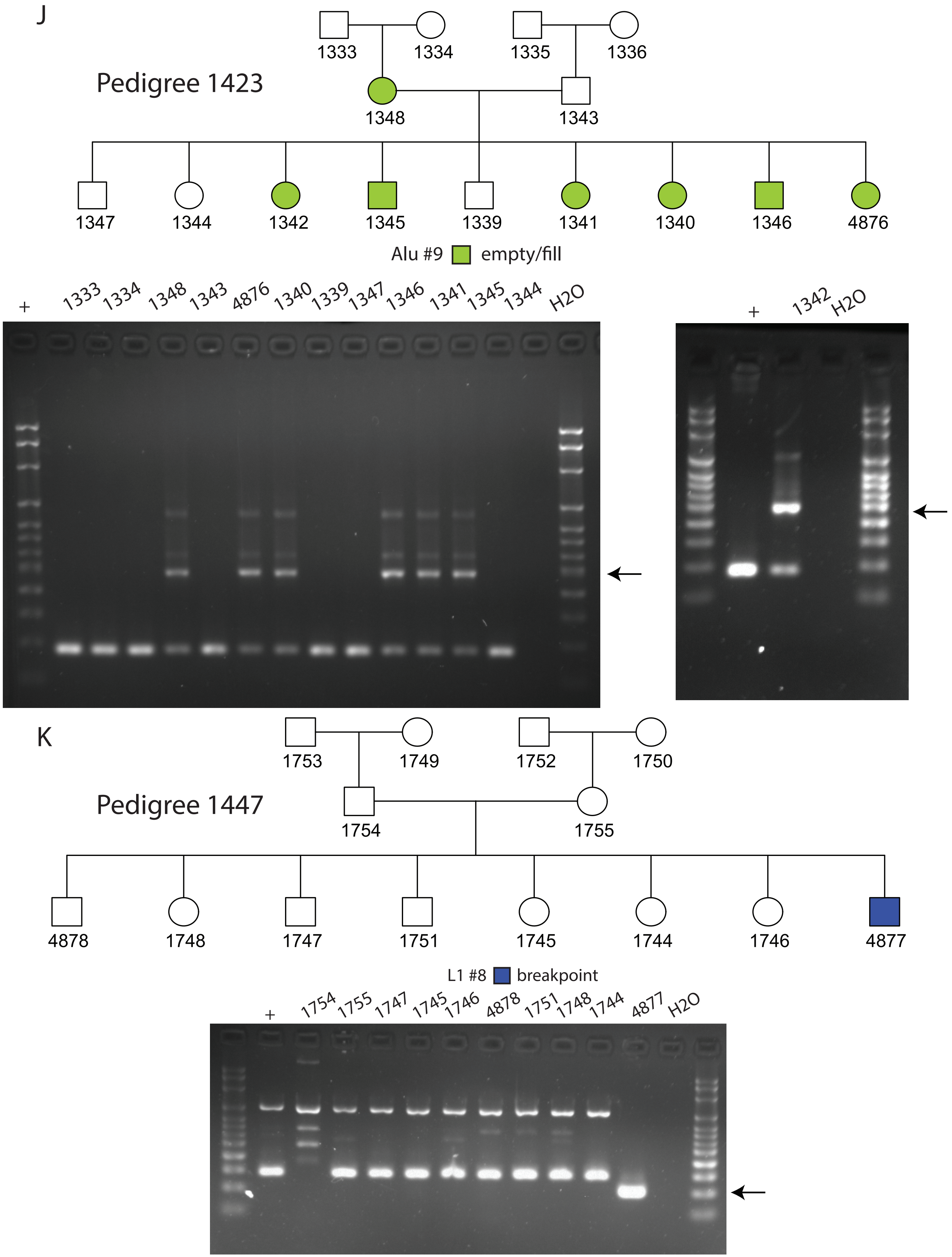
J
Pedigree 1423
1333
1334
1335
1336
1348
1343
1347
1344
1342
1345
1339
1341
1340
1346
4876
Alu #9 empty/fill
+ 1333 1334 1348 1343 4876 1340 1339 1347 1346 1341 1345 1344 H2O
+ 1342 H2O
K
Pedigree 1447
1753
1749
1752
1750
1754
1755
4878
1748
1747
1751
1745
1744
1746
4877
L1 #8 breakpoint
+ 1754 1755 1747 1745 1746 4878 1751 1748 1744 4877 H2O

L

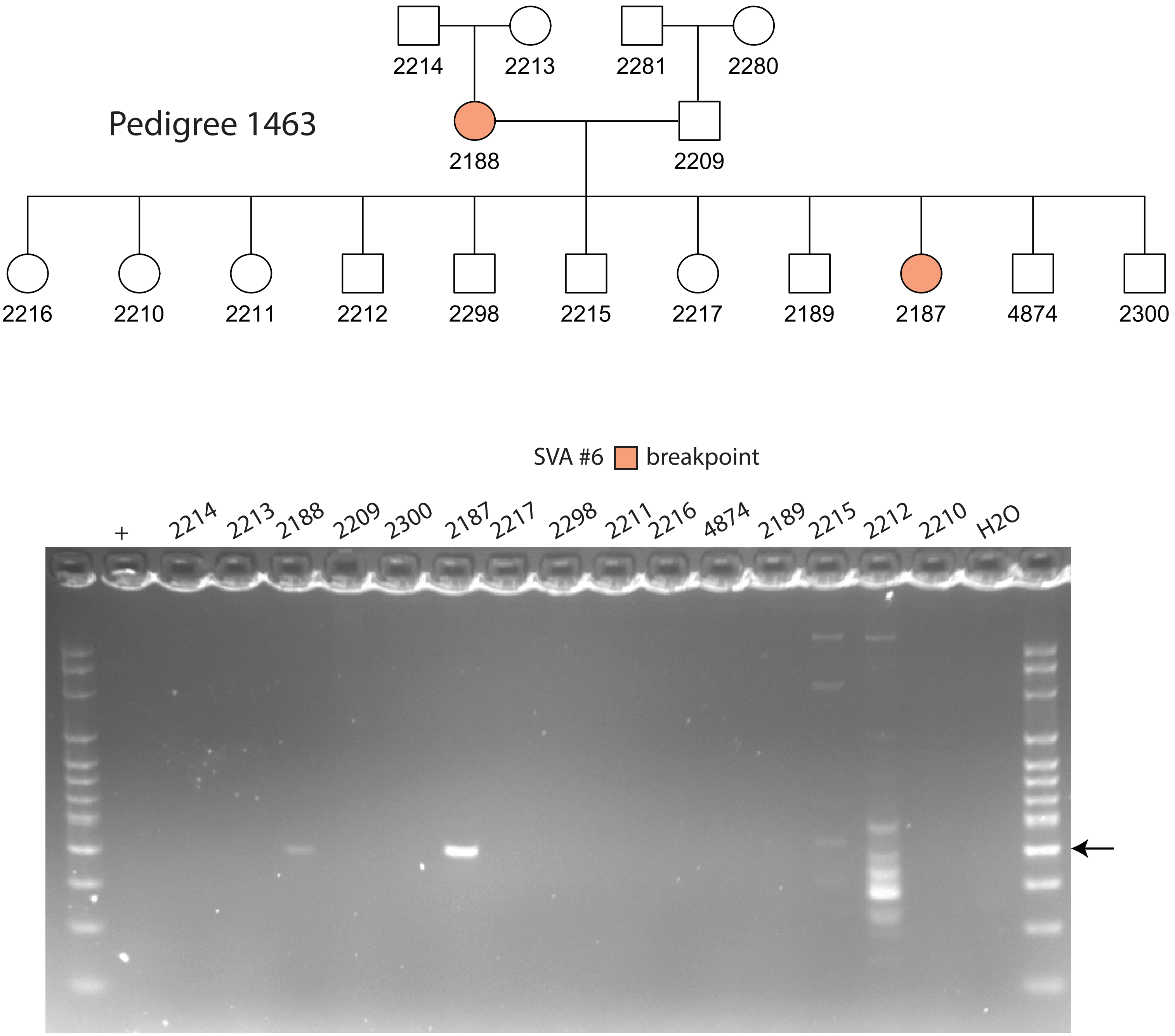
Pedigree 1463
2214
2213
2281
2280
2188
2209
2216
2210
2211
2212
2298
2215
2217
2189
2187
4874
2300
SVA #6
breakpoint
+
2214
2213
2188
2209
2300
2187
2217
2298
2211
2216
4874
2189
2215
2212
2210
H2O

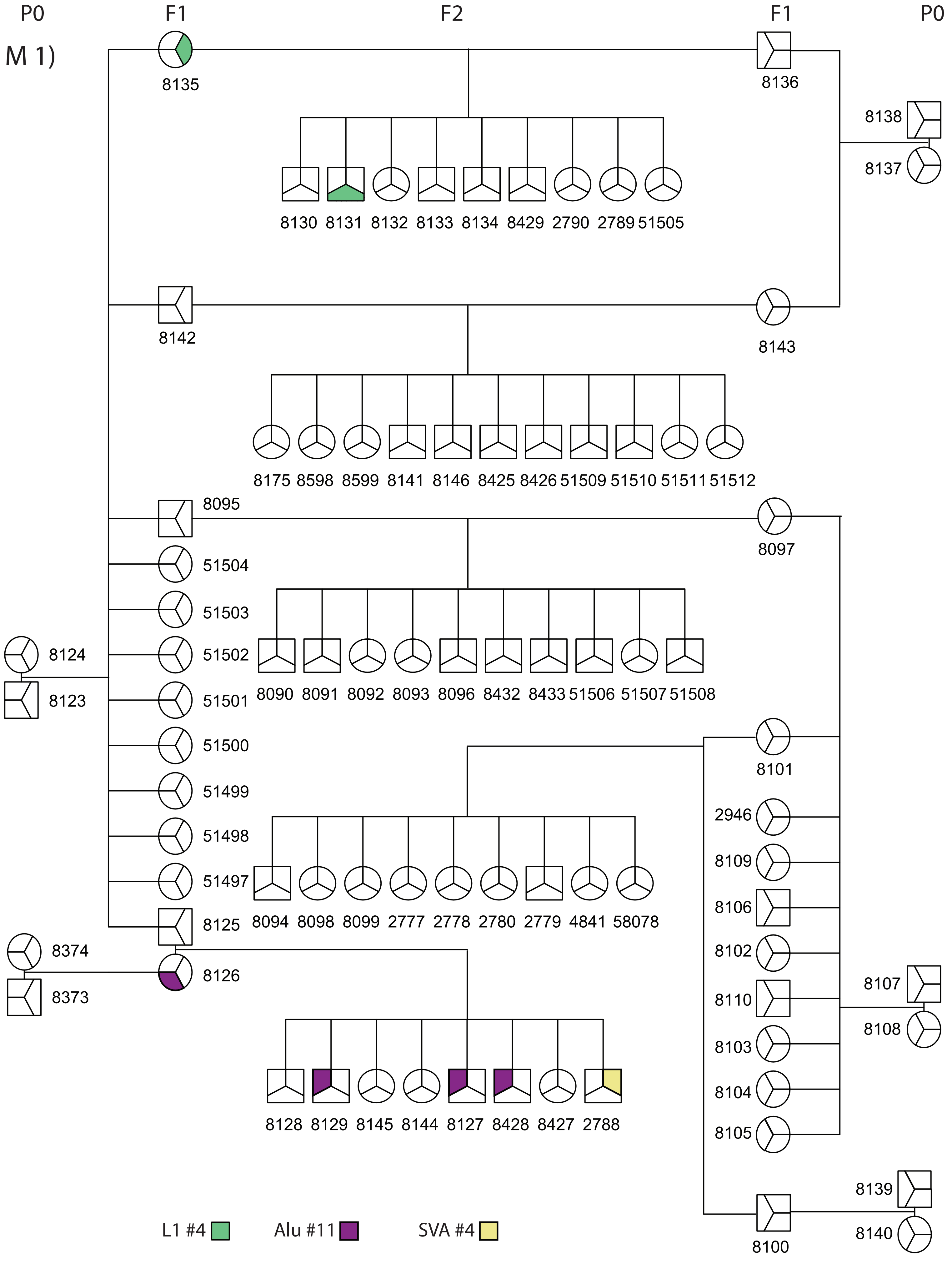

P0
F1
F2
F1
P0
M 1)
8135
8136
8138
8137
8130 8131 8132 8133 8134 8429 2790 2789 51505
8142
8143
8175 8598 8599 8141 8146 8425 8426 51509 51510 51511 51512
8095
8097
51504
51503
51502
51501
51500
51499
51498
51497
8125
8124
8123
8090 8091 8092 8093 8096 8432 8433 51506 51507 51508
8101
2946
8109
8106
8102
8110
8103
8104
8105
8107
8108
8094 8098 8099 2777 2778 2780 2779 4841 58078
8374
8373
8126
8128 8129 8145 8144 8127 8428 8427 2788
8139
8140
8100
L1 #4
Alu #11
SVA #4

M 2)

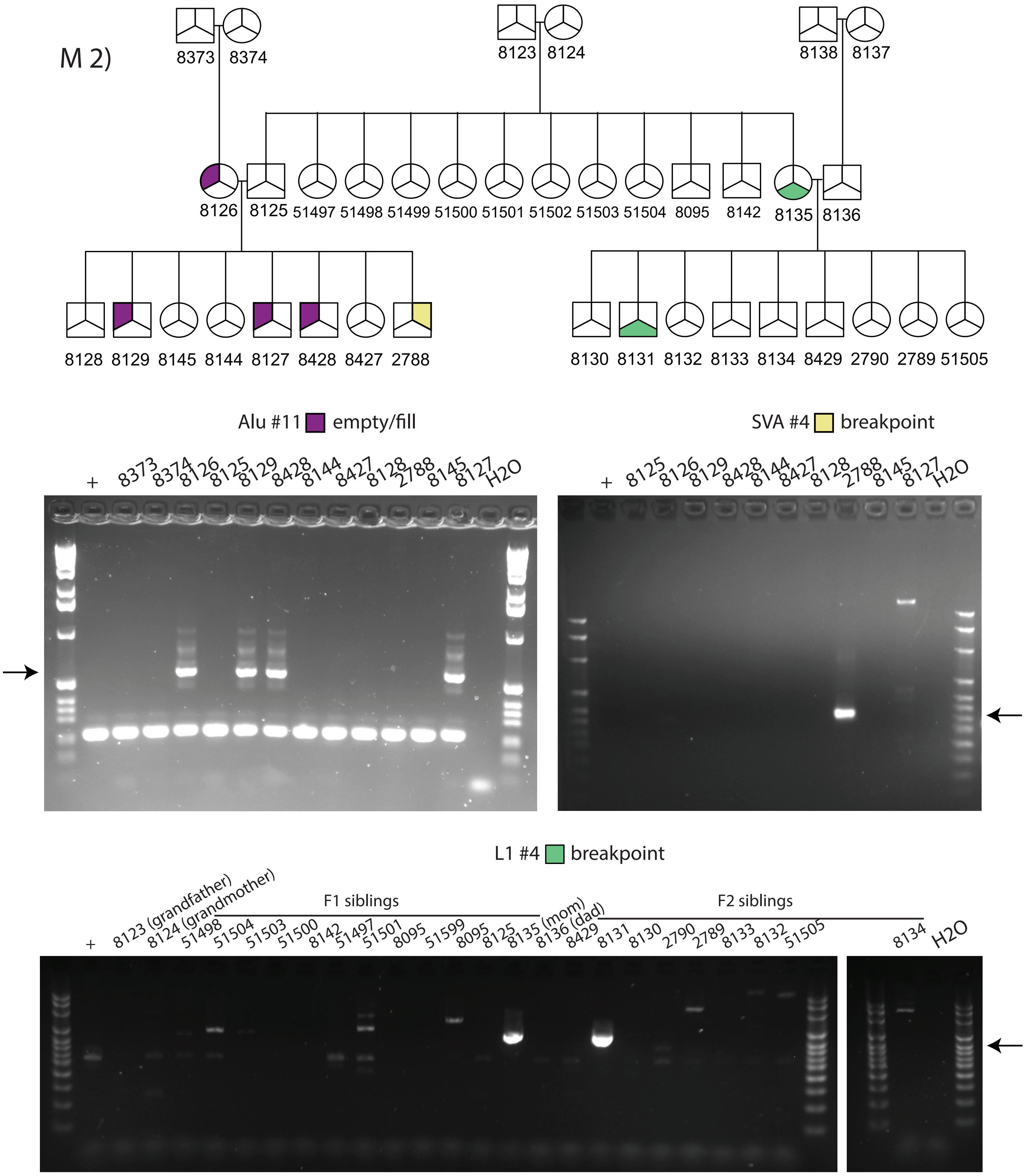
8373 8374
8123 8124
8138 8137
8126 8125 51497 51498 51499 51500 51501 51502 51503 51504 8095 8142 8135 8136
8128 8129 8145 8144 8127 8428 8427 2788
8130 8131 8132 8133 8134 8429 2790 2789 51505
Alu #11 empty/fill
+ 8373 8374 8126 8125 8129 8428 8144 8427 8128 2788 8145 8127 H2O
SVA #4 breakpoint
+ 8125 8126 8129 8428 8144 8427 8128 2788 8145 8127 H2O
L1 #4 breakpoint
F1 siblings
F2 siblings
+ 8123 (grandfather) 8124 (grandmother) 51498 51504 51503 51500 8142 51497 51501 8095 51599 8095 8125 8135 (mom) 8136 (dad) 8429 8131 8130 2790 2789 8133 8132 51505
8134 H2O

N

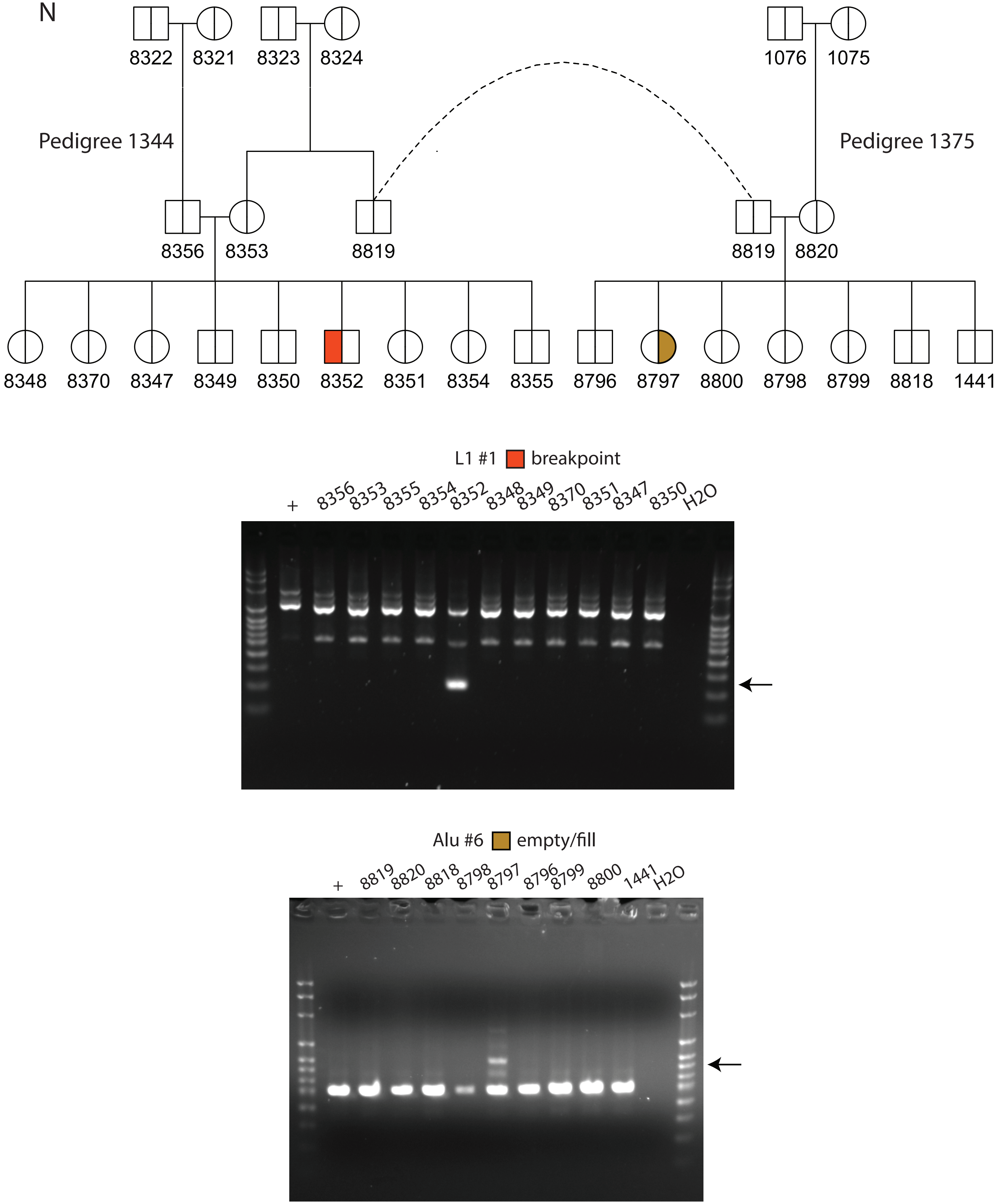

8322
8321
8323
8324
1076
1075
Pedigree 1344
Pedigree 1375
8356
8353
8819
8819
8820
8348
8370
8347
8349
8350
8352
8351
8354
8355
8796
8797
8800
8798
8799
8818
1441
L1 #1 breakpoint
+
8356
8353
8355
8354
8352
8348
8349
8370
8351
8347
8350
H2O
Alu #6 empty/fill
+
8819
8820
8818
8798
8797
8796
8799
8800
1441
H2O

O

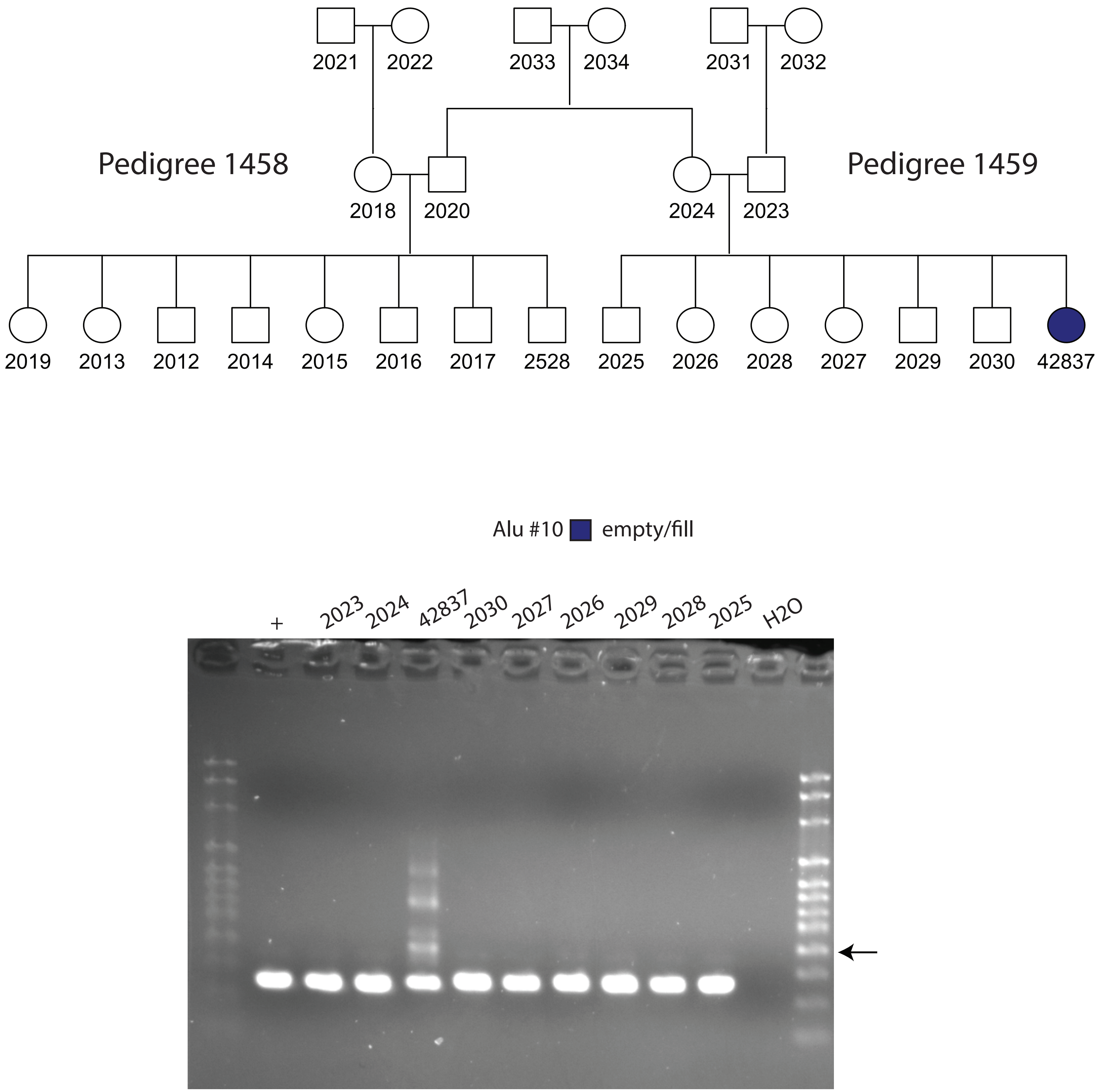
Pedigree 1458
Pedigree 1459
2021
2022
2033
2034
2031
2032
2018
2020
2024
2023
2019
2013
2012
2014
2015
2016
2017
2528
2025
2026
2028
2027
2029
2030
42837
Alu #10 empty/fill
+
2023
2024
42837
2030
2027
2026
2029
2028
2025
H2O

P

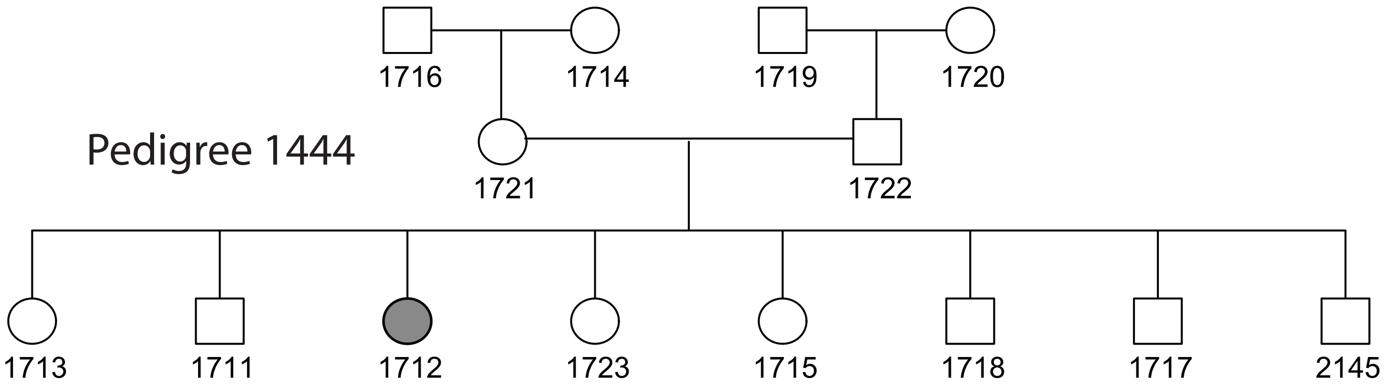


SVA #8 breakpoint

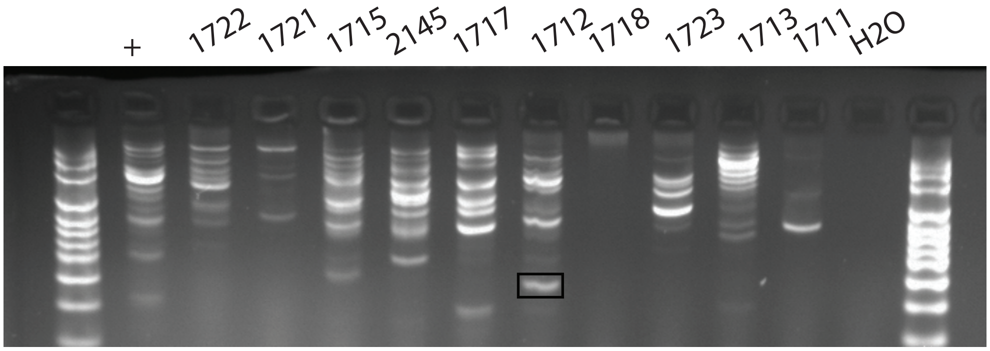

**Supplemental Figure S1**. Pedigree structure and PCR of the 27 *de novo* MEI. Each PCR reaction was run with a non-pedigree individual control DNA ("+") and H2O. Empty/fill refers to amplifying around the MEI, including amplification of the WT chromosome. Breakpoint refers to amplification using one internal primer and one primer outside of the MEI. The black arrow indicates the MEI band.

A) Pedigree 1331 and Alu #1-2, L1 #5, #7, and SVA #1, #5. Amplification of L1 #7 leads to non-specific bands of the same size as the amplicon; therefore, a full pedigree gel is not provided.
B) Pedigree 1341 and Alu #3.
C) Pedigree 1345 and Alu #4. 1) UCSC genome browser screenshot of the MEI insertion. This Alu caused a 1.7kb 5' deletion. The three primer sets are labeled in red (Alu primer set), black (SNP primer set) and purple (Alu + SNP primer set). 2) Pedigree diagram. 3) Gel of amplification of Alu #4 using the Alu primer set. The Alu band amplifies faintly in individual 8439. 4) Sanger sequence of 8327, 8439, and their parents at this locus. When the locus is amplified around a nearby SNP, 8327 is heterozygous G/C and 8439 is homozygous G/G. When the MEI insertion and SNP are amplified together (Alu + SNP primer set), the C allele is on the same band as the Alu band, indicating that the Alu landed on the maternal chromosome. Individual 8439 has a C in the Alu band. Given the haplotype discrepancy and lack of recombination evidence, we suspect that the 8439 DNA has low-level contamination of individual 8327.
D) Pedigree 1346 and Alu #5.
E) Pedigree 1347 and L1 #2 and #6.
F) Pedigree 1353 and SVA #2.
G) Pedigree 1354 and L1 #3.
H) Pedigree 1358 and SVA #7.
I) Pedigree 1420/1477 and Alu #7-8.
J) Pedigree 1423 and Alu #9.
K) Pedigree 1447 and L1 #8.
L) Pedigree 1463 and SVA #6.
M) Pedigree 1328/1329 and Alu #11, L1 #4, and SVA #4. 1) Full diagram of the pedigree. 2) Subset of pedigree and gel of the three MEI.
N) Pedigree of 1344/1375 and Alu #6 and L1 #1.
O) Pedigree of 1458/1459 and Alu #10.
P) Pedigree of 1444 and SVA #8

### Alu #1
### chr4q24

Empty site:

1st strand cleavage

5'-ATGCAAAAAGAAAGAAACTGCTTTTTATTATAAGAAATAACTCAA-3'
3'-TAGCTTTTTCTTTCTTTGACGAAAAATAATATTCTTTATTGAGTT-5'

Filled site:

5'-ATGCAAAAAGAAAGAAACTGCTTTTTT$_{63}$ AluYb8 GAAACTGCTTTTTATTATAAGAAATAACTCAA-3'
3'-TAGCTTTTTCTTTCTTTGACGAAAAAA$_{63}$ CTTTGACGAAAAATAATATTCTTTATTGAGTT-5'

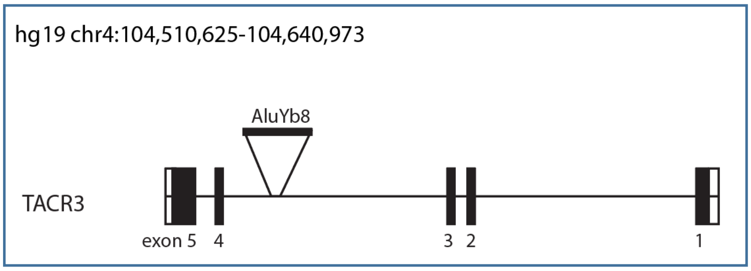


- strand
TSD: 13bp
chr4:104,530,552-104,530,567

### Alu #2
### chr1p36.22

Empty site:

5'-AAGACTCCATCTCAAAAATAGAAACAAAAACAAACAAA-3'
3'-TTCTGAGGTAGAGTTTTTATCTTTGTTTTTGTTTGTTT-5'

1st strand cleavage

Filled site:

5'-AAGACTCCATCTCAAAAATAGAAACA AluYb8 A$_{44}$AAAAAATAGAAACAAAAACAAACAAA-3'
3'-TTCTGAGGTAGAGTTTTTATCTTTGT T$_{44}$TTTTTTATCTTTGTTTTTGTTTGTTT-5'

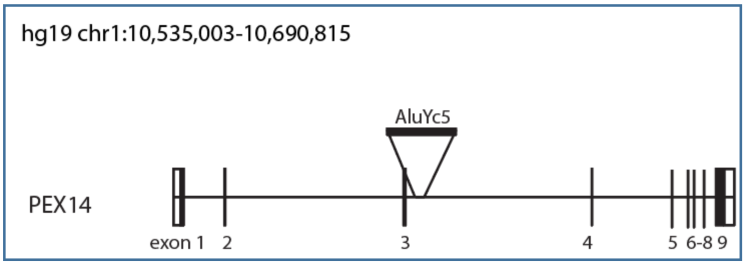


+ strand
TSD: 13
chr1:10,602,818-10,602,830

### Alu #3
### chr3p24.2

Empty site:

5′-TTCTACACATTTATTAAGATTTATCTATTGAACAGTCAACTCCT-3′
3′-AAGATGTGTAAATAATTCTAAATAGATAACTTGTCAGTTGAGGA-5′

$1^{st}$ strand cleavage

Filled site:

5′-TTCTACACATTTATTAAGATTTATCT AluY $A_{80}$AAGATTTATCTATTGAACAGTCAACTCCT-3′
3′-AAGATGTGTAAATAATTCTAAATAGA $T_{80}$TTCTAAATAGATAACTTGTCAGTTGAGGA-5′

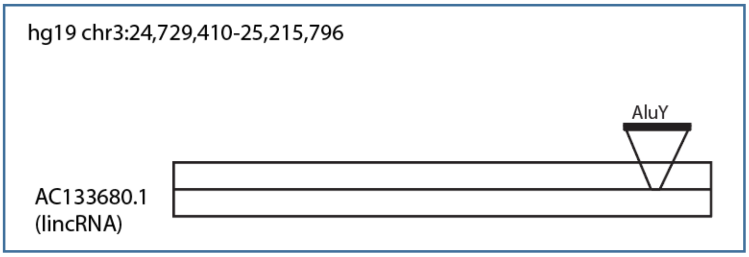


+ strand
TSD: 11
chr3:25,163,270-25,163,280

### Alu #4
### chr12p13.31

Empty site:

deleted

5′-GATTAATGGGTTGATGTATTA...1.7kb...TTTTGATAGAAATAGAAACATGGAACTCAATAGAAA-3′
3′-CTAATTACCCAACTACATAAT...1.7kb...AAAACTATCTTTATCTTTGTACCTTGAGTTATCTTT-5′

$1^{st}$ strand cleavage

Filled site:

5′-GATTAATGGGTTGATGTATT**T** AluYa2 $A_{41}$AGAAACATGGAACTCAATAGAAA-3′
3′-CTAATTACCCAACTACATAA**A** $T_{41}$TCTTTGTACCTTGAGTTATCTTT-5′

+ strand
TSD: 5′ deletion
5′ end: chr12:8,680,801-8,680,820
3′ end: chr12:8,682,497-8,682,519

Intergenic

### Alu #5
### chr6p22.2

Empty site:

5′-AATATACATGTAATAAAAAAAAATCACAAGGTGTGATATACATGTA-3′
3′-TTATATGTACATTATTTTTTTTTAGTGTTCCACACTATATGTACAT-5′

1st strand cleavage

Filled site:

5′-AATATACATGTAATAAAAAAAAATCAC AluYb8 $A_{66}$AAAAAAAAATCACAAGGTGTGATATACATGTA-3′
3′-TTATATGTACATTATTTTTTTTTAGTG $T_{66}$AAAAAAAAATCACAAGGTGTGATATACATGTA-5′

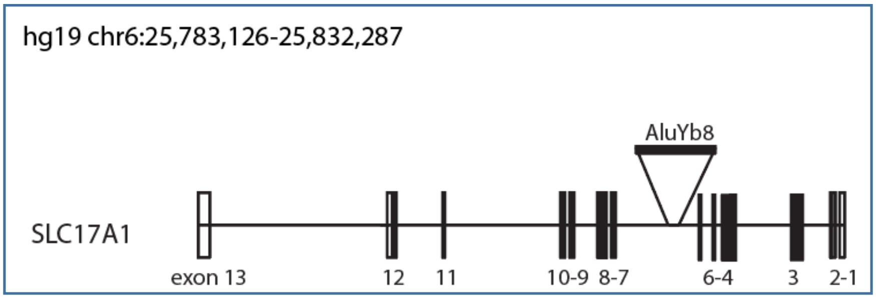


+ strand
TSD: 12
chr6:25,815,318-25,815,329

### Alu #6
### chr2p13.1

Empty site:

5′-AAGAGCTTGTTTTAAAAAATTAATTCGTTAATTAAAAAAAAAT-3′
3′-TTCTCGAACAAAATTTTTTAATTAAGCAATTAATTTTTTTTTA-5′

1st strand cleavage

Filled site:

5′-AAGAGCTTGTTTTAAAAAATTAATTCGTT AluYc1 $A_{24}$AAAAAATTAATTCGTTAATTAAAAAAAAAT-3′
3′-TTCTCGAACAAAATTTTTTAATTAAGCAA $T_{24}$TTTTTTAATTAAGCAATTAATTTTTTTTTA-5′

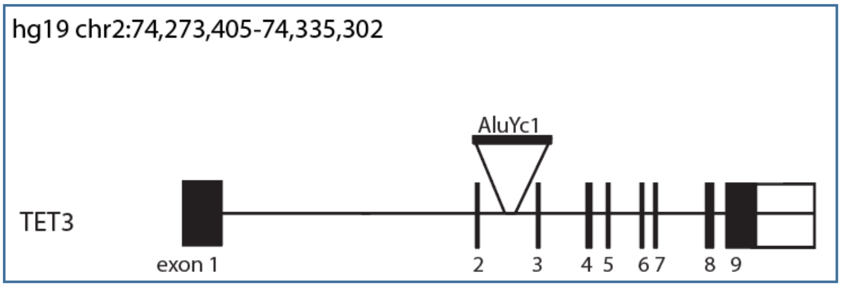


+ strand
TSD: 16
chr2:74,305,312-74,305,327

### Alu #7
### chr1p13.3

Empty site:

5’-TAAAAATAAACAAAAATAAGAAATAAAATCAGCCATAATCCTACT-3’
3’-ATTTTTATTTGTTTTTATTCTTTATTTTAGTCGGTATTAGGATGA-5’

1st strand cleavage

Filled site:

5’-TAAAAATAAACAAAAATAAGAAATAAAATCAG AluYe5 $A_{47}$AAGAAATAAAATCAGCCATAATCCTACT-3’
3’-ATTTTTATTTGTTTTTATTCTTTATTTTAGTC $T_{47}$TTCTTTATTTTAGTCGGTATTAGGATGA-5’

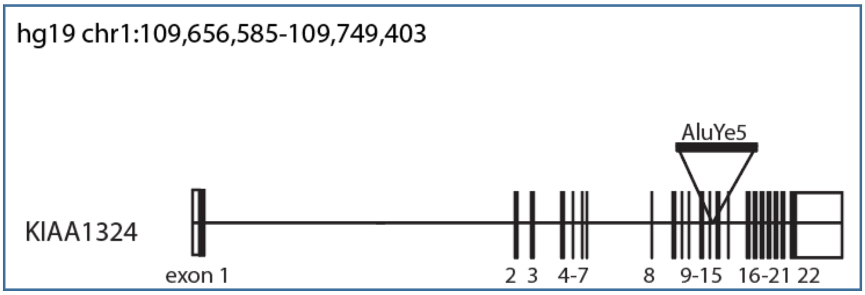


+ strand
TSD: 15
chr1:109,734,843-109,734,857

### Alu #8
### chr2q22.2

Empty site:

1st strand cleavage

5’-TTGGAACACCCAGATGTATAAAACAAATATTATTAGATCTAAAG-3’
3’-AACCTTGTGGGTCTACATATTTTGTTTATAATAATCTAGATTTC-5’

Filled site:

5’-TTGGAACACCCAGATGTATAAAACAAATATT$T_{113}$ AluYa5 ATAAAACAAATATTATTAGATCTAAAG-3’
3’-AACCTTGTGGGTCTACATATTTTGTTTATAA$A_{113}$ TATTTTGTTTATAATAATCTAGATTTC-5’

- strand
TSD: 14
chr2:143,161,204-143,161,217

Intergenic

### Alu #9
### chr7q31.1

Empty site:

5’-TAGAGTCAAGTTCCATCAAATCAGGAGGCACAAAATATCGGTGC-3’
3’-ATCTCAGTTCAAGGTAGTTTAGTCCTCCGTGTTTTATAGCCACG-5’

1st strand cleavage

Filled site:

5’-TAGAGTCAAGTTCCATCAAATCAGGAGGCAC AluYa5 $A_{107}$AAATCAGGAGGCACAAAATATCGGTGC-3’
3’-ATCTCAGTTCAAGGTAGTTTAGTCCTCCGTG AluYa5 $T_{107}$TTTAGTCCTCCGTGTTTTATAGCCACG-5’

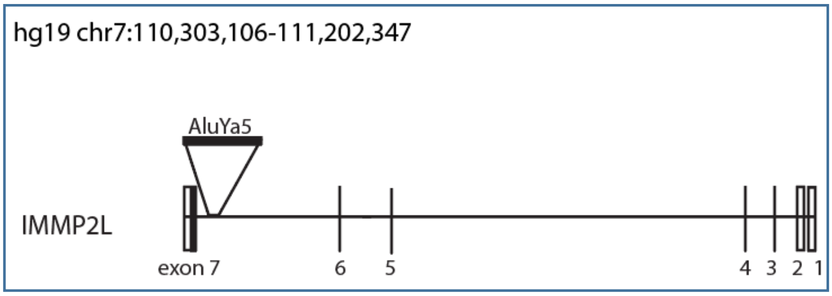


+ strand
TSD: 14
chr7:110,305,748-110,305,761

### Alu #10
### chr4q22.3

Empty site:

1st strand cleavage

5’-TTTTCCTGCTTTATTCTAAGTCATTTTAACTATATTTATTACAT-3’
3’-AAAAGGACGAAATAAGATTCAGTAAAATTGATATAAATAATGTC-5’

Filled site:

5’-TTTTCCTGCTTTATTCTAAGTCATTTT$T_{113}$ AluY TAAGTCATTTTAACTATATTTATTACAT-3’
3’-AAAAGGACGAAATAAGATTCAGTAAAA$A_{113}$ AluY ATTCAGTAAAATTGATATAAATAATGTC-5’

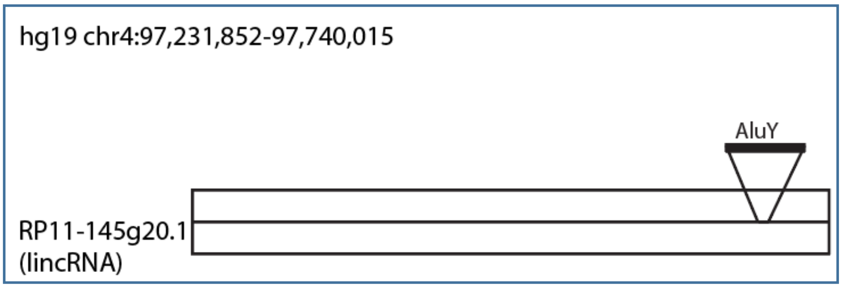


- strand
TSD: 11
chr4:97,701,273-97,701,283

### Alu #11
### chr17p11.2

Empty site:

1$^{st}$ strand cleavage

5’-ACCTACTATGTACCCATCAGATTTTTTAAAAAAGAAATTAA-3’
3’-TGGATGATACATGGGTAGTCTAAAAAATTTTTTCTTTAATT-5’

Filled site:

5’-ACCTACTATGTACCCATCAGATTTTTTT$_{113}$ AluYa5 CCATCAGATTTTTTAAAAAAGAAATTAA-3’
3’-TGGATGATACATGGGTAGTCTAAAAAAA$_{113}$ GGTAGTCTAAAAAATTTTTTCTTTAATT-5’

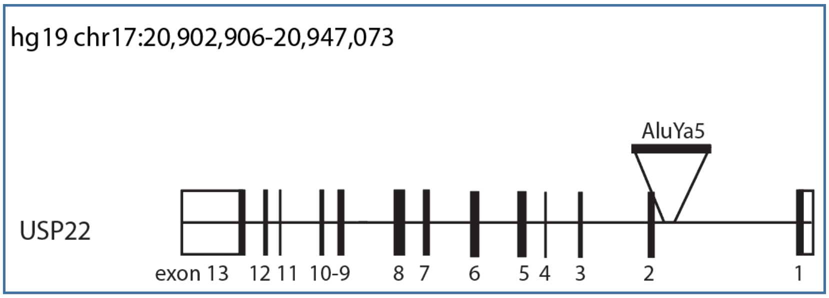


- strand
TSD: 14
chr17:20,932,724-20,932,737

# L1 #1
### chr5q15

Empty site:

double-strand break

5’-ATTTAAAATTAGTTTGAATCCTCTGCTTGTTCCTGATTAAA-3’
3’-TAAATTTTAATCAAACTTAGGAGACGAACAAGGACTAATTT-5’

Filled site:

L1 microhomology

5’-ATTTAAAATTAGTTTGA partial ORF2 TCCTCTGCTTGTTCCTGATTAAA-3’
3’-TAAATTTTAATCAAACT AGGAGACGAACAAGGACTAATTT-5’

- strand
1bp deletion
chr5:95,081,487

# L1 #2
### chr4q28.2

Empty site:

5′-TGCACTTAACAGTATCATGTAAGTATCAGTTTCT-3′
3′-ACGTGAATTGTCATAGTACATTCATAGTCAAAGA-5′

1st strand cleavage

Filled site:

5′-TGCACTTAACAGTATCATGTAA | UTR | ORF1 | ORF2/UTR | $A_{29}$ | 82 bp | $A_{96}$AACAGTATCATGTAAGTATCAGTTTCT-3′
3′-ACGTGAATTGTCATAGTACATT | | | | $T_{29}$ | | $T_{96}$TTGTCATAGTACATTCATAGTCAAAGA-5′

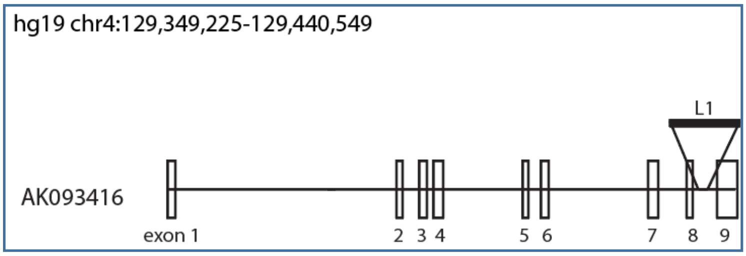


+ strand
TSD: 15
chr4:129,438,004-129,438,018
Transduced sequence:
chr4:112,628,960-112,629,041

# L1 #3
### chr3p14.3

Empty site:

1st strand cleavage

5′TTTTGGGTTTTTTGTATGTGTGTTTTTTTTAAAAAAAGAAACAG-3′
3′AAAACCCAAAAAACATACACACAAAAAAAATTTTTTTCTTTGTC-5′

Filled site:

5′-TTTTGGGTTTTTTGTATGTGTGTTTTTTTTT$_{??}$ | ORF2/UTR | GTATGTGTGTTTTTTTTAAAAAAAGAAACAG-3′
3′-AAAACCCAAAAAACATACACACAAAAAAAAA$_{??}$ | | CATACACACAAAAAAAATTTTTTTCTTTGTC-5′

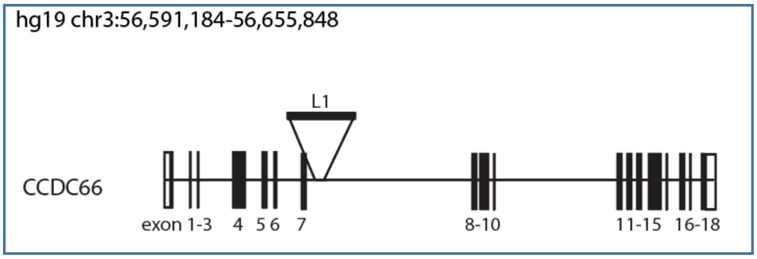


- strand
TSD: 17
chr3:56,605,741-56,605,757

# L1 #4
### chr1p31.2

Empty site:

5'-TCTCGTGTAGTTGTAAGAGGAGCAGCTTTGATAATTGTTC-3'
3'-AGAGCACATCAACATTCTCCTCGTCGAAACTATTAACAAG-5'

1st strand cleavage

Filled site:

5'-TCTCGTGTAGTTGTAAGAGGAGCAGCT [ORF2/UTR] $A_{32}$ [846 bp] $A_{94}$AAGAGGAGCAGCTTTGATAATTGTTC-3'
3'-AGAGCACATCAACATTCTCCTCGTCGA [ORF2/UTR] $T_{32}$ [846 bp] $T_{94}$TTCTCCTCGTCGAAACTATTAACAAG-5'

+ strand
TSD: 13
chr1:69,238,322-69,238,334
Transduced sequence:
chr5:112,703,068-112,703,913

Intergenic

# L1 #5
### chr4q28.2

Empty site:

1st strand cleavage

5'-TTTGAAAACAGAAAAGCA...CAATATATTTTAAAAAGAAAGA-3'
3'-AAACTTTTGTCTTTTCGT...GTTATATAAAATTTTTCTTTCT-5'

Filled site:

5'-TTTGAAAACAGAAAAGCA...CAATATATTTTAT$_{83}$ [UTR / ORF1] AACAGAAAAGCA...CAATATATTTTAAAAAGAAAGA-3'
3'-AAACTTTTGTCTTTTCGT...GTTATATAAAATA$_{83}$ [ORF2 / ORF1] TTGTCTTTTCGT...GTTATATAAAATTTTTCTTTCT-5'

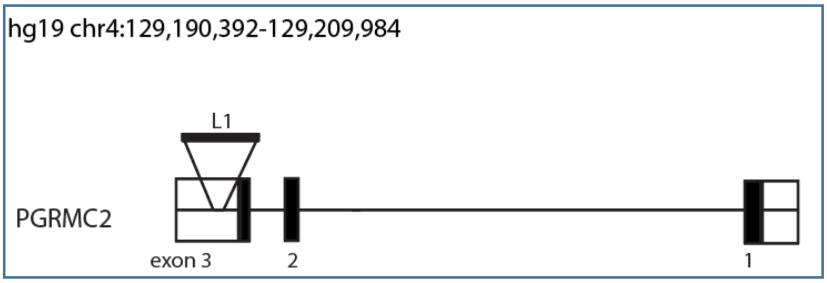


- strand
TSD: 628
chr4:129,191,510-129,192,137

# L1 #6
### chrXq13.1

Empty site:

5’-GAAGACGAAGTTGAAACTTAATAACTTGCCTACGGGCACAGGCTGTTATA -3’
3’-CTTCTGCTTCAACTTTGAATTATTGAACGGATGCCCGTGTCCGACAATAT-5’

$1^{st}$ strand cleavage

Filled site:

5’-GAAGACGAAGTTGAAACTTAATAACTT [497bp] $A_{59}$AATAACTTGCCTACGGGCACAGGCTGTTATA-3’
3’-CTTCTGCTTCAACTTTGAATTATTGAA $T_{59}$TTATTGAACGGATGCCCGTGTCCGACAATAT-5’

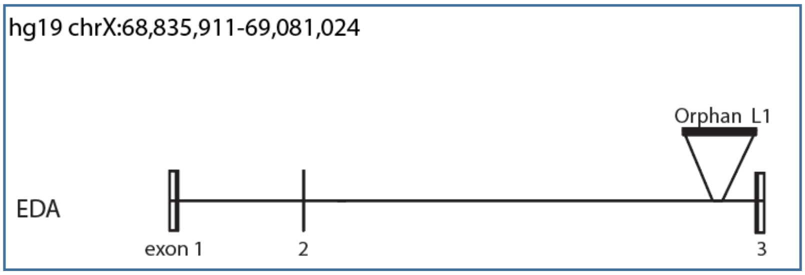


+ strand
TSD: 8
chrX:68,964,693-68,964,700
Orphan Transduction
chr13:61,460,432-61,460,928

# L1 #7
### chr6q15

Empty site:

**exonic**

5’-**GCACA**GTAAGAACTTTTAAAAGTTAATCTAAGTTACAAT-3’
3’-**CGTGT**CATTCTTGAAAATTTTCAATTAGATTCAATGTTA-5’

$1^{st}$ strand cleavage

Filled site:

5’-**GCACA**GTAAGAACTTTTAAAAGTTAATCTA [UTR | ORF1 | ORF2/UTR] $A_{XX}$AAAAGTTAATCTAAGTTACAAT-3’
3’-**CGTGT**CATTCTTGAAAATTTTCAATTAGAT $T_{XX}$TTTTCAATTAGATTCAATGTTA-5’

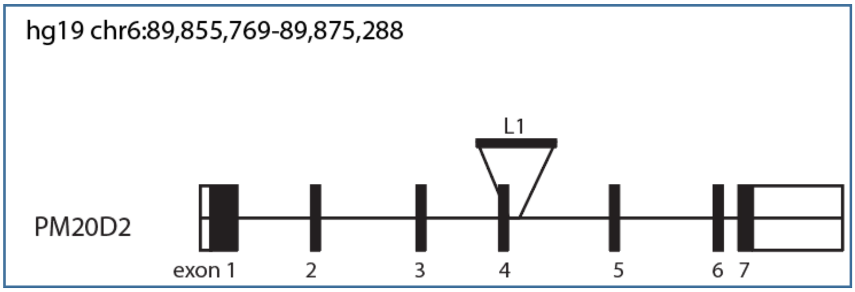


+ strand
TSD: 13
chr6:89,864,634-89,864,646

# L1 #8
### chrYq11.21

Empty site:

1st strand cleavage

5'-ATGTTTATTACCCACTTGTGTTAGTTTCTTAAGGCTGCTGTGACAAATCA-3'
3'-TACAAATAATGGGTGAACACAATCAAAGAATTCCGACGACACTGTTTAGT-5'

Filled site:

5'-ATGTTTATTACCCACTTGTGTTAGTTTCTTTT?? | ORF2/UTR | TGTTAGTTTCTTAAGGCTGCTGTGACAAATCA-3'
3'-TACAAATAATGGGTGAACACAATCAAAGAAAT?? | | ACAATCAAAGAATTCCGACGACACTGTTTAGT-5'

Intergenic

- strand
TSD: 12
chrY:13,871,221-13,871,232

### SVA #1
### chr6p22.1

Empty site:

1st strand cleavage

5'-GCAATCTGATTTGGCAAATATTTTTAAAGATGATATTTGAAT-3'
3'-CGTTAGACTAAACCGTTTATAAAAATTTCTACTATAAACTTA-5'

Filled site:

19bp upstream

5'-GCAATCTGATTTGGCAAATATTTTTT74 | SINE_R | VNTR | Alu | | GGCAAATATTTTTAAAGATGATATTTGAAT-3'
3'-CGTTAGACTAAACCGTTTATAAAAAA74 | | | | | GCGTTTATAAAAATTTCTACTATAAACTTA-5'

SVA_D

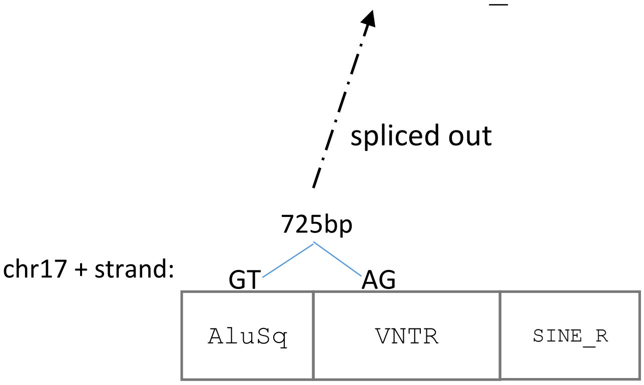


- strand
TSD: 13
chr6:29,684,013-29,684,026
Source SVA:
chr17: 42,314,401-42,316,970

Intergenic

### SVA #2
### chr4q31.1

Empty site:

5’-GCTATGCTTAAGAAAACAAAGCTGCATTTGGGAGGCTG-3’
3’-CGATACGAATTCTTTTGTTTCGACGTAAACCCTCCGAC-5’

1st strand cleavage

Filled site:

hexamer repeats

5’-GCTATGCTTAAGAAAACAAAGCTGCAT | AluSq | VNTR | SINE_R | $A_{45}$AAGAAAACAAAGCTGCATTTGGGAGGCTG-3’
3’-CGATACGAATTCTTTTGTTTCGACGTA | | | | $T_{45}$TTCTTTTGTTTCGACGTAAACCCTCCGAC-5’

SVA_E

+ strand
TSD: 18
chr4:140408752-140408769

Intergenic

### SVA #3
### chr4p11

Empty site:

1st strand cleavage

5’-CATAGTATTCCTGGAATACTATATTTTCAATTCTTCTCTTTAG-3’
3’-GTATCATAAGGACCTTATGATATAAAAGTTAAGAAGAGAAATC-5’

Filled site:

5’-CATAGTATTCCTGGAATACTATATTTTCT$_{??}$ | SINE_R | VNTR | CTGGAATACTATATTTTCAATTCTTCTCTTTAG-3’
3’-GTATCATAAGGACCTTATGATATAAAAGA$_{??}$ | | | GACCTTATGATATAAAAGTTAAGAAGAGAAATC-5’

SVA_F

- strand
TSD: 18
chr4:48,828,529-48,828,546

Intergenic

### SVA #4
### chr11p15.4

Empty site:

5’-CCTATCTATCTATCTAAAACTGAGGGTCAAAAATAGCCTCCA-3’
3’-GGATAGATAGATAGATTTTGACTCCCAGTTTTTATCGGAGGT-5’

↑
1st strand cleavage

Filled site:

5’-CCTATCTATCTATCTAAAACTGAGGGTC | MAST2 | VNTR | SINE_R | $A_{115}$TAAAACTGAGGGTCAAAAATAGCCTCCA-3’
3’-GGATAGATAGATAGATTTTGACTCCCAG | | | | $T_{115}$ATTTTGACTCCCAGTTTTTATCGGAGGT-5’

SVA_F1

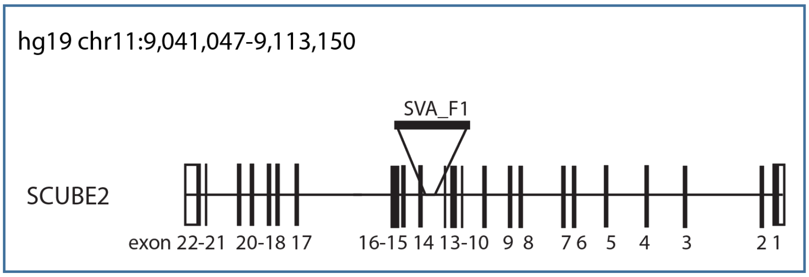


+ strand
TSD: 14
chr11:9,072,822-9,072,835

### SVA #5
### chr8p12

Empty site:

5’-GAAGATAAACTTAGAAAATTTTGTGTCACAAGGTATAT-3’
3’-CTTCTATTTGAATCTTTTAAAACACAGTGTTCCATATA-5’

↑
1st strand cleavage

Filled site:

479bp

5’-GAAGATAAACTTAGAAAATTTTGTGTCA | MAST2 | VNTR | SINE_R | AluSp | $A_{62}$AGAAAATTTTGTGTCACAAGGTATAT-3’
3’-CTTCTATTTGAATCTTTTAAAACACAGT | | | | | $T_{62}$TCTTTTAAAACACAGTGTTCCATATA-5’

SVA_F1

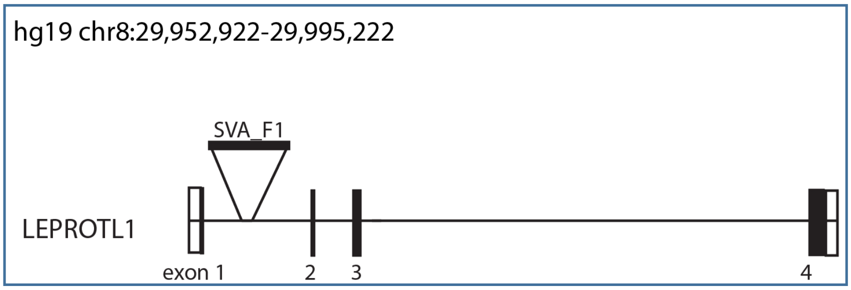


+ strand
TSD: 16
chr8:29,955,718-29,955,733

### SVA #6
### chr3p13

Empty site:

1st strand cleavage

5′-TGGAAAAAGAAGAAAATAGATTATGTTTCAATAAACCCATAAGGA-3′
3′-ACCTTTTTCTTCTTTTATCTAATACAAAGTTATTTGGGTATTCCT-5′

Filled site:

22bp deletion

5′-GAAAAAGAAGAAAATAGATTATGTTTCT$_{40}$ | SINE_R | VNTR | MAST2 | AluSc | GATTATGTTTCAATAAACCCATAAGGA-3′
3′-CTTTTTCTTCTTTTATCTAATACAAAGA$_{40}$ | | | | | CTAATACAAAGTTATTTGGGTATTCCT-5′

SVA_F1 122bp

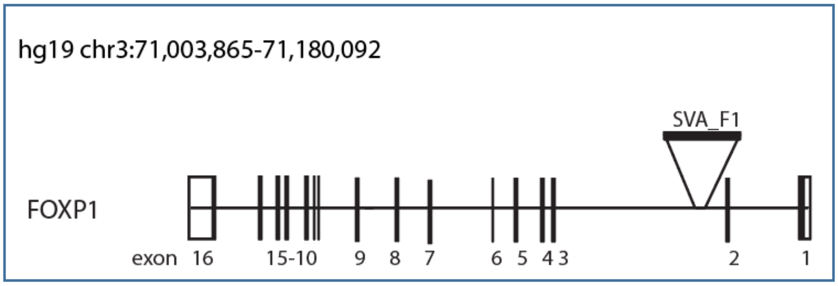


\- strand
TSD: 11
chr3:71,599,786-71,599,796
Source element:
chr3:48,251,893-48,254,907

### SVA #7
### chr12p13.2

Empty site:

5′-GAGCCTCTGTCTCAAAGAAAGAAAGAGTATGTACCTGGATATGTATTTTT -3′
3′-CTCGGAGACAGAGTTTCTTTCTTTCTCATACATGGACCTATACATAAAAA-5′

1st strand cleavage

Filled site:

5′-GAGCCTCTGTCTCAAAGAAAGAAAGAGTATG | VNTR | SINE_R | A$_{126}$AAAGAAAGAGTATGTACCTGGATATGTATTTTT-3′
3′-CTCGGAGACAGAGTTTCTTTCTTTCTCATAC | | | T$_{126}$TTTCTTTCTCATACATGGACCTATACATAAAAA-5′

SVA_E

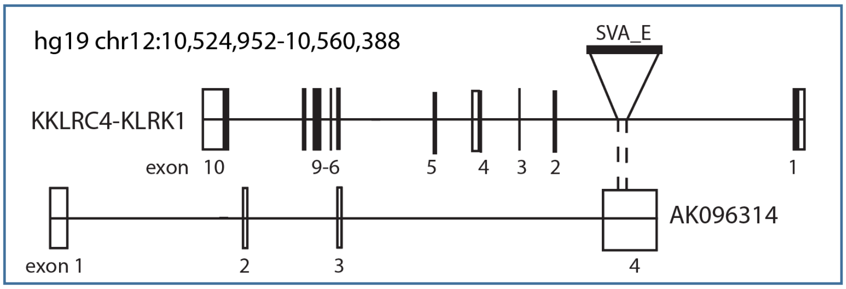


\+ strand
TSD: 14
chr12:10,549,100-10,549,113

### SVA #8
### chr1q42.2

Empty site:

5′-CATATGTTATGATTTACATGAAAAAACTAACCTGCTCAAAAGATT-3′
3′-GTATACAATACTAAATGTACTTTTTTGATTGGACGAGTTTTCTAA-5′

1st strand cleavage

Filled site:

5′-CATATGTTATGATTTACATGAAAAAACTA | MAST2 | VNTR | SINE_R | $A_{58}$GAAAAAACTAACCTGCTCAAAAGATT-3′
3′-GTATACAATACTAAATGTACTTTTTTGAT | | | | $T_{58}$CTTTTTTGATTGGACGAGTTTTCTAA-5′

SVA_F1

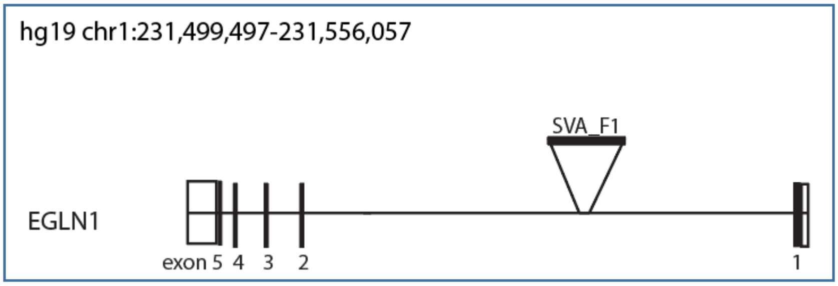


+ strand
TSD: 10
chr1:231,536,301-231,536,310

**Supplemental Figure S2.** Detailed insertion information for the 27 *de novo* MEI. The figures are listed from Alu #1 – SVA #8. Empty site refers to the non-MEI chromosome, and filled site refers to the MEI chromosome. The TSD is labeled in red. All chromosome regions are in hg19. The MEI are depicted in relation to a gene where applicable (not drawn to scale). For L1 #1, the microhomology site is labeled in green, and the deleted region is in purple. For SVA #1, there is a schematic that shows that SVA #1 likely has internal splicing from its source element on chr17.

A False negative F1 inheritance MEI locus

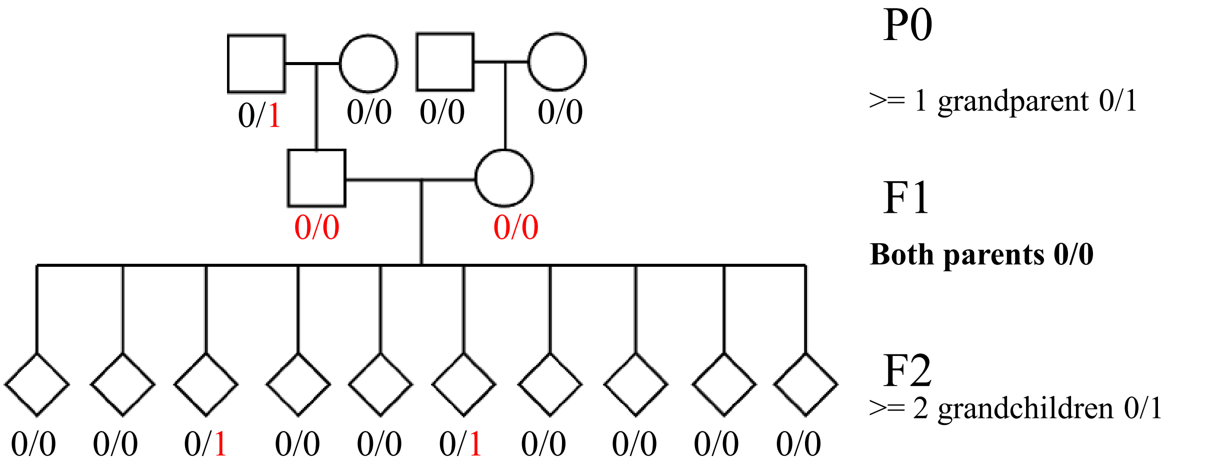


B MEI locus inherited regardless of F1 genotype

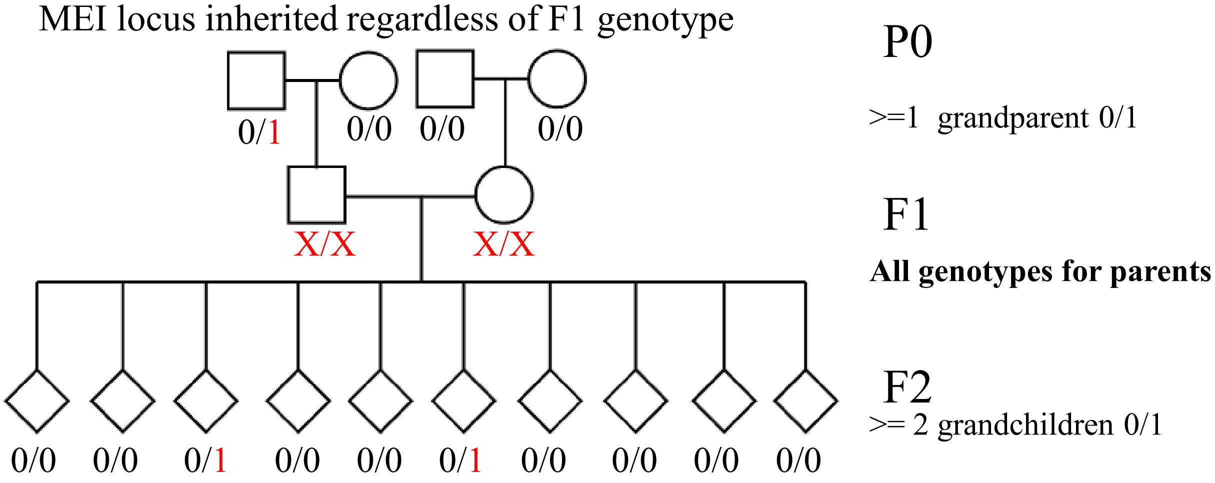


C Sensitivity rates in inherited MEI loci

| | All loci | | | Filtered loci | | |
|---|---|---|---|---|---|---|
| MEI | False Negative (#) | All inherited (#) | Sensitivity | False Negative (#) | All inherited (#) | Sensitivity |
| Alu | 2259 | 6860 | 67.1% | 221 | 3741 | 94.1% |
| L1 | 284 | 1022 | 72.2% | 26 | 506 | 94.9% |
| SVA | 128 | 480 | 73.3% | 23 | 231 | 90.0% |
| **Total** | **2,671** | **8362** | **68.1%** | **270** | **4,478** | **94.0%** |

**Supplemental Figure S3.** False-negative inheritance rate of MELT in CEPH dataset. A) Example of a false-negative inheritance MEI insertion. The pedigree has the MEI present in one P0 individuals and at least two F2 individuals, but was not called in the F1. One locus has to fail in only one pedigree to count. B) Example of inherited MEI regardless of F1 genotype. C) Sensitivity rates in inherited MEI loci. Sensitivity is calculated by (1 – False Negative Rate/Inherited loci).

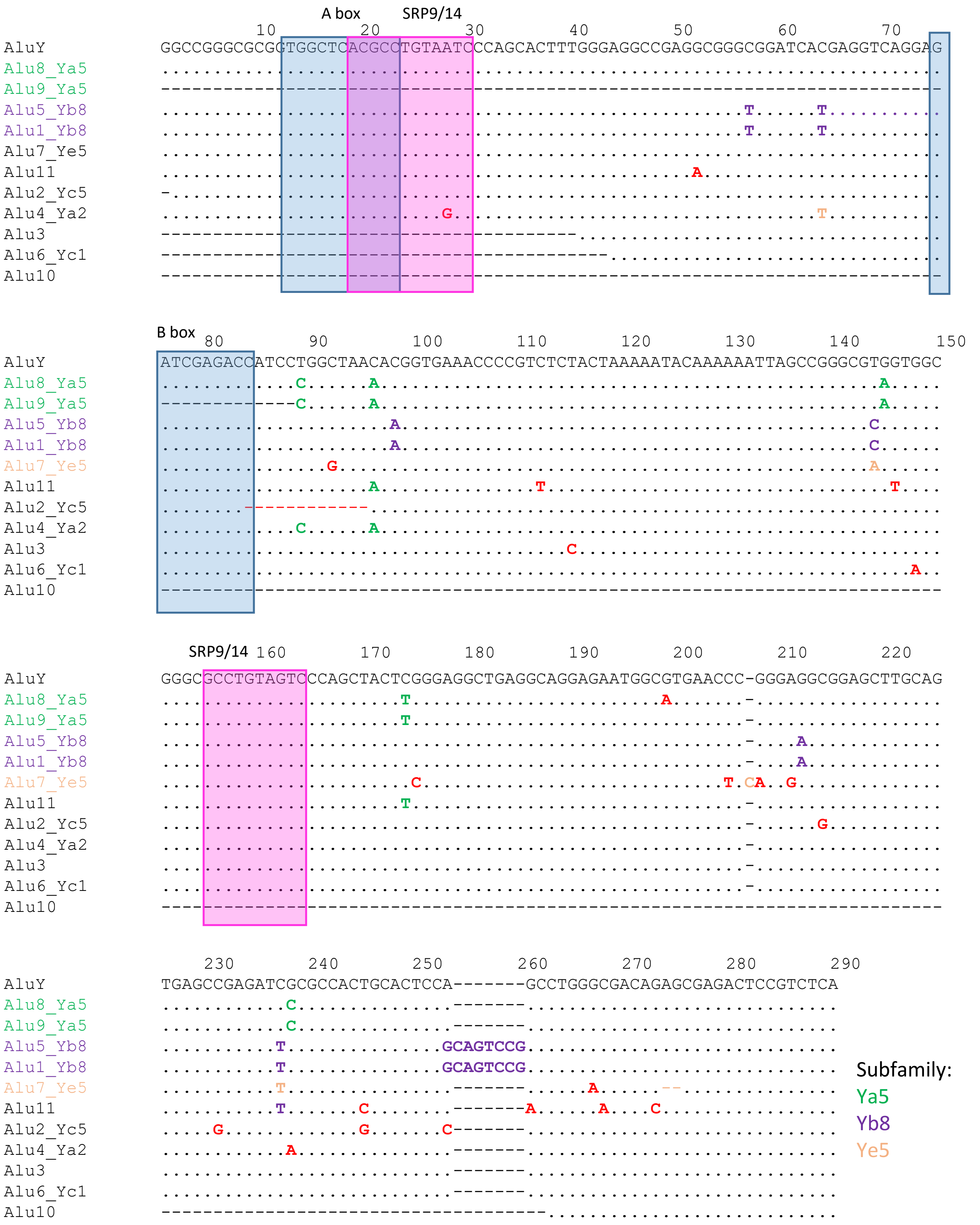

**Supplemental Figure S4.** Alignment of 11 *de novo* Alu elements to AluY consensus.

A

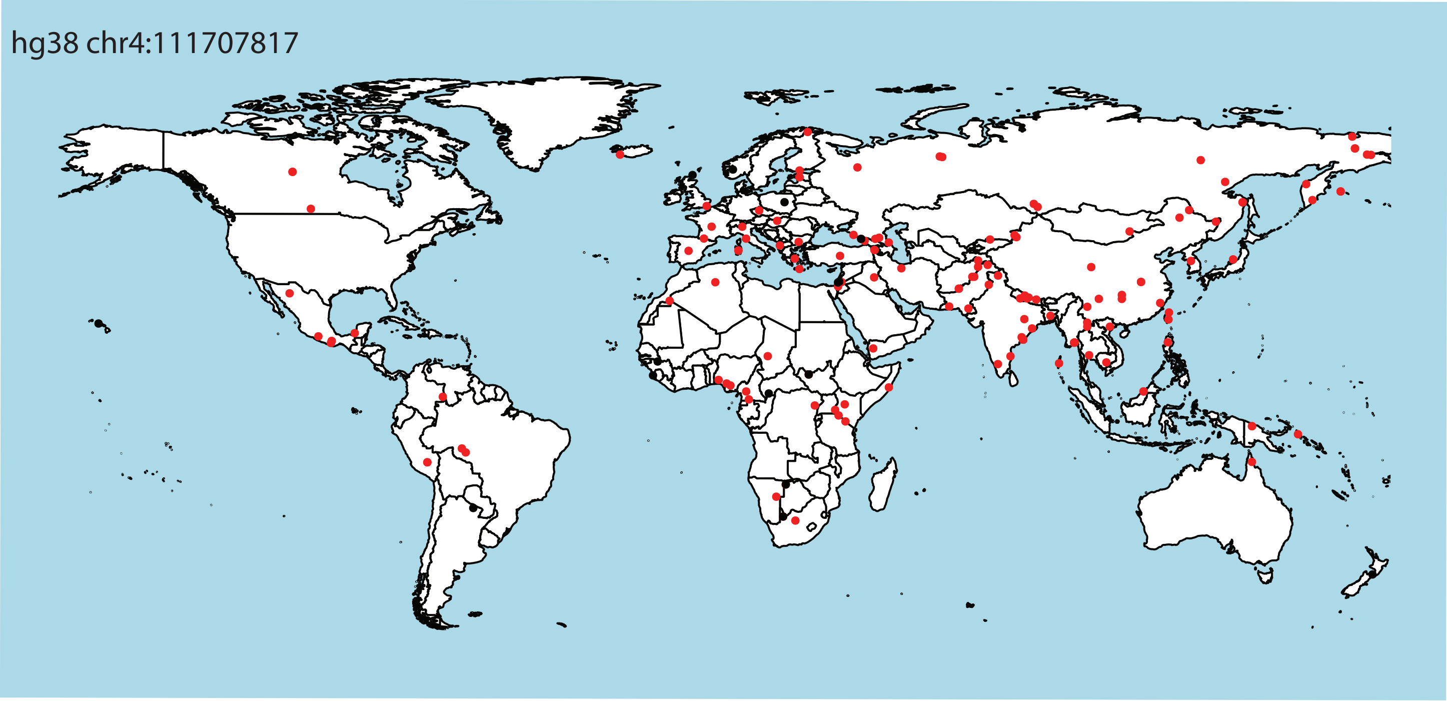


B

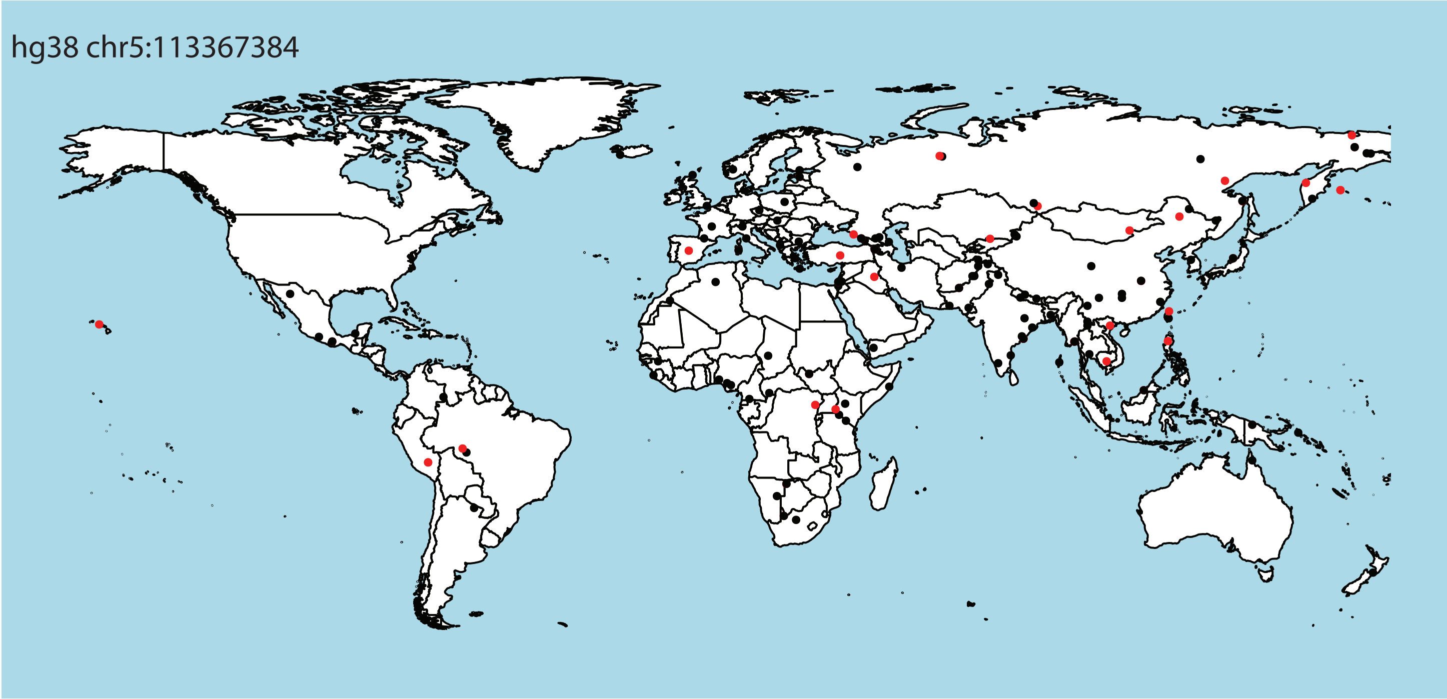

C

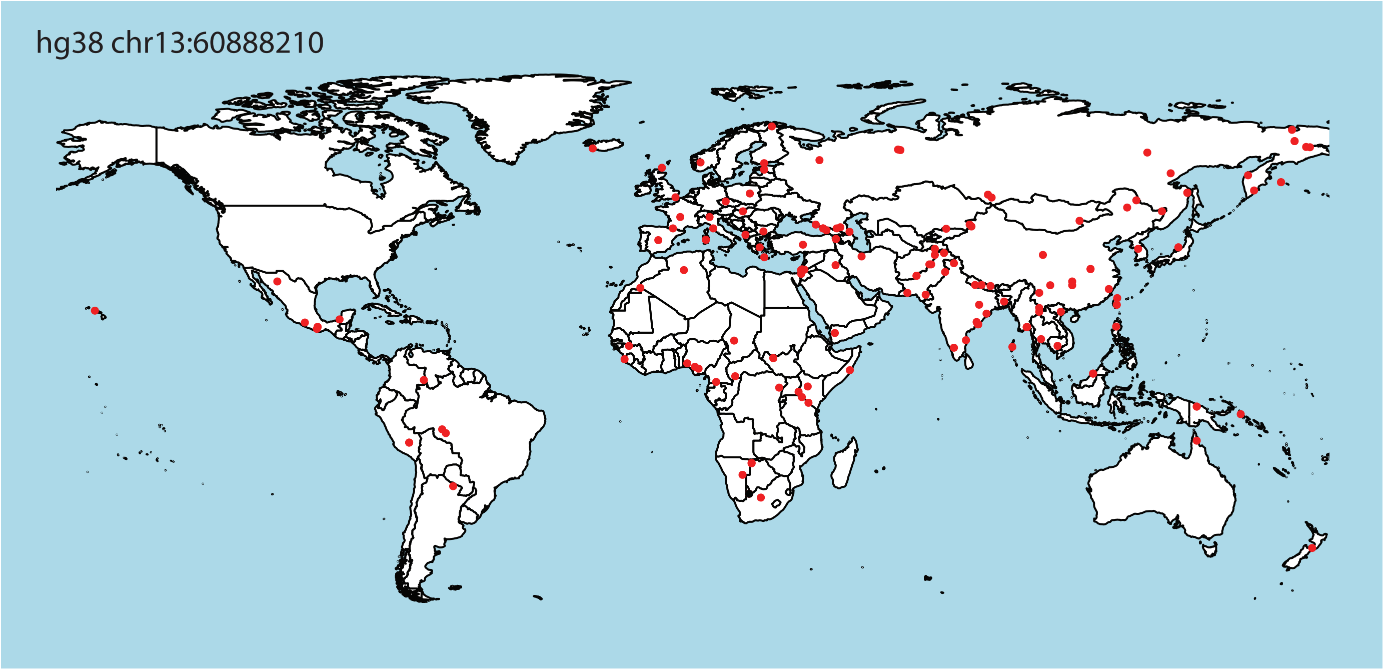


**Supplemental Figure S5.** Presence of source L1 elements within SGDP populations. A red circle indicates that the L1 element was present in that population. A black circle indicates absence of the L1 element.
