## Supplemental Methods for "Pedigree-based estimation of human mobile element retrotransposition rates"

*ME-Scan sequencing library construction for the 16 grandparent-parent trios*

Genomic DNA samples from 48 individuals were obtained from Coriell Cell Repositories (https://coriell.org/). The samples contain 16 parent-offspring trios with northern and western European ancestry from the CEPH collection. Information including population, family and individual relationships is shown in Table S1.

The ME-Scan libraries were prepared following the ME-Scan protocol described previously (Ha et al. 2017). All the adaptor and primer sequences are described previously (Feusier et al. 2017). For each sample, five µg genomic DNA was fragmented to about 1 kb in size using Covaris system (Covaris, Woburn, MA, USA) using the following protocol: duty cycle: 5%; intensity: 3; cycles/burst: 200; time: 15 seconds. The fragmented samples were concentrated using AMPure XP beads (cat. no. A63881, Beckman Coulter, Brea, CA, USA), following the manufacturer’s protocol. The concentrated DNA fragments were then used to construct the sequencing library using KAPA Library Preparation Kits with SPRI solution for Illumina (KAPA Biosystems, Wilmington, MA, USA, cat. no KK8201). For each sample, after the DNA fragments were end-repaired, A-tailed on both ends, and ligated with adaptors, the concentration of ligated DNA was quantified using Nanodrop. The 48 individual libraries were then pooled into one single library with equal concentration. All of the following steps were performed using the pooled library.

*Alu*Yb-specific first amplification was conducted for 10 cycles with 360 ng of pooled template DNA and 2.5 µl of primer, following the KAPA kit amplification protocol (initial denaturation at 98 ºC for 45 seconds, followed by the thermocycling conditions of 98 ºC for 15 seconds, 65 ºC for 30 seconds, and 72 ºC for 30 seconds, and a final extension at 72 ºC for 1 minute). The amplified PCR product was electrophoresed at 120 volts for 90 minutes on a 2 % NuSieve^R^ GTG^R^ Agarose gel (cat. no. 50080, Lonza, Rockland, Maine, USA). Fragments around 600 bp were size selected and purified using Wizard SV Gel and PCR Clean-up system (cat. no. A9281, Promega, Madison, WI, USA). After size selection, biotinylated *Alu*-enriched DNA fragments were magnetically separated from other genomic DNA fragments using 5 µl Dynabeads^R^ M-270 Streptavidin (cat. no. 65305, Invitrogen, Life Technologies, Oslo, Norway) following the manufacturer’s protocol. Second amplification was conducted for 20 cycles under the same condition as first amplification, with 3 µl of biotinylated *Alu*-enriched DNA as template using the P7 primer and a mix of six *Alu*_head primers with the Illumina P5 sequence in a 50 µl reaction. The amplified PCR product was electrophoresed at 120 volts for 90 minutes on a 2 % NuSieve^R^ GTG^R^ Agarose gel (cat. no. 50080, Lonza, Rockland, Maine, USA). Fragments around 400 bp were size selected and purified using Wizard SV Gel and PCR Clean-up system (cat. no. A9281, Promega, Madison, WI, USA). Before the library was sequenced, its fragment size and concentration was determined using Bioanalyzer and quantitative PCR by the RUCDR Infinite Biologics (Piscataway, NJ, USA). The library was sequenced using the Illumina Hiseq 2000 with 100PE format at RUCDR Infinite Biologics.
