## Supplemental Data 4 for "Pedigree-based estimation of human mobile element retrotransposition rates"

Supplemental Data S4

Key:

TSD

Transduced sequence

**TTTT signal**

MEI sequence (*Contig sequence)*

Cleavage site

>Alu1

ACTACACAGTGTAATGAATATCATAGCACAGGCTTCATTATGCTGTGGAATTATTAGTGATAAGCTACTAATAGAAAAAGATAGATATTTGTTGGCCTTATTCACTGCATTCTATTTCAACAGTGGCCACAAAATTATTTCCATACACAAATGTTTTTGTAGGATGTAATATTGTGTTAGTGTGATTTATTTACACCCTAAAGCAAATATAATTCAAAATATATGTTATAGAAGCTCTTGCTGTCTTCCCTCAATATCAGCAATGC**AAAAA**GAAAGAAACTGCTTTTTTTTTTTTTTTTTTTTTTTTTTTTTTTTTTTTTTTTTTTTTTTTTTTTTTTTTTTTTTTTTTTTGAGACGGAGTCTCGCTCTGTCGCCCAGGCCGGACTGCGGACTGCAGTGGCGCAATCTCGGCTCACTGCAAGCTCCGCTTCCCGGGTTCACGCCATTCTCCTGCCTCAGCCTCCCGAGTAGCTGGGACTACAGGCGCCCGCCACCGCGCCCGGCTAATTTTTTGTATTTTTAGTAGAGACGGGGTTTCACCTTGTTAGCCAGGATGGTCTCGATCTCCTGACCTCATGATCCACCCGCCTCGGCCTCCCAAAGTGCTGGGATTACAGGCGTGAGCCACCGCGCCCGGCCGAAACTGCTTTTTATTATAAGAAATAACTCAAAAAGGAGCAACATAAAAGTACATTTGG

>Alu2

AATCGCTTTAACCCGGGAGACAAAGGTTGTAGTGAGCCGAGATCGTGTCACCGCACTCCAGCCTGGGTGACAGAGCAAGACTCCATCTCAAAAATAGAAACAGCCGGGCGCGGTGGCTCACGCCTGTAATCCCAGCACTTTGGGAGGCCGAGGCGGGCGGATCACGAGGTCAGGAGATCGAGACCACGGTGAAACCCCGTCTCTACTAAAAATACAAAAAATTAGCCGGGCGTGGTGGCGGGCGCCTGTAGTCCCAGCTACTCGGGAGGCTGAGGCAGGAGAATGGCGTGAACCCGGGAGGGGGAGCTTGCAGTGAGCGGAGATCGCGCCACGGCACTCCCGCCTGGGCGACAGAGCGAGACTCCGTCTCAAAAAAAAAAAAAAAAAAAAAAAAAAAAAAAAAAAAAAAAAAAAAAAAAAAAAAAAAAAAAAAAATAGAAACAAAAACAAACAAAAAATTAGCCCCAAATTTGTGAGCATTTAGGTTGATTCCTATCTTCTGCTGTCTCAA

>Alu3

AATGGGGATTTGCTCTTCCTAAGGGTAGTTTGCAAAGAATATTTTTTTATGTGTAATGGTATTTTGGATTTGGTATGTATTTTGATATCCAGTAACTTGTACATAGGAGGTGTTTAGTAAATACTTGCTGAATTAATAGGCTACTCAAAATAATTTATGTTTTTTAAATTTCTACACATTTATTAAGATTTATCTGGGAGGCCGAGGCGGGCGGATCACGAGGTCAGGAGATCGAGACCATCCTGGCTAACACGGTGAAACCCCGTCTCCACTAAAAATACAAAAAATTAGCCGGGCGTGGTGGCGGGCGCCTGTAGTCCCAGCTACTCGGGAGGCTGAGGCAGGAGAATGGCGTGAACCCGGGAGGCGGAGCTTGCAGTGAGCCGAGATCGCGCCACTGCACTCCAGCCTGGGCGACAGAGCGAGACTCCGTCTCAAAAAAAAAAAAAAAAAAAAAAAAAAAAAAAAAAAAAAAAAAAAAAAAAAAAAAAAAAAAAAAAAAAAAAAAAAAAAAAAGATTTATCTATTGAACAGTCAACTCCTAAAAACACTAGGAATAATAACCTTGCAGGTATTTGAAGACCATTTGGAGGTTCATTGAGCTTTTTGGAGGATTATATCCCAGAGGGGACCTGTCACGTTCTTCTACTT

>Alu4

ATTATGATGAGACAGCCTGGGTATCTTGGTTGCGGGTGAAGGATCCACTCACTTATTGCGGTTTTTTTCCGTGGGAGCCTCAAAACACCACTGACTGAGAGGTGGGGCATTTAAGAAGCAATCAGATAATGAGGGTTCTATCCTCATGAATGAATTAACTCCCAGTTATAGAATAGTGGATTAATGGGTTGATGTATT**T**GGCCGGGCGCGGTGGCTCACGCCTGTAGTCCCAGCACTTTGGGAGGCCGAGGCGGGCGGATCATGAGGTCAGGAGATCGAGACCATCCCGGCTAAAACGGTGAAACCCCGTCTCTACTAAAAATACAAAAAATTAGCCGGGCGTGGTGGCGGGCGCCTGTAGTCCCAGCTACTCGGGAGGCTGAGGCAGGAGAATGGCGTGAACCCGGGAGGCGGAGCTTGCAGTGAGCCGAGATCACGCCACTGCACTCCAGCCTGGGCGACAGAGCGAGACTCCGTCTCAAAAAAAAAAAAAAAAAAAAAAAAAAAAAAAAAAAAAAAAAGAAACATGGAACTCAATAGAAAAGAAGCATCTAAACGGTTGAATACGATTGCCACTTAGGTGTGGGAATCAGAAGTGGGCAGGAAACAGCTA**TTTTT**AATTACAATCCTCTAGAAGAATTTCTTTTTTTAAACAATGTGCATATATGTACATAGTGTGCAAATCTATTAATGTAATTAATCTTATCAACATTTTAAAAGAGCTGCTAAAGCATCATCTAGTAAAATCAATAAACCACTTAGTGTTAAAACATT

>Alu5

CACACACACACACACACACACACACACACACACACACACAAAGAATGAAATATCTAAGAACTGTAGGACAATTATAAAAGCTGTAATATACATGTAATAAAAAAAATCACGGCCGGGCGCGGTGGCTCACGCCTGTAATCCCAGCACTTTGGGAGGCCGAGGCGGGTGGATCATGAGGTCAGGAGATCGAGACCATCCTGGCTAACAAGGTGAAACCCCGTCTCTACTAAAAATACAAAAAATTAGCCGGGCGCGGTGGCGGGCGCCTGTAGTCCCAGCTACTCGGGAGGCTGAGGCAGGAGAATGGCGTGAACCCGGGAAGCGGAGCTTGCAGTGAGCCGAGATTGCGCCACTGCAGTCCGCAGTCCGGCCTGGGCGACAGAGCGAGACTCCGTCTCAAAAAAAAAAAAAAAAAAAAAAAAAAAAAAAAAAAAAAAAAAAAAAAAAAAAAAAAAAAAAAAAAATCACAAGGTGTGATATACATGTAAAGAAACATACTGGAAGGAAGAAAGAAAGAAATAGAGAAAGAGAGAAAGAGAAAAAA

>Alu6

CATCCTTTTGCTAAGATCTTGAGTGAAATAGAGAACATGGAACTCAGAGGAACCTTAGATCAGCATTTTAAAAAAATGTTTGTGGCCTGACAAGAGCTTGTTTTAAAAAATTAATTCGTTAGGCCGAGGCGGGCGGATCACGAGGTCAGGAGATCGAGACCATCCTGGCTAACACGGTGAAACCCCGTCTCTACTAAAAATACAAAAAATTAGCCGGGCGTGGTAGCGGGCGCCTGTAGTCCCAGCTACTCGGGAGGCTGAGGCAGGAGAATGGCGTGAACCCGGGAGGCGGAGCTTGCAGTGAGCCGAGATCGCGCCACTGCACTCCAGCCTGGGCGACAGAGCGAGACTCCGTCTCAAAAAAAAAAAAAAAAAAAAAAAATTAATTCGTTAATTAAAAAAAAATTATAAACTGTTCTCTTTTTCAATTACAAAAATGAAACAAAGGCT

>Alu7

TTTCTTAACTCAAAGGAATACAGGCTCATTGCAGAAGACACAGAAAATAAAAATAAACAAAAATAAGAAATAAAATCAGGGCCGGGCGCGGTGGCTCACGCCTGTAATCCCAGCACTTTGGGAGGCCGAGGCGGGCGGATCACGAGGTCAGGAGATCGAGACCATCCTGGGTAACACGGTGAAACCCCGTCTCTACTAAAAATACAAAAAATTAGCCGGGCGAGGTGGCGGGCGCCTGTAGTCCCAGCTACTCCGGAGGCTGAGGCAGGAGAATGGCGTGAACTCCAGGGGGCGGAGCCTGCAGTGAGCCGAGATTGCGCCACTGCACTCCAGCCTGGACGACAGCGAGACTCCGTCTCAAAAAAAAAAAAAAAAAAAAAAAAAAAAAAAAAAAAAAAAAAAAAAAGAAATAAAATCAGCCATAATCCTACTACTCAGAG

>Alu8

ATGACAATTATAAATATATATGCACCCAACATTGGAACACCCAGATGTAT**AAAA**CAAATATTTTTTTTTTTTTTTTTTTTTTTTTTTTTTTTTTTTTTTTTTTTTTTTTTTTTTTTTTTTTTTTTTTTTTTTTTTTTTTTTTTTTTTTTTTTTTTTTTTTTTTTTTTTTTTTGAGACGGAGTCTCGCTCTGTCGCCCAGGCTGGAGTGCAGTGGCGGGATCTCGGCTCACTGCAAGCTCCGCCTCCCGGGTTCATGCCATTCTCCTGCCTCAGCCTCCCAAGTAGCTGGGACTACAGGCGCCCGCCACTACGCCCGGCTAATTTTTTGTATTTTTAGTAGAGACGGGGTTTCACCGTTTTAGCCGGGATGGTCTCGATCTCCTGACCTCGTGATCCGCCCGCCTCGGCCTCCCAAAGTGCTGGGATTACAGGCGTGAGCCACCGCGCCCGGCCATAAAACAAATATTATTAGATCTAAAGAAAGAGATAAACTCTAATACAATAGTAGTTGGTGACTTCAACATCCCACT

>Alu9

GTGGTCATTATTTTGCTTCGGTTTACATTTAATTTCTTTTAATCTGGAGTCCCAGCAAGCTGCCTTGTAAAATGTCATATATCTGGAATTGCTTGATTTTTTTTCCCTCATGACTAGAGTCAAGTTCCATCAAATCAGGAGGCACCGGCTAAAACGGTGAAACCCCGTCTCTACTAAAAATACAAAAAATTAGCCGGGCGTAGTGGCGGGCGCCTGTAGTCCCAGCTACTTGGGAGGCTGAGGCAGGAGAATGGCGTGAACCCGGGAGGCGGAGCTTGCAGTGAGCCGAGATCCCGCCACTGCACTCCAGCCTGGGCGACAGAGCGAGACTCCGTCTCAAAAAAAAAAAAAAAAAAAAAAAAAAAAAAAAAAAAAAAAAAAAAAAAAAAAAAAAAAAAAAAAAAAAAAAAAAAAAAAAAAAAAAAAAAAAAAAAAAAAAAAAAAATCAGGAGGCACAAAATATCGGTGCATCTCATTGTTGTGATGCCAAGTTTGATCACTTAAGGCAATATTGACTAGATTTCTCCATCATAAATAAATCTTTTTCCTTCTGTAATTAATAACTGTGAGATATTTTGTAATAATGTAAATATCCTATATGCCAGCT

>Alu10

CTGGCCTCCTCTGCTGGGAGTGGCTCTTCCTCCCAAACTCCAAGTGTGCTGCTGTGTCAGGCCCCAATGTAGCCACAGTCCTCTGTGGAAGTGTCCCTATTCTCGTTTTCCTGCTTTATTCTAAGTCATTTTTTTTTTTTTTTTTTTTTTTTTTTTTTTTTTTTTTTTTTTTTTTTTTTTTTTTTTTTTTTTTTTTTTTTTTTTTTTTTTTTTTTTTTTTTTTTTTTTTTTTTTTTTTTTTGAGACGGAGTCTCGCTCTGTCGCCCAGTAAGTCATTTTAACTATATTTATTACATACAAAATGTCCCTTCCAAACTAAAATATAACCCTTTGAGGGTATGGCTTTCG

>Alu11

TCCAGGTTCCAGGACCACAGCAAATTTAGAAGCCAACAAGAAAAAGGGAGGGAGACATGAATCAAAGCAATGTACAGAAGAAAGAAAAAGGAGCAGGGCATTACCAAGAAACAGACCTCACAAACCTGAAATGTTAATCACGGCAATGATGAGCACAGTGGTAAACAACCGTTACACCCCCAACCACCCAGGAGTGCCACTGACTCACAATACTAACGGGACGTGTGAACAGGGACAGCCGCCACTGGCAATAGTTGCACAATTCGAGCACAGAAATGAGTAAGAGGAATTATTTATACTGAGTTTT**AAAA**GAATCCGTAAGTCCAGATTGATAAACATATACACCTACTATGTACCCATCAGATTTTTTTTTTTTTTTTTTTTTTTTTTTTTTTTTTTTTTTTTTTTTTTTTTTTTTTTTTTTTTTTTTTTTTTTTTTTGAGACGGAGTCTCGCTGTGTCTCCCAGGTTGGAGTGCGGTGGCGCAATCTCGGCTCACTGCAAGCTCCGCCTCCCGGGTTCACGCCATTCTCCTGCCTCAGCCTCCCAAGTAGCTGGGACTACAGGCGCCCGCCAACACGCCCGGCTAATTTTTTGTATTTTTAGTAGAAACGGGGTTTCACCGTGTTAGCCAAGATGGTCTCGATCTCCTGACCTCGTGATCCGCCCGTCTCGGCCTCCCAAAGTGCTGGGATTACAGGCGTGAGCCACCGCGCCCGGCCCCATCAGATTTTTTAAAAAAGAAATTAATACTATACAAACACAAAGTCTTCTTCGCAGGAGAATTCCATCTAATAAATATCGTAAAAAGACAAAAATACAAGATCTTCATTTTATGCTTATCATAGTAGTAACTAATTCAGGCAAGAATCATCAGTAGATGCTAGAACCACTGGGTAAAAAAATACTGTGTCATTCACAACTGTGAAAATAGTAGAAAAGGTCCATATAGTCATCAACAGATGAGTGGATGAACAAACTGTAGCATATACACAATGGAATATTAATTACTCCACCACAGAAAGGAGTGAAGTACTGACACATGCTACA

>L1_1

CCATGCCGCACCACAATTTAAAATTAGTTTGACTTCCTCTTTTCCTAATTGAATACCCTTTATTTCCTTCTCCTGCCTGATTGCCCTGGCCAGAACTTCCAACACTATGTTGAATAGGAGCGGTGAGAGAGGGCATCCCTGTCTTGTGCCGGTTTTCAAAGGGAATGCTTCCAGTTTTTGCCCATTCAGTATGATATTGGCTGTGGGTTTGTCATAGATAGCTCTTATTATTTTGAAATACGTCCCATCAATACCTAATTTATTGAGAGTTTTTAGCATGAAGGGTTGTTGAATTTTGTCAAAGGCTTTTTCTGCATCTATTGAGATAATCATGTGGTTTTTGTCTTTGGCTCTGTTTATATGCTGGATTACATTTATTGATTTGCGTATATTGAACCAGCCTTGCATCCCAGGGATGAAGCCCACTTGATCATGGTGGATAAGCTTTTTGATGTGCTGCTGGATTCGGTTTGCCAGTATTTTATTGAGGATTTTTGCATCAATGTTCATCAAGGATATTGGTCTAAAATTCTCTTTTTTGGTTGTGTCTCTGCCCGGCTTTGGTATCAGAATAATGCTGGCCTCATAAAATGAGTTAGGGAGGATTCCCTCTTTTTCTATTGATTGGAATAGTTTCAGAAGGAATGGTACCAGTTCCTCCTTGTACCTCTGGTAGAATTCGGCTGTGAATCCATCTGGTCCTGGACTCTTTTTGGTTGGTAAACTATTGATTATTGCCACAATTTCAGAGCCTGTTATTGGTCGATTCAGAGATTCAACTTCTTCCTGGTTTAGTCTTGGGAGAGTGTATGTGTCGAGGAATGTATCCATTTCTTCTAGATTTTCTAGTTTATTTGCGTAGAGGTGTTTGTAGTATTCTCTGATGGTAGTTTGTATTTCTGTGGGATCGGTGGTGATATCCCCTTTATCATTTTTTATTGTGTCTATTTGATTCTTCTCTCTTTTTTTCTTTATTAGTCTTGCTAGCGGTCTATCAATTTTGTTGATCCTTTCAAAAAACCAGCTCCTGGATTCATTGATTTTTTGAAGGGTTTTTTGTGTCTCTATTTCCTTCAGTTCTGCTCTGATTTTAGTTATTTCTTGCCTTCTGCTAGCTTTTGAATGTGTTTGCTCTTGCTTTTCTAGTTCTTTTAAATGTGATGTTAGGGTGTCAATTTTGGATCTTTCCTGCTTTCTCTTGTAGGCATTTAGTGCTATAAATTTCCCTCTACACACTGCTTTGAATGCGTCCCAGAGATTCTGGTATGTGGTGTCTTTGTTCTCGTTGGTTTCAAAGAACATCTTTATTTCTGCCTTCATTTCGTTATGTACCCAGTAGTCATTCAGGAGCAGGTTGTTCAGTTTCCATGTAGTTGAGCGGCTTTGAGTGAGATTCTTAATCCTGAGTTCTAGTTTGATTGCACTGTGGTCTGAGAGATAGTTTGTTATAATTTCTGTTCTTTTACATTTGCTGAGGAGAGCTTTACTTCCAACTATGTGGTCAATTTTGGAATAGGTGTGGTGTGGTGCTGAAAAAAATGTATATTCTGTTGATTTGGGGTGGAGAGTTCTGTAGATGTCTATTAGGTCTGCTTGGTGCAGAGCTGAGTTCAATTCCTGGGTATCCTTGTTGACTTTCTGTCTCGTTGATCTGTCTAATGTTGACAGTGGGGTGTTAAAGTCTCCCATTATTAATGTGTGGGAGTCTAAGTCCTCTGCTTGTTCCTGATTAAATGGTTAATGTAAATTAATTTAATTTAGAATCGGTTCTACTGCTATAACCAATACAAGCCAGACTTCTACTTGGAATCTTAGATCATATAGCTGACTGATATTGAAGAATATAGGAATGGTTCAAAAGAGAACCACAAAGTAACTATTAAGAAACAAAAGCCTCTCAAGAAGGTTTTAAGAAGTAAGATTTTAGAATCCAGATAAAGGGTAGGTAGCTAAGTGGCAACACAGTTACAGCATAATATGGTGTAGAAAGACTAGACATTTTTTCCTTTCTGGTGAGTGAACAAAAACAAATGTCTTCAGTGACATAGAAGAGATACAGGGGCACTTTCTAGGAACATGTGTAGGTAAGTTGCTTACAACCACGCACAGTGGAAAAACCACCCTGCTA

>L1_2

TTATACAAGCTGTTGGACCTCTTAAGGACGTTGACATAAGGACCACTTTCTGTGAAATAAGAATTAGTGGTTTAAAAAGAGTGCACTTAACAGTATCATGTAATAGAGCCAAGATGGCCGAATAGGAACAGCTCCGGTCTACAGCTCCCAGCGTGAGCGACGCAGAAGACGGTGATTTCTGCATTTCCATCTGAGGTACCGGGTTCATCTCACTAGGGAGTGCCAGACAGTGGGCGCAGGCCAGTGTGTGTGCGCACCGTGCGCGAGCCGAAGCAGGGCGAGGCATTGCCTCACCTGGGAAGCGCAAGGGGTCAGGGAGTTCCCTTTCCGAGTCAAAGAAAGGGGTGACGGACGCACCTGGAAAATCGGGTCACTCCCACCCGAATATTGCGCTTTTCAGACCGGCTTAAGAAACGGCGCACCATGAGACTATATCCCACACCTGGCTCGGAGGGTCCTACGCCCACGGAATCTCGCTGATTGCTAGCACAGCAGTCTGAGATCAAACTGCAAGGCGGCAACGAGGCTGGGGGAGGGGCGCCCGCCATTGCCCAGGCTTGCTTAGGTAAACAAAGCAGCCGGGAAGCTCGAACTGGGTGGAGCCCACCACAGCTCAAGGAGGCCTGCCTGCCTCTGTAGGCTCCACCTCTGGGGGCAGGGCACAGACAAACAAAAAGACAGCAGTAACCTCTGCAGACTTAAGTGTCCCTGTCTGACAGCTTTGAAGAGAGCAGTGGTTCTCCCAGGAGGCAGCTGGAGATCTGAGAACGGGCAGACTGCCTCCTCAAGTGGGTCCCTGACTCCTGACCCCCGAGCAGCCTAACTGGGAGGCACCCCCCAGCAGGGGCACACTGACACCTCACACGGCAGGGTATTCCAACAGACCTGCAGCTGAGGGTCCTGTCTGTTAGAAGGAAAACTAACAACCAGAAAGGACATCTACACCGAAAACCCATCTGTACATCACCATCATCAAAGACCAAAAGTAGATAAAACCACAAAGATGGGGAAAAAACAGAACAGAAAAACTGGAAACTCTAAAACGCAGAGCGCCTCTCCTCCTCCAAAGGAACGCAGTTCCTCACCAGCAACAGAACAAAGCTGGATGGAGAATGATTTTGACGAGCTGAGAGAAGAAGGCTTCAGACGATCAAATTACTCTGAGCTACGGGAGGACATTCAAACCAAAGGCAAAGAAGTTGAAAACTTTGAAAAAAATTTAGAAGAATGTATAACTAGAATAACCAATACAGAGAAGTGCTTAAAGGAGCTGATGGAGCTGAAAACCAAGGCTCGAGAACTACGTGAAGAATGCAGAAGCCTCAGGAGCCGATGCGATCAACTGGAAGAAAGGGTATCAGCAATGGAAGATGAAATGAATGAAATGAAGCGAGAAGGGAAGTTTAGAGAAAAAAGAATAAAAAGAAATGAGCAAAGCCTCCAAGAAATATGGGACTATGTGAAAAGACCAAATCTACGTCTGATTGGTGTACCTGAAAGTGATGTGGAGAATGGAACCAAGTTGGAAAACACTCTGCAGGATATTATCCAGGAGAACTTCCCCAATCTAGCAAGGCAGGCCAACGTTCAGATTCAGGAAATACAGAGAACGCCACAAAGATACTCCTCGAGAAGAGCAACTCCAAGACACATAATTGTCAGATTCACCAAAGTTGAAATGAAGGAAAAAATGTTAAGGGCAGCCAGAGAGAAAGGTCGGGTTACCCTCAAAGGAAAGCCCATCAGACTAACAGCGGATCTCTCGGCAGAAACCCTACAAGCCAGAAGAGAGTGGGGGCCAATATTCAACATTCTTAAAGAAAAGAATTTTCAACCCAGAATTTCATATCCAGCCAAACTAAGCTTCATAAGTGAAGGAGAAATAAAATACTTTATAGACAAGCAAATGTTGAGAGATTTTGTCACCACCAGGCCTGCCCTAAAAGAGCTCCTGAAGGAAGCGCTAAACATGGAAAGGAACAACCGGTACCAGCCGCTGCAAAATCATGCCAAAATGTAAAGACCATCGAGACTAGGAAGAAACTGCATCAACTAATGAGCAAAATCACCAGCTAACATCATAATGACAGGATCAAATTCACACATAACAATATTAACTTTAAATATAAATGGACTAAATTCTGCAATTAAAAGACACAGACTGGCAAGTTGGATAAAGAGTCAAGACCCATCAGTGTGCTGTATTCAGGAAACCCATCTCACGTGCAGAGACACACATAGGCTCAAAATAAAAGGATGGAGGAAGATCTACCAAGCCAATGGAAAACAAAAAAAGGCAGGGGTTGCAATCCTAGTCTCTGATAAAACAGACTTTAAACCAACAAAGATCAAAAGAGACAAAGAAGGCCATTACATAATGGTAAAGGGATCAATTCAACAAGAGGAGCTAACTATCCTAAATATTTATGCACCCAATACAGGAGCACCCAGATTCATAAAGCAAGTCCTCAGTGACCTACAAAGAGACTTAGACTCCCACACATTAATAATGGGAGACTTTAACACCCCACTGTCAACATTAGACAGATCAACGAGACAGAAAGTCAACAAGGATACCCAGGAATTGAACTCAGCTCTGCACCAAGCAGACCTAATAGACATCTACAGAACTCTCCACCCCAAATCAACAGAATATACATTTTTTTCAGCACCACACCACACCTATTCCAAAATTGACCACATAGTTGGAAGTAAAGCTCTCCTCAGCAAATGTAAAAGAACAGAAATTATAACAAACTATCTCTCAGACCACAGTGCAATCAAACTAGAACTCAGGATTAAGAATCTCACTCAAAGCCGCTCAACTACATGGAAACTGAACAACCTGCTCCTGAATGACTACTGGGTACATAACGAAATGAAGGCAGAAATAAAGATGTTCTTTGAAACCAACGAGAACAAAGACACCACATACCAGAATCTCTGGGACGCATTCAAAGCAGTGTGTAGAGGGAAATTTATAGCACTAAATGCCTACAAGAGAAAGCAGGAAAGATCCAAAATTGACACCCTAACATCACAATTAAAAGAACTAGAAAAGCAAGAGCAAACACATTCAAAAGCTAGCAGAAGGCAAGAAATAACTAAAATCAGAGCAGAACTGAAGGAAATAGAGACACAAAAAACCCTTCAAAAAATCAATGAATCCAGGAGCTGGTTTTTTGAAAGGATCAACAAAATTGATAGACCGCTAGCAAGACTAATAAAGAAAAAGAGAGAGAAGAATCAAATAGACACAATAAAAAATGATAAAGGGGATATCACCACCGATCCCACAGAAATACAAACTACCATCAGAGAATACTACAAACACCTCTACGCAAATAAACTAGAAAATCTAGAAGAAATGGATACATTCCTCGACACATACACTCTCCCAAGACTAAACCAGGAAGAAGTTGAATCTCTGAATAGACCAATAACAGGCTCTGAAATTGTGGCAATAATCAATAGTTTACCAACCAAAAAGAGTCCAGGACCAGATGGATTCACAGCCGAATTCTACCAGAGGTACATGGAGGAACTGGTACCATTCCTTCTGAAACTATTCCAATCAATAGAAAAAGAGGGAATCCTCCCTAACTCATTTTATGAGGCCAGCATCATTCTGATACCAAAGCCGGGCAGAGACACAACCAAAAAAGAGAATTTTAGACCAATATCCTTGATGAACATTGATGCAAAAATCCTCAATAAAATACTGGCAAACCGAATCCAGCAGCACATCAAAAAGCTTATCCACCATGATCAAGTGGGCTTCATCCCTGGGATGCAAGGCTGGTTCAATATACGCAAATCAATAAATGTAATCCAGCATATAAACAGAGCCAAAGACAAAAACCACATGATTATCTCAATAGATGCAGAAAAAGCCTTTGACAAAATTCAACAACCCTTCATGCTAAAAACTCTCAATAAATTAGGTATTGATGGGACGTATTTCAAAATAATAAGAGCTATCTATGACAAACCCACAGCCAATATCATACTGAATGGGCAAAAACTGGAAGCATTCCCTTTGAAAACCGGCACAAGACAGGGATGCCCTCTCTCACCGCTCCTATTCAACATAGTGTTGGAAGTTCTGGCCAGGGCAATCAGGCAGGAGAAGGAAATAAAGGGTATTCAATTAGGAAAAGAGGAAGTCAAATTGTCCCTGTTTGCAGATGACATGATTGTTTATCTAGAAAACCCCATCGTCTCAGCCCAAAATCTCCTTAAGCTGATAAGCAACTTCAGCAAAGTCTCAGGATACAAAATCAATGTACAAAAATCACAAGCATTCTTATACACCAACAACAGACAAACAGAGAGCCAGATCATGGGTGAACTCCCATTCACAATTGCTTCAAAGAGAATAAAATACCTAGGAATCCAACTTACAAGGGATGTGAAGGACCTCTTCAAGGAGAACTACAAACCACTGCTCAAGGAAATAAAAGAGGAGACAAACAAATGGAAGAACATTCCATGCTCATGGGTAGGAAGAATCAATATCGTGAAAATGGCCATACTGCCCAAGGTAATTTACAGATTCAATGCCATCCCCATCAAGCTACCAATGACTTTCTTCACAGAATTGGAAAAAACTACTTTAAAGTTCATATGGAACCAAAAAAGAGCCCGCATTGCCAAGTCAATCCTAAGCCAAAAGAACAAAGCTGGAGGCATCACACTACCTGACTTCAAACTATACTACAAGGCTACAGTAACCAAAACAGCATGGTACTGGTACCAAAACAGAGATATAGATCAATGGAACAGAACAGAGCCCTCAGAAATAATGCCGCATATCTACAACTATCTGATCTTTGACAAACCTGAGAAAAACAAGCAATGGGGAAAGGATTCCCTATTTAATAAATGGTGCTGGGAAAACTGGCTAGCCATATGTAGAAAGCTGAAACTGGATCCCTTCCTTACACCTTATACAAAAATCAATTCAAGATGGATTAAAGATTTAAACGTTAAACCTAAAACCATAAAAACCCTAGAAGAAAACCTAGGCATTACCATTCAGGACATAGGCGTGGGCAAGGACTTCATGTCCAAAACACCAAAAGCAATGGCAACAAAAGACAAAATTGACAAATGGGATCTAATTAAACTAAAGAGCTTCTGCACAGCAAAAGAAACTACCATCAGAGTGAACAGGCAACCTACAACATGGGAGAAAATTTTTGCAACCTACTCATCTGACAAAGGGCTAATATCCAGAATCTACAATGAACTCAAACAAATTTACAAGAAAAAAACAAACAACCCCATCAAAAAGTGGGCGAAGGACATGAACAGACACTTCTCAAAAGAAGACATTTATGCAGCCAAAAAACACATGAAGAAATGCTCATCATCACTGGCCATCAGAGAAATGCAAATCAAAACCACTATGAGATATCATCTCACACCAGTTAGAATGGCAATCATTAAAAAGTCAGGAAACAACAGGTGCTGGAGAGGATGCGGAGAAATAGGAACACTTTTACACTGTTGGTGGGACTGTAAACTAGTTCAACCATTGTGGAAGTCAGTGTGGCGATTCCTCAGGGATCTAGAACTAGAAATACCATTTGACCCAGCCATCCCATTACTGGGTATATACCCAAATGAGTATAAATCATGCTGCTATAAAGACACATGCACACGTATGTTTATTGCGGCACTATTCACAATAGCAAAGACTTGGAACCAACCCAAATGTCCAACAATGATAGACTGGATTAAGAAAATGTGGCACATATACACCATGGAATACTATGCAGCCATAAAAAATGATGAGTTCATATCCTTTGTAGGGACATGGATGAAATTGGAAACCATCATTCTCAGTAAACTATCGCAAGAACAAAAAACCAAACACCGCATATTCTCACTCATAGGTGGGAATTGAACAATGAGATCACATGGACACAGGAAGGGGAATATCACACTCTGGGGACTGTGGTGGGGTCGGGGGAGGGGGGAGGGATAGCATTGGGAGATATACCTAATGCTAGATGACACATTAGTGGGTGCAGCGCACCAGCATGGCACATGTATACATATGTAACTAACCTGCACAATGTGCACATGTACCCTAAAACTTAGAGTATAATAAAAAAAAAAAAAAAAAAAAAAAAAAAAATAAAGTATCTCATAAACTTAAATGATGTTAAGGGCAAATGTTTATATTAAAGTATCCATAAAAAGAACTAAATTTGTAAAAAAAAAAAAAAAAAAAAAAAAAAAAAAAAAAAAAAAAAAAAACCCAAAAAAAAAAAAAAAAAAAAAAAAAAAAAAAAAAAAAAAAAAAAAAAAAAACAGTATCATGTAAGTATCAGTTTCTGTATGTCATTACTAGCCAGGCTTTTTTTAAAAAACTGGACTGTGCTGCCTTGTGTGTACATTTAAATGAGGCCCTTAAAGGGAAATCTGAGCAATGTCTTCAAACAAAGACTTATAATCAGAAGGGTCTTTTTAAAAAAAAAAAAAAAAAAGAATAACTATCTTCAGCAGACCTAAAACCAGGATGGAAAGGTGGATACTTCCCCACCTGGGCCCCAAAGCAGGGCCCTTCAAAATACTGAAGGTCCAGCATTCCCTGGGCAATTCTTCCTCCTTCTCTGTCCTCCCCACCCCCATCAATACCCAGCTTGCAGTGATATTCTTTTACCCTTTACTAATCTAATGCCAAACACATGTAGGTTCTATCCTGTTTCCAGGAAAGAGGTTTCCATGAATTGCCTCATCAGCCTTACTAATAGCTACAATATATGGAGCACTTTGTGCCCAGCACTCGATTCCTTTTATACTTGATCTCTTTTCATCCTTTCTTCACAATAGCCCCATGAGGTGGGTGCTATAATTATTTCTAATTTTACAGATAAGGAAACCAAAGCTCAGAGAGTCTGAGCAATTTGCTTAAAGTTGCTTAACAGCTGTATTAGCCTACTACTTATAATTCGACTACCTGTGTCCAAGGCTACATAAATTCAGGTCGCCTGACTCAAAGCTCTTATTCAACGTGCTAGCCTCCTT

>L1_3

*ACCGTGCCCAGCCTATTTAAAAAAACTTGTCATATTTTTGGGTTTTTTGTATGTGTGTTTTTTTTTTTTTTTTTTTTTTTTTTTTTTTTTTTTTTTTTTTTTTTTTTTTTTTTTTTTTTTTTTTTTTTTTTTTTTTTTTTTTTTTTTTTTTTTTTTTTTTTTTTTTTTTTTTTTTTTTTTTTTTTTTTTTTTTTTTTTTTTTTTTTTTTTTTTTTTTTTTTTTATTATACTCTAAGTTTTAGGGTACATGTGCA*NNNNNNNNNNNNNNNNNNNNNNNNNNNNNNNNNNNNNNNNNNNNNNNNNNNNNNNNNNNNNNNNNNNNNNNNNNNNNNNNNNNNNNNNNNNNNNNNNNNNNNNNNNNNNNNNNNNNNNNNNNNNNNNNNNNNNNNNNNNNNNNNNNNNNNNNNNNNNNNNNNNNNNNNNNNNNNNNNNNNNNNNNNNNNNNNNNNNNNNNNNNNNNNNNNNNNNNNNNNNNNNNNNNNNNNNNNNNNNNNNNNNNNNNNNNNNNNNNNNNNNNNNNNNNNNNNNNNNNNNNNNNNNNNNNNNNNNNNNNNNNNNNNNNNNNNNNNNNNNNNNNNNNNNNNNNNNNNNNNNNNNNNNNNNNNNNNNNNNNNNNNNNNNNNNNNNNNNNNNNNNNNNNNNNNNNNNNNNNNNNNNNNNNNNNNNNNNNNNNNNNNNNNNNNNNNNNNNNNNNNNNNNNNNNNNNNNNNNNNNNNNNNNNNNNNNNNNNNNNNNNNNNNNNNNNNNNNNNNNNNNNNNNNNNNNNNNNNNNNNNNNNNNNNNNNNNNNNNNNNNNNNNNNNNNNNNNNNNNNNNNNNNNNNNNNNNNNNNNNNNNNNNNNNNNNNNNNNNNNNNNNNNNNNNNNNNNNNNNNNNNNNNNNNNNNNNNNNNNNNNNNNNNNNNNNNNNNNNNNNNNNNNNNNNNNNNNNNNNNNNNNNNNNNNNNNNNNNNNNNNNNNNNNNNNNNNNNNNNNNNNNNNNNNNNNNNNNNNNNNNNNNNNNNNNNNNNNNNNNNNNNNNNNNNNNNNNNNNNNNNNNNNNNNNNNNNNNNNNNNNNNNNNNNNNNNNNNNNNNNNNNNGCCATTGCTTTTGGTGTTTTGGACATGAAGTCCTTGCCCACGCCTATGTCCTGAATGGTAATGCCTAGGTTTTCTTCTAGGGTTTTTATGGTTTTAGGTTTAACGTTTAAATCTTTAATCCATCTTGAATTGATTTTTGTATAAGGTGTAAGGAAGGGATCCAGTTTCAGCTTTCTACATATGGCTAGCCAGTTTTCCCAGCACCATTTATTAAATAGGGAATCCTTTCCCCATTGCTTGTTTTTCTCAGGTTTGTCAAAGATCAGATAGTTGTAGATATGCGGCATTATTTCTGAGGGCTCTGTTCTGTTCCATTGATCTATATCTCTGTTTTGGTACCAGTACCATGCTGTTTTGGTTACTGTAGCCTTGTAGTATAGTTTGAAGTCAGGTAGTGTGATGCCTCCAGCTTTGTTCTTTTGGCTTAGGATTGACTTGGCAATGCGGGCTCTTTTTTGGTTCCATATGAACTTTAAAGTAGTTTTTTCCAATTCTGTGAAGAAAGTCATTGGTAGCTTGATGGGGATGGCATTGAATCTGTAAATTACCTTGGGCAGTATGGCCATTTTCACGATATTGATTCTTCCTACCCATGAGCATGGAATGTTCTTCCATTTGTTTGTCTCCTCTTTTATTTCCTTGAGCAGTGGTTTGTAGTTCTCCTTGAAGAGGTCCTTCACATCCCTTGTAAGTTGGATTCCTAGGTATTTTATTCTCTTTGAAGCAATTGTGAATGGGAGTTCACCCATGATCTGGCTCTCTGTTTGTCTGTTGTTGGTGTATAAGAATGCTTGTGATTTTTGTACATTGATTTTGTATCCTGAGACTTTGCTGAAGTTGCTTATCAGCTTAAGGAGATTTTGGGCTGAGACGATGGGGTTTTCTAGATAAACAATCATGTCATCTGCAAACAGGGACAATTTGACTTCCTCTTTTCCTAATTGAATACCCTTTATTTCCTTCTCCTGCCTGATTGCCCTGGCCAGAACTTCCAACACTATGTTGAATAGGAGCGGTGAGAGAGGGCATCCCTGTCTTGTGCCGGTTTTCAAAGGGAATGCTTCCAGTTTTTGCCCATTCAGTATGATATTGGCTGTGGGTTTGTCATAGATAGCTCTTATTATTTTGAAATACGTCCCATCAATACCTAATTTATTGAGAGTTTTTAGCATGAAGGGTTGTTGAATTTTGTCAAAGGCTTTTTCTGCATCTATTGAGATAATCATGTGGTTTTTGTCTTTGGCTCTGTTTATATGCTGGATTACATTTATTGATTTGCGTATATTGAACCAGCCTTGCATCCCAGGGATGAAGCCCACTTGATCATGGTGGATAAGCTTTTTGATGTGCTGCTGGATTCGGTTTGCCAGTATTTTATTGAGGATTTTTGCATCAATGTTCATCAAGGATATTGGTCTAAAATTCTCTTTTTTGGTTGTGTCTCTGCCCGGCTTTGGTATCAGAATGATGCTGGCCTCATAAAATGAGTTAGGGAGGATTCCCTCTTTTTCTATTGATTGGAATAGTTTCAGAAGGAATGGTACCAGTTCCTCCATGTACCTCTGGTAGAATTCGGCTGTGAATCCATCTGGTCCTGGACTCTTTTTGGTTGGTAAACTATTGATTATTGCCACAATTTCAGAGCCTGTTATTGGTCTATTCAGAGATTCAACTTCTTCCTGGTTTAGTCTTGGGAGAGTGTATGTGTCGAGGAATGTATCCATTTCTTCTAGATTTTCTAGTTTATTTGCGTAGAGGTGTTTGTAGTATTCTCTGATGGTAGTTTGTATTTCTGTGGGATCGGTGGTGATATCCCCTTTATCATTTTTTATTGTGTCTATTTAATTCTTCTCTCTCTTTTTCTTTATTAGTCTTGCTAGCGGTCTATCAATTTTGTTGATCCTTTCAAAAAACCAGCTCCTGGATTCATTGATTTTTTGAAGGGTTTTTTGTGTCTCTATTTCCTTCAGTTCTGCTCTGATTTTAGTTATTTCTTGCCTTCTGCTAGCTTTTGAATGTGTTTGCTCTTGCTTTTCTAGTTCTTTTAATTGTGATGTTAGGGTGTCAATTTTGGATCTTTCCTGCTTTCTCTTGTAGGCATTTAGTGCTATAAATTTCCCTCTACACACTGCTTTGAATGCGTCCCAGAGATTCTGGTATGTGGTGTCTTTGTTCTCGTTGGTTTCAAAGAACATCTTTATTTCTGCCTTCATTTCGTTATGTACCCAGTAGTCATTCAGGAGCAGGTTGTTCAGTTTCCATGTAGTTGAGCGGCTTTGAGTGAGATTCTTAATCCTGAGTTCTAGTTTGATTGCACTGTGGTCTGAGAGATAGTTTGTTATAATTTCTGTTCTTTTACATTTGCTGAGGAGAGCTTTACTTCCAACTATGTGGTCAATTTTGGAATAGGTGTGGTGTGGTGCTGAAAAAAATGTATATTCTGTTGATTTGGGGTGGAGAGTTCTGTAGATGTCTATTAGGTCTGCTTGGTGCAGAGCTGAGTTCAATTCCTGGGTATCCTTGTTGACTTTCTGTCTCGTTGATCTGTCTAATGTTGACAGTGGGGTGTTAAAGTCTCCCATTATTAATGTGTGGGAGTCTAAGTCTCTTTGTAGGTCACTGAGGACTTGCTTTATGAATCTGGGTGCTCCTGTATTGGGTGCATAAATATTTAGGATAGTTAGCTCCTCTTGTTGAATTGATCCCTTTACCATTATGTAATGGCCTTCTTTGTCTCTTTTGATCTTTGTTGGTTTAAAGTCTGTTTTATCAGAGACTAGGATTGCAACCCCTGCCTTTTTTTGTTTTCCATTGGCTTGGTAGATCTTCCTCCATCCTTTTATTTTGAGCCTATGTGTGTCTCTGCACGTGAGATGGGTTTCCTGAATACAGCACACTGATGGGTCTTGACTCTTTATCCAACTTGCCAGTCTGTGTCTTTTAATTGCAGAATTTAGTCCATTTATATTTAAAGTTAATATTGTTATGTGTGAATTTGATCCTGTCATTATGATGTTAGCTGGTGATTTTGCTCATTAGTTGATGCAGTTTCTTCCTAGTCTCGATACACACTCGTATGTGTGTTTTTTTTAAAAAAAGAAACAGGGTCTTGCTCTGTTGCTCAGGCAGGTATGCAGTGGTGAAAACATAATTCCCTGCAGCCTTGAACTTCTGGACTGAAGGGATCCTCCTGCCTCTGCCTCCCAAATGGCTGGTACTACAGGTGTACTCCACCATGCCCAGCTAATTAAAAAAAATTTTTTTTAGAGACAGGGTCTCACTGTGTTGCTCAGGCTGGTGTCAAACTTCTGACCTCAAGCGA

>L1_4

GCAAAACTTTGTAGCCCAATTTGTTCAACTATTGAAGTGGTGGTTGTGTGATGTGCAGTTAGGCGTTGTCGTGAAGAATTGGACCCTTTCTGTTGACCAATGCTGGCTGCAGGTGTTGCAGTTTTTGGTGCATCTCATTGATTTGCTGAGCATACTTCTCAGATGTAACGGTTTCGCTGGGATTCAGAAACCTGTAGTGGATCAGAGCGACAGCAGACCACCAAACAGTGACCATGACCTTTTTTGGTGCAAGTTTGGCTTCGGGAAATGCTTTGGAGCTTCTTCTCAGTCCAATAACTGAGCTATTCATTGTCGGTTGTTGTAGAAAATCCACTTTTCATTGCATGTCACATCCAATCGAGAAATGGGTCCTTGTTGTTGTGTAGAATAAGAGAAGACAACACTTTAAAACAACAATTTTTTTTTTTTTTTTACTCAACTCATGAGGCACCCATTTACTGAGATTTCTCACCTTTCCAATTTGCTTCAAATGCTGAATGACCACAGAATGGTCGACGTTGAGTTCTTCACCAACTTCTCGTGTAGTTGTAAGAGGAGCAGCTATTTAATAAATGGTGCTGGGAAAACTGGCTAGCCATATGTAGAAAGCTGAAACTGGATCCCTTCCTTACACCTTATACAAAAATCAATTCAAGATGGATTAAAGATTTAAACGTTAAACCTAAAACCATAAAAACCCTAGAAGAAAACCTAGGCATTACCATTCAGGACATAGGCGTGGGCAAGGACTTCATGTCCAAAACACCAAAAGCAATGGCAACAAAAGACAAAATTGACAAATGGGATCTAATTAAACTAAAGAGCTTCTGCACAGCAAAAGAAACTACCATCAGAGTGAACAGGCAACCTACAACATGGGAGAAAATTTTTGCAACCTACTCATCTGACAAAGGGCTAATATCCAGAATCTACAATGAACTCAAACAAATTTACAAGAAAAAAACAAACAACCCCATCAAAAAGTGGGCGAAGGACATGAACAGACACTTCTCAAAAGAAGACATTTATGCAGCCAAAAAACACATGAAGAAATGCTCATCATCACTGGCCATCAGAGAAATGCAAATCAAAACCACTATGAGATATCATCTCACACCAGTTAGAATGGCAATCATTAAAAAGTCAGGAAACAACAGGTGCTGGAGAGGATGCGGAGAAATAGGAACACTTTTACACTGTTGGTGGGACTGTAAACTAGTTCAACCATTGTGGAAGTCAGTGTGGCGATTCCTCAGGGATCTAGAACTAGAAATACCATTTGACCCAGCCATCCCATTACTGGGTATATACCCAAATGAGTATAAATCATGCTGCTATAAAGACACATGCACACGTATGTTTATTGCGGCACTATTCACAATAGCAAAGACTTGGAACCAACCCAAATGTCCAACAATGATAGACTGGATTAAGAAAATGTGGCACATATACACCATGGAATACTATGCAGCCATAAAAAATGATGAGTTCATATCCTTTGTAGGGACATGGATGAAATTGGAAACCATCATTCTCAGTAAACTATCGCAAGAACAAAAAACCAAACACCGCATATTCTCACTCATAGGTGGGAATTGAACAATGAGATCACATGGACACAGGAAGGGGAATATCACACTCTGGGGACTGTGGTGGGGTCGGGGGAGGGGGGAGGGATAGCATTGGGAGATATACCTAATGCTAGATGACACATTAGTGGGTGCAGCGCACCAGCATGGCACATGTATACATATGTAACTAACCTGCACAATGTGCACATGTACCCTAAAACTTAGAGTATAATAAAAAAAAAATAATAATAATAATAATAATAATAATAATTTAAGATGTACATATATTCCATCAAATTTAGTGTGAAGGCTTTCTCCTGGAGCTTTTTTTACCTACTCTTCTGGACTTGCTCCAATGTCCATTATGTTTCACTTCTCATCTGTTTTCATTCTCTTTCATGTTTCTGTTACTGACTAGAGGTTCTTGATTTCTCATTGCAATAGAAATTGACATGAGGCCAAAAGAATTTTCCCAGACAAGCCTTCATTAAAACTTATGCTTGGGCATAAGGAAGGCAGAGGGTGGGGGGGGGGAAGAGAGAGAGAAAGAGAGAGAGAGAGAGCGAGAGAGAGAATCCCCTGACTGACTCTGAAAAGAGCCAGTAGGGCTTTTTTATTAGACTAAGCAAAGGAAATGACATCAGGGTAGGGTATGCTGGCTGAGGGGTAGGGCATATAGGTCAGCATTATCTGGTCATTAGAGTTATCTTGAGTAATGGGCCACCTGGTGGTCTGGCCAGTGGCAACAAGACTATAAATCAATTGTCCAACATTCCTTACTGAGGGGGGACACTGCAACCTTGCTTATCTCCTAAGGCCAGTTCCTAGAATTATTTAAGTAAAAGGACTATAGCAGTGAGGTAGTAGTGTGGGTTTTATGATCAGTGGGAATACATGAAAGAATGCTCTAGTAGGGGTAAGCTGAAGCCAAGCCCCATCTCTACTCAGTCTCATTTCCAAATACAGAAAATATTTATATTCTATTCTAACATTTCATTAATAATTTGATTGGTAAGGGATTCTAGGTTGGAAATAATTTTTCTTCACAATTTTGCGTTGGTCACTTAACATCCAGCATTTGTGAGATGTCTGAAGCTATTCTGATACTGAAAAAAAAAAAAAAAAAAAAAAAAAAAAAAAAAAAAAAAAAAAAAAAAAAAAAAAAAAAAAAAAAAAAAAAAAAAAAAAAAAAAAAAAAAAAAAAAGAGGAGCAGCTTTGATAATTGTTCTCAATTGGCTATTGTCAACTTCCGATGGCCAGCCGCTATGCTCCTCATCTTCAAGGCTCTCATCTCCTTTGCAAAACTTCTTGAACACCACTGCACTGTACGTTC

>L1_5

AGGCCTCACACATTCAGACACATCCATACTGACACCCACTCCACAGCCCCATGCACCCACACCAGTGAGATGTGGAGGGTCAGACTCCTAAAATTAGGCAGCTGTTGGGGATAAGAGTTGATTTGTTTTTCAATTTTTTTGAAAACAGAAAAGCATGGGGGAATGCATTTGGCCATTACAATGCTAATTGAGTTTGTGTATATTACATATATGGCAGTTAACACTGTAATATTCCTTTTACATTCTATATACACAGAATGATATCAAGGTTTTATGGTCAACAGAATATTCCAACTTCAGTCTTAATGCTGCTTGTAGTGATTTCTGAATTCATTATAGGGGCTTTCCCTAAAAATAATTCAAGTCTATGTTAAGTGAAATAAGGCACAATTAATATTGATTTGATTTAGGGAAAGGGAAGGAAAGAGAAGGGATAAAAACTGGTTTCAAATGATCTTTCTGTTGGGGAACTAGCTCTGACTTAAACCCACCTGAAATTCCTTCTCCTAATTTCCAAAGATTTCTTTATAAAGATATTTTTCTTTTCCCTGACAAAGCCAAAAAGAAAAGTTTGGGAATTCACATTTTAATGTTTCAGTAGCCTGAACAACTCAAATTGATGTGTACCCCCACCTCCCCTGATCAGGTCGCCTCCCTCACCCATGTTAAAATAAAAACACAGTATGTGACACAGGAGTCCCTTGATGTTTCTTGAGGAGATCTGTACACTTGAGTAGCAAATACATATCTGGTTGTCTTCAATATATTTTATTTTTTTTTTTTTTTTTTTTTTTTTTTTTTTTTTTTTTTTTTTTTTTTTTTTTTTTTTTTTTTTTTTTATTATTTTTTTTTTTTTATTATACTCTAAGTTTTAGGGTACATGTGCACATTGTGCAGGTTAGTTACATATGTATACATGTGCCATGCTGGTGCGCTGCACCCACTAATGTGTCATCTAGCATTAGGTATATCTCCCAATGCTATCCCTCCCCCCTCCCCCGACCCCACCACAGTCCCCAGAGTGTGATATTCCCCTTCCTGTGTCCATGTGATCTCATTGTTCAATTCCCACCTATGAGTGAGAATATGCGGTGTTTGGTTTTTTGTTCTTGCGATAGTTTACTGAGAATGATGGTTTCCAATTTCATCCATGTCCCTACAAAGGATATGAACTCATCATTTTTTATGGCTGCATAGTATTCCATGGTGTATATGTGCCACATTTTCTTAATCCAGTCTATCATTGTTGGACATTTGGGTTGGTTCCAAGTCTTTGCTATTGTGAATAGTGCCGCAATAAACATACGTGTGCATGTGTCTTTATAGCAGCATGATTTATACTCATTTGGGTATATACCCAGTAATGGGATGGCTGGGTCAAATGGTATTTCTAGTTCTAGATCCCTGAGGAATCGCCACACTGACTTCCACAATGGTTGAACTAGTTTACAGTCCCACCAACAGTGTAAAAGTGTTCCTATTTCTCCGCATCCTCTCCAGCACCTGTTGTTTCCTGACTTTTTAATGATTGCCATTCTAACTGGTGTGAGATGATATCTCATAGTGGTTTTGATTTGCATTTCTCTGATGGCCAGTGATGATGAGCATTTCTTCATGTGTTTTTTGGCTGCATAAATGTCTTCTTTTGAGAAGTGTCTGTTCATGTCCTTCGCCCACTTTTTGATGGGGTTGTTTGTTTTTTTCTTGTAAATTTGTTTGAGTTCATTGTAGATTCTGGATATTAGCCCTTTGTCAGATGAGTAGGTTGCAAAAATTTTCTCCCATGTTGTAGGTTGCCTGTTCACTCTGATGGTAGTTTCTTTTGCTGTGCAGAAGCTCTTTAGTTTAATTAGATCCCATTTGTCAATTTTGTCTTTTGTTGCCATTGCTTTTGGTGTTTTGGACATGAAGTCCTTGCCCACGCCTATGTCCTGAATGGTAATGCCTAGGTTTTCTTCTAGGGTTTTTATGGTTTTAGGTTTAACGTTTAAATCTTTAATCCATCTTGAATTGATTTTTGTATAAGGTGTAAGGAAGGGATCCAGTTTCAGCTTTCTACATATGGCTAGCCAGTTTTCCCAGCACCATTTATTAAATAGGGAATCCTTTCCCCATTGCTTGTTTTTCTCAGGTTTGTCAAAGATCAGATAGTTGTAGATAAGCGGCATTATTTCTGAGGGCTCTGTTCTGTTCCATTGATCTATATCTCTGTTTTGGTACCAGTACCATGCTGTTTTGGTTACTGTAGCCTTGTAGTATAGTTTGAAGTCAGGTAGTGTGATGCCTCCAGCTTTGTTCTTTTGGCTTAGGATTGACTTGGCAATGCGGGCTCTTTTTTGGTTCCATATGAACTTTAAAGTAGTTTTTTCCAATTCTGTGAAGAAAGTCATTGGTAGCTTGATGGGGATGGCATTGAATCTGTAAATTACCTTGGGCAGTATGGCCATTTTCACGATATTGATTCTTCCTACCCATGAGCATGGAATGTTCTTCCATTTGTTTGTCTCCTCTTTTATTTCCTTGAGCAGTGGTTTGTAGTTCTCCTTGAAGAGGTCCTTCACATCCCTTGTAAGTTGGATTCCTAGGTATTTTATTCTCTTTGAAGCAATTGTGAATGGGAGTTCACCCATGATTTGGCTCTCTGTTTGTCTGTTGTTGGTGTATAAGAATGCTTGTGATTTTTGTACATTGATTTTGTATCCTGAGACTTTGCTGAAGTTGCTTATCAGCTTAAGGAGATTTTGGGCTGAGACGATGGGGTTTTCTAGATAAACAATCATGTCGTCTGCAAACAGGGACAATTTGACTTCCTCTTTTCCTAATTGAATACCCTTTATTTCCTTCTCCTGCCTGATTGCCCTGGCCAGAACTTCCAACACTATGTTGAATAGGAGCGGTGAGAGAGGGCATCCCTGTCTTGTGCCGGTTTTCAAAGGGAATGCTTCCAGTTTTTGCCCATTCAGTATGATATTGGCTGTGGGTTTGTCATAGATAGCTCTTATTATTTTGAAATACGTCCCATCAATACCTAATTTATTGAGAGTTTTTAGCATGAAGGGTTGTTGAATTTTGTCAAAGGCTTTTTCTGCATCTATTGAGATAATCATGTGGTTTTTGTCTTTGGCTCTGTTTATATGCTGGATTACATTTATTGATTTGCGTATATTGAACCAGCCTTGCATCCCAGGGATGAAGCCCACTTGATCATGGTGGATAAGCTTTTTGATGTGCTGCTGGATTCGGTTTGCCAGTATTTTATTGAGGATTTTTGCATCAATGTTCATCAAGGATATTGGTCTAAAATTCTCTTTTTTGGTTGTGTCTCTGCCTGGCTTTGGTATCAGAATGATGCTGGCCTCATAAAATGAGTTAGGGAGGATTCCCTCTTTTTCTATTGATTGGAATAGTTTCAGAAGGAATGGTACCAGTTCCTCCTTGTACCTCTGGTAGAATTCGGCTGTGAATCCATCTGGTCCTGGACTCTTTTTGGTTGGTAAACTATTGATTATTGCCACAATTTCAGAGCCTGTTATTGGTCGATTCAGAGATTCAACTTCTTCCTGGTTTAGTCTTGGGAGAGTGTATGTGTTGAGGAATGTATCCATTTCTTCTAGATTTTCTAGTTTATTTGCGTAGAGGTGTTTGTAGTATTCTCTGATGGTAGTTTGTATTTCTGTGGGATCGGTGGTGATATCCCCTTTATCATTTTTTATTGTGTCTATTTGATTCTTCTCTCTTTTTTTCTTTATTAGTCTTGCTAGCGGTCTATCAATTTTGTTGATCCTTTCAAAAAACCAGCTCCTGGATTCATTGATTTTTTGAAGGGTTTTTTGTGTCTCTATTTCCTTCAGTTCTGCTCTGATTTTAGTTATTTCTTGCCTTCTGCTAGCTTTTGAATGTGTTTGCTCTTGCTTTTCTAGTTCTTTTAATTGTGATGTTAGGGTGTCAATTTTGGATCTTTCCTGCTTTCTCTTGTAGGCATTTAGTGCTATAAATTTCCCTCTACACACTGCTTTGAATGCGTCCCAGAGATTCTGGTATGTGGTGTCTTTGTTCTCGTTGGTTTCAAAGAACATCTTTATTTCTGCCTTCATTTCGTTATGTACCCAGTAGTCATTCAGGAGCAGGTTGTTCAGTTTCCATGTAGTTGAGCGGCTTTGAGTGAGATTCTTAATCCTGAGTTCTAGTTTGATTGCACTGTGGTCTGAGAGATAGTTTGTTATAATTTCTGTTCTTTTACATTTGCTGAGGAGAGCTTTACTTCCAACTATGTGGTCAATTTTGGAATAGGTGTGGTGTGGTGCTGAAAAAAATGTATATTCTGTTGATTTGGGGTGGAGAGTTCTGTAGATGTCTATTAGGTCTGCTTGGTGCAGAGCTGAGTTCAATTCCTGGGTATCCTTGTTGACTTTCTGTCTCGTTGATCTGTCTAATGTTGACAGTGGGGTGTTAAAGTCTCCCATTATTAATGTGTGGGAGTCTAAGTCTCTTTGTAGGTCACTGAGGACTTGCTTTATGAATCTGGGTGCTCCTGTATTGGGTGCATAAATATTTAGGATAGTTAGCTCCTCTTGTTGAATTGATCCCTTTACCATTATGTAATGGCCTTCTTTGTCTCTTTTGATCTTTGTTGGTTTAAAGTCTGTTTTATCAGAGACTAGGATTGCAACCCCTGCCTTTTTTTGTTTTCCATTGGCTTGGTAGATCTTCCTCCATCCTTTTATTTTGAGCCTATGTGTGTCTCTGCACGTGAGATGGGTTTCCTGAATACAGCACACTGATGGGTCTTGACTCTTTATCCAACTTGCCAGTCTGTGTCTTTTAATTGCAGAATTTAGTCCATTTATATTTAAAGTTAATATTGTTATGTGTGAATTTGATCCTGTCATTATGATGTTAGCTGGTGATTTTGCTCATTAGTTGATGCAGTTTCTTCCTAGTCTCGATGGTCTTTACATTTTGGCATGATTTTGCAGCGGCTGGTACCGGTTGTTCCTTTCCATGTTTAGCGCTTCCTTCAGGAGCTCTTTTAGGGCAGGCCTGGTGGTGACAAAATCTCTCAACATTTGCTTGTCTATAAAGTATTTTATTTCTCCTTCACTTATGAAGCTTAGTTTGGCTGGATATGAAATTCTGGGTTGAAAATTCTTTTCTTTAAGAATGTTGAATATTGGCCCCCACTCTCTTCTGGCTTGTAGGGTTTCTGCCGAGAGATCCGCTGTTAGTCTGATGGGCTTTCCTTTGAGGGTAACCCGACCTTTCTCTCTGGCTGCCCTTAACATTTTTTCCTTCATTTCAACTTTGGTGAATCTGACAATTATGTGTCTTGGAGTTGCTCTTCTCGAGGAGTATCTTTGTGGCGTTCTCTGTATTTCCTGAATCTGAACGTTGGCCTGCCTTGCTAGATTGGGGAAGTTCTCCTGGATAATCTCCTGCAGAGTGTTTTCCAACTTGGTTCCATTCTGGTTCCAACAGAAAAGCATGGGGGAATGCATTTGGCCATTACAATGCTAATTGAGTTTGTGTATATTACATATATGGCAGTTAACACTGTAATATTCCTTTTACATTCTATATACACAGAATGATATCAAGGTTTTATGGTCAACAGAATATTCCAACTTCAGTCTTAATGCTGCTTGTAGTGATTTCTGAATTCATTATAGGGGCTTTCCCTAAAAATAATTCAAGTCTATGTTAAGTGAAATAAGGCACAATTAATATTGATTTGATTTAGGGAAAGGGAAGGAAAGAGAAGGGATAAAAACTGGTTTCAAATGATCTTTCTGTTGGGGAACTAGCTCTGACTTAAACCCACCTGAAATTCCTTCTCCTAATTTCCAAAGATTTCTTTATAAAGATATTTTTCTTTTCCCTGACAAAGCCAAAAAGAAAAGTTTGGGAATTCACATTTTAATGTTTCAGTAGCCTGAACAACTCAAATTGATGTGTACCCCCACCTCCCCTGATCAGGTCGCCTCCCTCACCCATGTTAAAATAAAAACACAGTATGTGACACAGGAGTCCCTTGATGTTTCTTGAGGAGATCTGTACACTTGAGTAGCAAATACATATCTGGTTGTCTTCAATATATTTTAAAAAGAAAGAAAGAGAAAGAAGAAAGGAAAGAAGGAAAAAAGGAAGGAAAGAAGTAATATTGTGACTTTCTTGCTCTTTAAAAAAATCCCTGTCTCCTTCTTTTCAGGATGATGAAGCCCCACTAGACATTACAAACAACTGCAACAAATGAGTTTGGCAGCATAAATAATGTCTAGTTATTCAAATCTTCATAGTGAGATTAATTTATCTTGTACACAATTT

>L1_6_Orphan_L1

TTAGGTTGTATCATCCTCATTTTCAGAAGACGAAGTTGAAACTTAATAACTTAGGGATTATGTCAGTTGTTTTGTATTCTCTATACTTCCTAGTCCATAGGCAGTGTTCATAAAAGCTAGGTGAGTTAAAAAGAAAAAAGTATACACGGAGCTAAATTTAAGATGAAATAATTTTTTCTTTATTCCCATCTCTTTAAACTTCAAGAAAGATTAACAATATGAAAGGTTGTATTTGAATTCCAGGATTCAAAACAGTATTTCACATAGATCTTTAAACTTAAAAATAAAATGACTTTAAAGGCTAATATGTCCAATAGATGGTCATCTCTAGAGAATATAATCCTTCAGAGGTTTTCCAACACTTTATTAATATGCTTCTTCTGGGAACTTGCCTGTGATCCATTTTTTTTCTGGAGTCTTAGCAAAATACTTTTCAATGAATTATCCTTTGCAAGAGAATTATCTTTAAAACAATGCAGAGCTAACCAAACTAACAAAATAAACTTTATGTGCAAAAGGCATCTTTGAATAAATCATGATTTAAATGTCAAAAAAAAAAAAAAAAAAAAAAAAAAAAAAAAAAAAAAAAAAAAAAAAAAAAAAAAAAAATAACTTGCCTACGGGCACAGGCTGTTATAGAATTTGGC

>L1_7

TCAGGGTGCACAGTAAGAACTTTTAAAAGTTAATCTAGAGGGAGGAGCCAAGATGGCCGAATAGGAACAGCTCCGGTCTACAGCTCCCAGCGTGAGCGACGCAGAAGACGGTGATTTCTGCATTTCCATCTGAGGTACCGGGTTCA*TCTCACTAGGGAGTGCCAGACAGTGGGCGCAGGCCAGTGTGTGTGCGCACCGTGCGCGAGCCGAAGCNNNNNNNNNNNNNNNNNNNNNNNNNNNNNNNNNNNNNNNNNNNNNNNNNNNNNNNNNNNNNNNNNNNNNNNNNNNNNNNNNNNNNNNNNNNNNNNNNNNNNNNNNNNNNNNNNNNNNNNNNNNNNNNNNNNNNNNNNNNNNNNNNNNNNNNNNNNNNNNNNNNNNNNNNNNNNNNNNNNNNNNNNNNNNNNNNNNNNNNNNNNNNNNNNNNNNNNNNNNNNNNNNNNNNNNNNNNNNNNNNNNNNNNNNNNNNNNNNNNNNNNNNNNNNNNNNNNNNNNNNNNNNNNNNNNNNNNNNNNNNNNNNNNNNNNNNNNNNNNNNNNNNNNNNNNNNNNNNNNNNNNNNNNNNNNNNNNNNNNNNNNNNNNNNNNNNNNNNNNNNNNNNNNNNNNNNNNNNNNNNNNNNNNNNNNNNNNNNNNNNNNNNNNNNNNNNNNNNNNNNNNNNNNNNNNNNNNNNNNNNNNNNNNNNNNNNNNNNNNNNNNNNNNNNNNNNNNNNNNNNNNNNNNNNNNNNNNNNNNNNNNNNNNNNNNNNNNNNNNNNNNNNNNNNNNNNNNNNNNNNNNNNNNNNNNNNNNNNNNNNNNNNNNNNNNNNNNNNNNNNNNNNNNNNNNNNNNNNNNNNNNNNNNNNNNNNNNNNNNNNNNNNNNNNNNNNNNNNNNNNNNNNNNNNNNNNNNNNNNNNNNNNNNNNNNNNNNNNNNNNNNNNNNNNNNNNNNNNNNNNNNNNNNNNNNNNNNNNNNNNNNNNNNNNNNNNNNNNNNNNNNNNNNNNNNNNNNNNNNNNNNNNNNNNNNNNNNNNNNNNNNNNNNNNNNNNNNNNNNNNNNNNNNNNNNNNNNNNNNNNNNNNNNNNNNNNNNNNNNNNNNNNNNNNNNNNNNNNNNNNNNNNNNNNNNNNNNNNNNNNNNNNNNNNNNNNNNNNNNNNNNNNNNNNNNNNNNNNNNNNNNNNNNNNNNNNNNNNNNNNNNNNNNNNNNNNNNNNNNNNNNNNNNNNNNNNNNNNNNNNNNNNNNNNNNNNNNNNNNNNNNNNNNNNNNNNNNNNNNNNNNNNNNNNNNNNNNNNNNNNNNNNNNNNNNNNNNNNNNNNNNNNNNNNNNNNNNNNNNNNNNNNNNNNNNNNNNTGCAGCGCACCAGCATGGCACATGTATACATATGTAACTAACCTGCACAATGTGCACATGTACCCTAAAACTTAGAGTATAATAAAAAAAAAAAAAAAAAAAAAAAAAAAAAAAAAAAAAAAAAAAAAAAAAAAAAAAAAAAAAAAAAAGTTAATCTAAGTTACAATCAGAGGTATATATGTGGGCTAGCACTGTTAGATTGAAATCTAACTTGACTCTCTTATTTGGTGACATGTTGAATATGCAAAATGA*

>L1_8

*CCTGTCAAGTCTCTGCTCCAGCACTGCACAGTGGGTAAATGTGTGTTGGTGGTAGGGGAGGGAGTGAGAGCCTATCTTTTCAGCTCATACGTTTCTAGTTCAAGACGAATTACATCTGACCTGACGTAGAAATAAATGTTTATTACCCACTTGTGTTAGTTTCTTTTTTTTTTTTTTTTTTTTTTTTTTTTTTTTTTTTTTTTTTTTTTTTTTTTTTTTTTTTTTTTTNNNNNNNNNNNNNNNNNNNNNNNNNNNNNNNNNNNNNNNNNNNNNNNNNNNNNNNNNNNNNNNNNNNNNNNNNNNNNNNNNNNNNNNNNNNNNNNNNNNNNNNNNNNNNNNNNNNNNNNNNNNNNNNNNNNNNNNNNNNNNNNNNNNNNNNNNNNNNNNNNNNNNNNNNNNNNNNNNNNNNNNNNNNNNNNNNNNNNNNNNNNNNNNNNNNNNNNNNNNNNNNNNNNNNNNNNNNNNNNNNNNNNNNNNNNNNNNNNNNNNNNNNNNNNNNNNNNNNNNNNNNNNNNNNNNN*ATGCGGCATTATTTCTGAGGGCTCTGTTCTGTTCCATTGATCTATATCTCTGTTTTGGTACCAGTACCATGCTGTTTTGGTTACCGTAGCCTTGTAGTATAGTTTGAAGTCAGGTAGTGTGATGCCTCCAGCTTTGTTCTTTTGTTAGTTTCTTAAGGCTGCTGTGACAAATCACCACAAAC

>SVA_1

ATCTGATTTGGCAAATATTTTTTTTTTTTTTTTTTTTTTTTTTTTTTTTTTTTTTTTTTTTTTTTTTTTTTTTTTTTTTTTTTTTTTTTTTTTTTTTTTTTTTTTTTTTTTTTTTTTTTTTTTTTTTTTTTTTTTTTTTTTTTTGTTTTTCTTTTTATTTATTTATTTATTTATTTATTTATTTTTTATTGATCATTCTTGGGTGTTTCTCGCAGAGGGGGATTTGGCAGGGTCATAGGACAATAGTGGAGGGAAGGTCAGCAGATAAACAAGTGAACAAAGGTCTCTGGTTTTCCTAGGCAGAGTGTGTGTGTCCCTGGGTACTTGAGATTAGGGAGTGGTGATGACTCTTAACGAGCATGCTGCCTTCAAGCATCTGTTTAACAAAGCACATCTTGCACCGCCCTTAATCCATTTAACCCTGAGTGGACACAGCACATGTTTCAGAGAGCACAGGGTTGGGGGTAAGGTCATAGATCAACAGGATCCCAAGGCAGAAGAATTTTTCTTAGTACAGAACAAAATGAAAAGTCTCCCATGTCTACTTCTTTCTACACAGACACAGCAACTATCCGATTTCTCAATCTTTTCCCCACCTTTCCCCCTTTTCTATTCCACAAAACCGCCATTGTCATCATGGCCCGTTCTCAATGAGCTGTTGGGTACACCTCCCAGACGGGGTGGTGGCCGGGCAGAGGGGCTCCTCACTTCCCAGTAGGGGCGGCCGGGCAGAGGCGCCCCTCACCTCCCGGACGGGGCGGCTGGCCGGGCAGGGGGCTGACCCCCCACCTCCCTCCCGGACGAGGCGTCTCGCCTGGCGGGGGGCTGACCCCCCCACCTCCCTCCCAGACGGGGCGGCTGGCCGGGCGGGGGGCTGACCCCCCCACCTCCCTCCCGGACGGGGTGGCTGGCCGGGCGGGGGGCTGACCCCCCCACCTCCCTCCCGGACGGGGCGGCTGGCCGGGCGGGGGGGCTGACCCCCCCCCACCTCCCTCCCGGACGTGGCGGCTGGCCGGGCAGAGGGGCTCCTCACTTCCCAGTAGGGGCGGCCGGGCAGAGGCGCCCCTCACCTCCCAGACAGGGCGGCTGGCTGGGCGGGGGGCTGACCCCCTCACCTCCCTCCCGGACGGGGCGGCTGGCCGGGCAGAGGGGCTCCTCGCTTCCCAGTAGGGGCGGCCGGGCAGAGGCGCCCCTCACCTCCCGGACGGGGCGGCTGGCTGGGCGGGGGGCTGACCCCCCCACCTCCCTCCCGGACGGGGCGGCTGGCCGGGCGGGGGGCTGACCCCCCCACCTCCCTCCCGGACGGGGCGGCTGGCCGGGCGGGGGGCTGACTCCCCCACCTCCCTCCCGGATGGGGCGGCTGGCCGAGCAGAGGGGCTCCTCACTTCCCAGTAGGGGCGGCCGGGCAGAGGCGCCCTCACCTCCCGGACGGGGTGGCTGGCCGGGTGGGGGGCTGACCCCCCACCTCCCTCCCGGACGGGGCGGCTGGCCTGGCGGGGGCTGACCCCCACCTCCCTTCCGGACGGGGTGGCTGCCGGGCGGAGACGCTCCTCACTTCCCAGACGGGGTGGCAGCCGGGCGGAGGAGCTCCTCACTTCTCAGATGGGGCAGTTGCCAGGTGGAGGGTCTCCTCACTTCTCAGACGGGGTGGCCGGGCAGAGACGCTCCTCACCTCCCAGACGGGGTCGCGGCCGGGCCGAGGTGCTCCTCACATCCCAGACGGGGCGGCGGGGCAGAGGCGCTCCCCACATCTCAGAGGATGGGCGGCCGGGCAGAGACGCTCCTCACTTCCTCAGCCTCCCAAGTAGCTGGGAATACAGGCGCCCGCCACCATGCCAGTTTAATTCTCGTATTTTTAGTAGAGATACAGTTTCACCATGTTGGCCAGGCTGGCAAACTCCTGACCTCAGGTGATCTGCCCGCCTGGACCTCCCAAAGTGCTGGGATTACAGGCATGAGCCACTGTGCCTGGCCAGCCTCACCTTTCTCTTTTGGCAAATATTTTTAAAGATGATATTTGAATGAGAAAATTGGCATTTGGGACATTCTTAAACTAAATTTGAGACATCTTAGGCAAAACAAATACTTATTTTTAAGGCACTATTGTTATGGCACTGAAGTCTTGGAACTATTTGATCTAGTTACTGTAAGTTCTCAGCTGTGTTGCAACTCATTAAAGAGAATATTGTTATTAAAGGTATTTGCAAGAAAAACTTAGAGATACTATAGTATCTCCTTTCTCTGTCTCAAACTTTTTTCCCCTCAATACCCAAGGCTCTGTGATGTCTCAAATTTTAATCATTACTTTAAAAAGAGAAGTTTAAAGCATTAAAGAATTATAATCAGATGAAAGCAGCTTTGGATTTATAAAATTCTGAAACAATAATTTTAATTTTGCTTTTAACATATATGCAAATTCTTTGATACTCTCCACTTTGCAGAGGTGCAGGTTCATTCCCTCCCTGTGA

>SVA_2

TGCTTAAGAAAACAAAGCTGCATGCCCTCTCCCTCTCCCTCTCCCTCTCCCTCTCCCTCTCCCTCTCCCTCCCCCTCCCTCTCCCTCTCCCTCTCCCTCTACCTCCACGGTCTCCCTCTGATGCCGAGCCAAGGCTGGACGGTACTG*CTGCCATCTCGGCTCACTGCAACCTCCCTGCCTGATTCTCCTGCCTCAGCCTGCCGAGTGCCTGCGATTGCAGGCGCGCACCGCCACGCCTGACTGGTTTTCGTTTTTTTTTGGTGGAGACGGGGTTTCGCTGTGTTGGCCGGGCTGGTCTCCAGCTCCTAGCCGCGAGTGATCCGCCAGCCTCGGCCTCCCGA*NNNNNNNNNNNNNNNNNNNNNNNNNNNNNNNNNNNNNNNNNNNNNNNNNNNNNNNNNNNNNNNNNNNNNNNNNNNNNNNNNNNNNNNNNNNNNNNNNNNNNNNNNNNNNNNNNNNNNNNNNNNNNNNNNNNNNNNNNNNNNNNNNNNNNNNNNNNNNNNNNNNNNNNNNNNNNNNNNNNNNNNNNNNNNNNNNNNNNNNNNNNNNNNNNNNNNNNNNNNNNNNNNNNNNNNNNNNNNNNNNNNNNNNNNNNNNNNNNNNNNNNNNNNNNNNNNNNNNNNNNNNNNNNNNNNNNNNNNNNNNNNNNNNNNNNNNNNNNNNNNNNNNNNNNNNNNNNNNNNNNNNNNNNNNNNNNNNNNNNNNNNNNNNNNNNNNNNNNNNNNNNNNNNNNNNNNNNNNNNNNNNNNNNNNNNNNNNNNNNNNNNNNNNNNNNNNNNNNNNNNNNNNNNNNNNNNNNNNNNNNNNNNNNNNNNNNNNNNNNNNNNNNNNNNNNNNNNNNNNNNNNNNNNNNNNNNNNNNNNNNNNNNNNNNNNNNNNNNNNNNNNNNNNNNNNNNNNNNNNNNNNNNNNNNNNNNNNNNNNNNNNNNNNNNNNNNNNNNNNNNNNNNNNNNNNNNNNNNNNNNNNNNNNNNNNNNNNNNNNNNNNNNNNNNNNNNNNNNNNNNNNNNNNNNNNNNNNNNNNNNNNNNNNNNNNNNNNNNNNNNNNNNNNNNNNNNNNNNNNNNNNNNNNNNNNNNNNNNNNNNNNNNNNNNNNNNNNNNNNNNNNNNNNNNNNNNNNNNNNNNNNNNNNNNNNNNNNNNNNNNNNNNNNNNNNNNNNNNNNNNNNN*TGTTCACTTGTTTATCTGCTGACCTTCCCTCCACTATTGTCCTATGACCCTGCCAAATC*CCCCTCTGCGAGAAACACCCAAGAATGATCAATAAAAAAAAAATAAATTAATTAAAAAAAAAAAAAAAAAAAAAAAAAAAAAAAAAAAAAAAAAAAAAGAAAACAAAGCTGCATTTGGGAGGCTGAGGCGG

>SVA_3

*TGCTTGGCCTCCCTCCTATATTTTCTTTTTTTTTTAAATTATACTTTAAGTTCTAGGGTACATGTGAACAACGTGCAGGTTTGTTACATATGCTTACATGTGCCATGTTGGTGTGCTGCACCCATTAACTCATAATTTAGCATTAGGTATATCTCCTAATGCGATCCCTCCTCCCTCCCCACAACCCACAACAGGCCCCAGTGTGTGATGTTCCCCACCCTGTGTCCGAGTGTTCTCATTGTTCAATTCCCACCTATGAGTGAGAACATGCGGTGTTTGGTTTTCTGTCCTTGCGATAGTTTGCTGAGAATGATGGTTTCCAGCTTCATCCATGTCCCTACAAAGGACATGAACTCATTCTTTTTTATGGCTGCATAGTATTCCTGGAATACTATATTTTCTTTTTTTTTTTTTTTTTTTTTTTTTTTTTTTTTTTTTTTTTTTTTTTTTTTTTTTTTTTTTTTTTTTTTTTTTTTTTTTTTTTTTTTTTTTTTTTTTTTTTTTTTTTTTTTTTTTTTTTTTTTTTTTTTTTTTTTTTTTTTTTTTTTTTTT*NNNNNNNNNNNNNNNNNNNNNNNNNNNNNNNNNNNNNNNNNNNNNNNNNNNNNNNNNNNNNNNNNNNNNNNNNNNNNNNNNNNNNNNNNNNNNNNNNNNNNNNNNNNNNNNNNNNNNNNNNNNNNNNNNNNNNNNNNNNNNNNNNNNNNNNNNNNNNNNNNNNNNNNNNNNNNNNNNNNNNNNNNNNNNNNNNNNNNNNNNNNNNNNNNNNNNNNNNNNNNNNNNNNNNNNNNNNNNNNNNNNNNNNNNNNNNNNNNNNNNNNNNNNNNNNNNNNNNNNNNNNNNNNNNNNNNNNNNNNNNNNNNNNNNNNNNNNNNNNNNNNNNNNNNNNNNNNNNNNNNNNNNNNNNNNNNNNNNNNNNNNNNNNNNNNNNNNNNNNNNNNNNNNNNNNNNNNNNNNNNNNNNNNNNNNNNNNNNNNNNNNNNNNNNNNNNNNNNNNNNNNNNNNNNNNNNNNNNNNNNNNNNNNNNNNNNNNNNNNNNNNNNNNNNNNNNNNNNNNNNNNNNNNNNNNNNNNNNNNNNNNNNNNNNNNNNNNNNNNNNNNNNNNNNNNNNNNNNNNNNNNNNNNNNNNNNNNNNNNNNNNNNNNNNNNNNNNNNNNNNNNNNNNNNNNNNNNNNNNNNNNNNNNNNNNNNNNNNNNNNNNNNNNNNNNNNNNNNNNNNNNNNNNNNNNNNNNNNNNNNNNNNNNNNNNNNNNNNNNNNNNNNNNNNNNNNNNNNNNNNNNNNNNNNNNNNNNNNNNNNNNNNNNNNNNNNNNNNNNNNNNNNNNNNNNNNNNNNNNNNNNNNNNNNNNNNNNNNNNNNNNNNNNNNNNNNNNNNNNNNNNN*CGGAGACGCTCCTCACTTCCCAGATGGGGTGGCTGCCGGGCGGAGAGGCTCCTCACTTCTCAGACGGGGTGGTTGCCAGGCAGAGGGTCTCCTCACTTCTCAGACGGGGCAGCCGGGCAGAGACGCTCCTCA*CCTCCCAGATGGGGTCTCGGCCGGGCAGAGGCGCTCCTCACATCCCAGATGGGGCGGCGGGGCAGAGGCGCTCCCCACATCTCAGACGATGGGCGGCCGGGCAGAGACGCTCCTCACTTCCCAGATCTGGAATACTATATTTTCAATTCTTCTCTTTAGTGGTTCTTTCCCTTTGCCAATT

SVA_4

GTGTTCAAATAAAACAGAGGCCAGCCTGGCGAACATGGTGAAACCCCATCTCTACTAAAAATACAAAAATTAGCCGGGCATGGTGGCACATGCCTGTAATCCCAGCTACTCGGGAGGCTGAGGCAGGAGAATTGCTTGAACCCAGGAGGCAGAGGTTGCAGTGAGCTGAGATTGCACCACTGCACTCTAGCCTAGGCAACAGAGTGAATGAGACTCCATTTCAAAAAAAACAAAAAATTATCTATCTATCTATCTATCTATCTATCTATCTATCTATCTACCTACCTACCTATCTATCTATCTATCTATCTACCTATCTATCTATCTAAAACTGAGGGTCGGCCGCGCCGGCGAGCGCCGCCCGGGAGGCAGCGGCTGGAGGAGCGGACGGGCCCCGCGGGGCCCGAGGGCAAGGAGCAGCCGCCTGCCTTGGCCTCCCAAAGTGCCGAGATTGCAGCCTCTGCCCGGCTGCCACCCCGTCTGGGAAGTGAGGAGTGTCTCTGCCTGGCCACCCATCGTCTGGGATGTGAGGAGCCCCTCTGCCTGGCTGCCCAGTCTGGAAAGTGAGGAGCGTCTCCGCCCGGCCGCCATCCCATCTAGGAAGTGAGGAGCGCCTCTTCCCAGCCGCCATCACATCTAGGAAGT*GAGGAGGGTCTCTGCCCGGCCGCCCATCGTCTGAGCTGTGGGGAGCGCCTCTGCCCCGCCNNNNNNNNNNNNNNNNNNNNNNNNNNNNNNNNNNNNNNNNNNNNNNNNNNNNNNNNNNNNNNNNNNNNNNNNNNNNNNNNNNNNNNNNNNNNNNNNNNNNNNNNNNNNNNNNNNNNNNNNNNNNNNNNNNNNNNNNNNNNNNNNNNNNNNNNNNNNNNNNNNNNNNNNNNNNNNNNNNNNNNNNNNNNNNNNNNNNNNNNNNNNNNNNNNNNNNNNNNNNNNNNNNNNNNNNNNNNNNNNNNNNNNNNNNNNNNNNNNNNNNNNNNNNNNNNNNNNNNNNNNNNNNNNNNNNNNNNNNNNNNNNNNNNNNNNNNNNNNNNNNNNNNNNNNNNNNNNNNNNNNNNNNNNNNNNNNNNNNNNNNNNNNNNNNNNNNNNNNNNNNNNNNNNNNNNNNNNNNNNNNNNNNNNNNNNNNNNNNNNNNNNNNNNNNNNNNNNNNNNNNNNNNNNNNNNNNNNNNNNNNNNNNNNNNNNNNNNNNNNNNNNNNNNNNNNNNNNNNNNNNNNNNNNNNNNNNNNNNNNNNNNNNNN*ATGGATTAAGGGCGGTGCAAGATGTGCTTTGTTAAACAGATGCTTGAAGGCAGCATGCTCGTTAAGAGTCATCACCAATCCCTAATCTCAAGTAATCAGGGACACAAACACTGCGGAAGGCCGCAGGGTCCTCTGCCTAGGAAAACCAGAGACCTTTGTTCACTTGTTTATCTGCTGACCTTCCCTCCACTATTGTCCCATGACCCTGCCAAATCCCCCTCTGTGAGAAACACCCAAGAATTATCAATAAAAAAATAAATTTAAAAAAAAAAAAAAAAAAAAAAAAAAAAAAAAAAAAAAAAAAAAAAAAAAAAAAAAAAAAAAAAAAAAAAAAAAAAAAAAAAAAAAAAAAAAAAAAAAAAAAAAAAAAAAAAAAATAAAACTGAGGGTCAAAAATAGCCTCCACGGTGAAATAGAACATGAAGTCATTTTTGTGCCT

>SVA_5

CTAAGGTGGCTTGCCTCTTGCGGAGATTACCGTGAACACGCTTTTTGAGATTTTTCTGCCTGGTTTAAATTTTTTTTGTTGTTTCCTCCCTCCTGTCAAAGGGCAATTCTTGCTTGTATGCAACTCTTTCTCCTAATTCCCATTTGTTTCAGTGTGGGAAGATAAACTTAGAAAATTTTGTGTCAGGTCGGTGGCTTAG*GAGCCCGTCCCGCCATGGTGGCCGCGGCTGGTGGTTGGCGCGGCTGCGCTGCGGCCCGGGGCAGTGCGGAGCCAGGACAGTCGCGGCGCTGACGCCCGCGGGCCCCAGCTGCAGATATGAAGCGGAGCCGCTGCCGCGACCGACCGCAGCCGCCGCCGCCCGACCGCCGGGAGGATGGAGTTCAGCGGGCAGCGGAGCTGTCTCAGTCTTTGCCGTCGCGCCGGCGAGCGCCGCCCGGGAGGCAGCGGCTGGAGGAGCGGACGGGCCCC*NNNNNNNNNNNNNNNNNNNNNNNNNNNNNNNNNNNNNNNNNNNNNNNNNNNNNNNNNNNNNNNNNNNNNNNNNNNNNNNNNNNNNNNNNNNNNNNNNNNNNNNNNNNNNNNNNNNNNNNNNNNNNNNNNNNNNNNNNNNNNNNNNNNNNNNNNNNNNNNNNNNNNNNNNNNNNNNNNNNNNNNNNNNNNNNNNNNNNNNNNNNNNNNNNNNNNNNNNNNNNNNNNNNNNNNNNNNNNNNNNNNNNNNNNNNNNNNNNNNNNNNNNNNNNNNNNNNNNNNNNNNNNNNNNNNNNNNNNNNNNNNNNNNNNNNNNNNNNNNNNNNNNNNNNNNNNNNNNNNNNNNNNNNNNNNNNNNNNNNNNNNNNNNNNNNNNNNNNNNNNNNNNNNNNNNNNNNNNNNNNNNNNNNNNNNNNNNNNNNNNNNNNNNNNNNNNNNNNNNNNNNNNNNNNNNNNNNNNNNNNNNNNNNNNNNNNNNNNNNNNNNNNNNNNNNNNNNNNNNNNNNNNNNNNNNNNNNNNNNNNNNNNNNNNNNNNNNNNNNNNNNNNNNNNNNNNNNNNNNNNNNNNNNNNNNNNNNNNNNNNNNNNNNNNNNNNNNNNNNNNNNNNNNNNNNNNNNNNNNNNNNNNNNNNNNNNNNNNNNNNNNNNNNNNNNNNNNNNNNNNNNNNNNNNNNNNNNNNNNNNNNNNNNNNNNNNNNNNNNNNNNNNNNNNNNNNNNNNNNNNNNNNNNNNNNNNNNNNNNNNNNNNNNNNNNNNNNNNNNNNNNNNNNNNNNNNNNNNNNNNNNNNNNNNNNNNNNNNNNNNNNNNNNNNNNNNNNNNNNNNNNNCTGACCTTCCCTCCACTATTGTCCCATGACCCTGCCAAATCCCCCTCTGTGAGAAACACCCAAGAATTATCAATAAAAAAATAAATTAAAAAAAAAAAAAAAAAAAGAAAAAAAAGAATGGACTTTCCCAGGCCAGCTGTGGTGGCTCACGACTGTAATCCCAGCACTGTGGCAGGCCAAGGTGGGCAGATCACCTGAGATCAGGAGTTCAAGACCAGCCTGACCAACACGGAGAAACCCCGTCTCTACTAAAAATAAAAAAAATTAGCTGGGCGTGGTGGTGCATGCCTGTAATCCCAGCTACTTGGGAGGCTGAGGCAGGAGAATTGCTTGAACCCAGGAGGCAGAGGTTGTTGTGAGCTGAGATTGCACCATTGCACTCCAGCCTGGGCAACAAGAGGGAAACTCCATCAAAAAAAAAAAAAAAAAGGACTTTCTCAAAGAAAATGTATTTAAATGTCTGCACCAATAATTCCAGCATGTGTATGAATAAATATGATATGTCCTTTACAGTGAAGGTCTAATAAGATTTACTTATATGCCTTTTCCTTCTTAGAAAGTCTCTAAGAAATAAAATATCTTTACAATCAAAAAAAAAAAAAAAAAAAAAAAAAAAAAAAAAAAAAAAAAAAAAAAAAAAAAAAAAAAAAAGAAAATTTTGTGTCACAAGGTATATAGGTAATAGTGATTACTTTTGATAAGAGAAAGTTCCTTTTCACTTAATTTTTGTTCTGGGTATATGAATTACCTATTGAAAAATATATATAAGTGAGCATAGATGATTCTTTAAAATAAAAATGTGTTTTACTCTGAATTCCCATCTAACAATTGTGTCACGAAAATGAGTTAGTATAAATAAGAACTTAAGGGTCTGTTTATAGAGAATGTTGGCTTTAGTATATGGGCT

>SVA_6

TACTGGAAAAAGAAGAAAATAGATTATGTTTCTTTTTTTTTTTTTTTTTTTTTTTTTTTTTTTTTTTTTAAATTTATTTTTTTATTGATAATTCTTGGGTGTTTCTCACAGAGGGGGATTTGGCAGGGTCATGGGACAATAGTGGAGGGAAGGTCAGCAGATAAACAAGTGAACAAAGGTCTCTGGTTTTCCTAGGCAGAGGACCCTGCGGCCTTCCGCAGTGTTTGTGTCCCTGATTACTTGAGATTAGGGATTGGTGATGACTCTTAACGAGCATGCTGCCTTCAAGCATCTGTTTAACAAAGCACATCTTGCACCGCCCTTAATCCATTTAACCCTGAGTGGACACAGCACATGTTNNNNNNNNNNNNNNNNNNNNNNNNNNNNNNNNNNNNNNNNNNNNNNNNNNNNNNNNNNNNNNNNNNNNNNNNNNNNNNNNNNNNNNNNNNNNNNNNNNNNNNNNNNNNNNNNNNNNNNNNNNNNNNNNNNNNNNNNNNNNNNNNNNNNNNNNNNNNNNNNNNNNNNNNNNNNNNNNNNNNNNNNNNNNNNNNNNNNNNNNNNNNNNNNNNNNNNNNNNNNNNNNNNNNNNNNNNNNNNNNNNNNNNNNNNNNNNNNNNNNNNNNNNNNNNNNNNNNNNNNNNNNNNNNNNNNNNNNNNNNNNNNNNNNNNNNNNNNNNNNNNNNNNNNNNNNNNNNNNNNNNNNNNNNNNNNNNNNNNNNNNNNNNNNNNNNNNNNNNNNNNNNNNNNNNNNNNNNNNNNNNNNNNNNNNNN*GCGGCGGCAAAGACTGAGACAGCTCCGCTGCCCGCTGAACTCCATCCTCCTGGCGGTCGGGCGGCGGCGGCTGCGGTCGGTCGCGGCAGCGGCTCCGCTTCATATCTGCAGCTGGGGCCCGCGGGCGTCAGCGCCGCCGCGCCAACCACCAGCCGCGGCCACCATGGCCAGACGGGCTCCCTAAGCCACCGACCCCAGCCCGCGGCGCCTTCGACCCTTCTGGGGCCTCCGGCGCCGCGACCTCCTCTGCCTGAAATTTCTTTTTTCTTTTCCTTTTATTTTATTTTATTTTTTGAGACGGAGTCTTGCTCTGTTGTC*TGGGTGGAGTGCAGTGGTGCAATCTCGGCTCACTGCAACCTCTGCCTCCGATTATGTTTCAATAAACCCATAAGGATACCACTTATGTGAACAAAGCCTCCATTTTTATTAATTCAACTTGGTTTGTGGTTCAAACCCTTAAGATCAGTCCATTAGACACCAGCTCATATATTCGGGTAAAAAAAGAGAATTTCTTTAGCTATTTACATTTATCATATTTATACCACTCCGCAGCATACAGCTGAGGTAAAAAGCAGCAGGCTTGCCATGCCATTGCCGCTTCCGAGAGGCCCCCTTCAAACAGAAGGCACTCTCTGCGGCTCCTCCTCAGACACGATGAAGCCGCCATTGTTGCCAGGGTTTCCTTCC

>SVA_7

AATAAGTAACATAGCAGGGCGCGGTGGCTCATGCCTGTAATCCCTCACGCCTACAATCCCTCACGCCTGTAATCTCAGCAGTTTGGGAGGCCGAGGTGGACAGATTGCTTGCGGTAAGGAGTTTGAGACCAGCGTAGCCAACATGGCGAAACCCCATCTCTACGAAAAATTAGCCAGGCATGGTGGTGAGTGCCTGTAGTCCCAGCTACTTCGGAGGTGGAGGCCGAAGAATTGCTTGAGCCTGGAAGGCTGAGATCTCAGTGAGCCGAGACTGCCACTGCACGGGTGACAGAGCGAGCCTCTGTCTCAAAGAAAGAAAGAGTATGGAGCTCCTAACCGCGAGTGATCCGCCAGCCTCGGCCTCCCGAGGTGCCGGGATTGCAGATGGAGTCTCGTTCACTCAGTGCTCAATGGTGCCCAGGCTGGAGTGCAGTGGCGTGATCTCGGCTCGCTACAA*CACCTCCCAGCCGCCTGCCTTGGCCTCCCAAAGAGCCGACATTGCAGCCTCTGCCCGGCCGCCACCCCGTCTGGGAAGTGAGGAGCGT*NNNNNNNNNNNNNNNNNNNNNNNNNNNNNNNNNNNNNNNNNNNNNNNNNNNNNNNNNNNNNNNNNNNNNNNNNNNNNNNNNNNNNNNNNNNNNNNNNNNNNNNNNNNNNNNNNNNNNNNNNNNNNNNNNNNNNNNNNNNNNNNNNNNNNNNNNNNNNNNNNNNNNNNNNNNNNNNNNNNNNNNNNNNNNNNNNNNNNNNNNNNNNNNNNNNNNNNNNNNNNNNNNNNNNNNNNNNNNNNNNNNNNNNNNNNNNNNNNNNNNNNNNNNNNNNNNNNNNNNNNNNNNNNNNNNNNNNNNNNNNNNNNNNNNNNNNNNNNNNNNNNNNNNNNNNNNNNNNNNNNNNNNNNNNNNNNNNNNNNNNNNNNNNNNNNNNNNNNNNNNNNNNNNNNNNNNNNNNNNNNNNNNNNNNNNNNNNNNNNNNNNNNNNNNNNNNNNNNNNNNNNNNNNNNNNNNNNNNNNNNNNNNNNNNNNNNNNNNNNNNNNNNNNNNNNNNNNNNNNNNNNNNNNNNNNNNNNNNNNNNNNNNNNNNNNNNNNNNNNNNNNNNNNNNNNNNNNNNNNNNNNNNNNNNNNNNNNNNNNNNNNNNNNNNNNNNNNNNNNNNNNNNNNNNNNNNNNNNNNNNNNNNNNNNNNNNNNNNNNNNNNNNNNNNNNNNNNNNNNNNNNNNNNNNNNNNNNNNNNNNNNNNNNNNNNNNNNNNNNNNNNNNNNNNNNNNNNNNNNNNNNNNNNNNNNNNNNNNNNNNNNNNNNNNNNNNNNNNNNNNNNNNNNNNNNNNNNNNNNNNNNN*CCTCTGCCTAGGAAAACCAGAGACCTTTGTTCACTTGTTTATCTGCTGACCTTCCCTCCACTATTGTCCTATGACCCTGCCAAATCCC*CCTCTGCGAGAAACACCCAAGAATGATCAATAAAAATAAAAAATAAAAAATAAAAAATAAAAAAAAAAAAAAAAAAAAAAAAAAAAAAAAAAAAAAAAAAAAAAAAAAAAAAAAAAAAAAAAAAAAAAAAAAAAAAAAAAAAAAAAGAAAGAGTATGTACCTGGATATGTATTTTTAAAATAA

>SVA_8

CAGAAGATCTGTAGTGGGGCCCTGGCATTAATGTGATCTTGATCCAAAACTTATATACTTACAGAAACTATGACACCCTCTGCATACATTGCCTGAGTTACTTGAAAACAGTATCTTAGTAAATGTTTAAGGTTAATATGAACTTTACTACTTGACCAATACCATATGTTATGATTTACATGAAAAAACTAGAGGCAGAGGAGGTCGCGGCGCCGGAGGCCCCAGAAGGGTCGAAGGCGCCGCGGGCTGGGGTCGGTGGCTTAGGGAGCCCGTCTGGCCATGGTGGCCGCGGCTGGTGGTTGGCGCGGCTGCGCTGCGGCCCGGGGCAGTGCGGAGCCAGGACAGTCGCGGCGCTGACGCCCGCGGGCCCCAGCTGCAGATATGAAGCG*GAGCCGCTGCCGCGACCGACCGCAGCCGCCGCCGCCCGACCGCCGGGAGGATGGAGTTCAGCGGGCAGCGGAGCTGTCTCAGTCTTTGCCG*NNNNNNNNNNNNNNNNNNNNNNNNNNNNNNNNNNNNNNNNNNNNNNNNNNNNNNNNNNNNNNNNNNNNNNNNNNNNNNNNNNNNNNNNNNNNNNNNNNNNNNNNNNNNNNNNNNNNNNNNNNNNNNNNNNNNNNNNNNNNNNNNNNNNNNNNNNNNNNNNNNNNNNNNNNNNNNNNNNNNNNNNNNNNNNNNNNNNNNNNNNNNNNNNNNNNNNNNNNNNNNNNNNNNNNNNNNNNNNNNNNNNNNNNNNNNNNNNNNNNNNNNNNNNNNNNNNNNNNNNNNNNNNNNNNNNNNNNNNNNNNNNNNNNNNNNNNNNNNNNNNNNNNNNNNNNNNNNNNNNNNNNNNNNNNNNNNNNNNNNNNNNNNNNNNNNNNNNNNNNNNNNNNNNNNNNNNNNNNNNNNNNNNNNNNNNGGGCGGTGCAAGATGTGCTTTGTTAAACAGATGCTTGAAGGCAGCATGCTCGTTAAGAGTCATCACCAATCCCTAATCTCAAGTAATCAGGGACACAAACACTGCGGAAGGCCGCAGGGTCCTCTGCCTAGGAAAACCAGAGACCTTTGTTCACTTGTTTATCTGCTGACCTTCCCTCCACTATTGTCCCATGACCCTGCCAAATCCCCCTCTGTGAGAAACACCCAAGAATTATCAATAAAAAAATAAATTAAAAAAAAAAAAAAAAAAAAAAAAAAAAAAAAAAAAAAAAAAAAAAAAAAGAAAAAACTAACCTGCTCAAAAGATTTGTATACGA
